# Warm-loving species perform well under limiting resources: trait combinations for future climate

**DOI:** 10.1101/2025.07.04.662912

**Authors:** Sarah Levasseur, Vanessa Weber de Melo, Janneke Hille Ris Lambers, Christopher A. Klausmeier, Colin Kremer, Elena Litchman, Marta Reyes, Mridul K. Thomas, Anita Narwani

**Affiliations:** Department of Aquatic Ecology, Eawag, Dübendorf, Switzerland; Department of Environmental Systems Science, ETH Zürich, Zürich, Switzerland; Department of Plant Biology & Integrative Biology, W. K. Kellogg Biological Station, Michigan State University, Hickory Corners, MI, USA; Department of Ecology & Evolutionary Biology, University of Connecticut, Storrs, CT, USA; Department of Integrative Biology and W. K. Kellogg Biological Station, Michigan State University, Hickory Corners, MI, USA; Department F.-A. Forel for Environmental and Aquatic Sciences and Institute for Environmental Sciences, University of Geneva, Geneva, Switzerland

**Keywords:** growth rate, thermal optimum, thermal maximum, thermal minimum, thermal breadth, resources, temperature, performance, phytoplankton, *R\**

## Abstract

Ecosystems are warming alongside shifts in other abiotic factors, leading to interactive effects on populations and communities. This underscores the importance of studying how organisms respond to multiple environmental changes simultaneously. In aquatic ecosystems, as surface waters of lakes and oceans warm, longer and stronger periods of thermal stratification lead to changes in resource (light and nutrient) availability. We investigate the combined effect of temperature and resource availability on 19 populations (comprising 17 species) of freshwater phytoplankton to examine how temperature influences the minimum resource requirements (and Monod parameters) for light, nitrogen, and phosphorus. We also evaluate how resource availability affects each population’s thermal traits (i.e. thermal performance curve -TPC-parameters). When averaged across all populations, the requirements for light and phosphorus tended to display a U-shaped relationship along temperatures. Individual populations varied greatly in their responses to temperature, leading to shifts in the identity of the best competitor across the thermal gradient, particularly for nitrogen and phosphorus. TPC responses to resource limitation were highly variable, but thermal optima and maxima of individual populations often decreased with resource limitation, and thermal breadths (range where growth is 80% or more of its maximum) often increased due to a flattening of TPCs. Across all populations and resource types, the maximum optimum temperature across resource levels (maximum *Topt*) tended to be positively correlated with the temperature at which they had the lowest resource requirements (minimum *R\**). However, the temperature at which populations were the best competitors tended to be ∼5 °C colder on average than the temperature at which they grew the fastest. The populations with the highest thermal optima also had the lowest minimum resource requirements. Our findings reveal trait associations suggesting that some taxa already exhibit trait combinations that would support high performance under future warm and resource-limited conditions.

## Introduction

Anthropogenic climate change is increasing average annual temperatures in the majority of local ecosystems worldwide, including lakes (Calvin et al., 2023; Grant et al., 2021; Pörtner et al., 2021). Temperature effects on biochemical reaction rates, organismal metabolism, populations and communities are relatively well-documented, and supported by some mechanistic understanding (Allen et al., 2005; Bernacchi et al., 2001; Brown et al., 2004). However, while basic tenets of the impacts of temperature are understood, predicting the consequences of warming for populations, local community assembly and biodiversity remains a grand challenge in ecology, because warming is not happening in isolation from other aspects of environmental change.

Climate change is directly and indirectly altering the availability of limiting resources in many ecosystems. For example, in terrestrial systems, changes in patterns of precipitation with warming will lead to concurrent osmotic stress and altered patterns of resource availability to plants (Korell et al., 2021). In lakes, thermal stratification is becoming stronger and longer due to warming, often reducing the supply of deep-water nutrients to the surface and effectively increasing light availability (Kraemer et al., 2021b; Yankova et al., 2017). Additionally, independently of climate change, human activities are also influencing lake nutrient dynamics, with both intentional (Frenken et al., 2023; Hu & Huser, 2014) and unintentional (Higgins & Zanden, 2010; Perret-Gentil et al., 2024) nutrient reductions. In other cases, human activities are increasing nutrient loads as waters are warming. Nutrient loading can enhance productivity, potentially leading to a turbid state in which light availability is reduced (Ansari et al., 2010; Olea-Olea & Escolero, 2018). Accounting for these concurrent and potentially interactive effects of resource availability with warming may help to improve our understanding of the effects of climate change on population and community dynamics and improve the accuracy of predictions of the climate change impacts on biodiversity. Since climate change may be accompanied by changes in resource availability for many autotrophs, interactive effects of temperature and resource availability could have dramatic consequences for metabolism and growth (Allen & Gillooly, 2009; Huey & Kingsolver, 2019), and therefore also species distributions (Benedetti et al., 2021; Henson et al., 2021; Thomas et al., 2017). The importance of these interactions on processes across levels of ecological organization has already been noted (Cross et al., 2015), and a growing number of studies have investigated how species-level information relates to outcomes of community assembly under conditions of warming and resource limitation (Bestion et al., 2018b; Lewington-Pearce et al., 2019; Sunday et al., 2024). This was accompanied by a call for more multi-driver studies, particularly those investigating the combined impacts of warming and other axes of environmental change (Collins et al., 2022; Litchman & Thomas, 2023).

In this study, we investigate the combined effects of temperature and the availability of resources on phytoplankton population-level growth rates. Here, we follow up on this work by combining two theoretical frameworks that describe either the effects of resource availability (minimum resource requirement – R*) or temperature on population growth rates (thermal performance curves -TPC). These combined frameworks allow us to observe how temperature and resource limitation interact to influence populations, and how thermal and resource traits are related across populations of multiple species. To investigate the effects of resource limitation, we use the Monod equation and the Resource Competition Theory (Chase & Leibold, 2003; Tilman, 1982). The RCT proposes that an individual population’s growth rate as a function of resource availability can be used to characterize its minimum resource requirement for growth (*R\**, Fig. 1A). *R\** is the amount of resource needed by a species to accomplish a growth rate equal to its mortality. When competing, the population with the lowest *R\** for any given resource is predicted to win under constant conditions. Temperature may shift the identity of the species with the lowest *R\** for any given resource, altering the identity of the predicted winner of a competition (Fig. 1A). We can also investigate the impacts of temperature on the other parameters of the Monod equation, including the maximum specific growth rate (*µmax*, Fig. 1A), which represents the maximum growth rate attainable by a species in the absence of limitation. Separately, to investigate growth rates as a function of temperature, we draw on the concept of the thermal performance curve (TPC, Fig 1B). We investigate how multiple parameters describing thermal performances depend on resource availability (Fig. 1B): the optimum temperature (*Topt*), the upper thermal limit (*Tmax*), the lower limit (*Tmin*), and the thermal breadth (*Tbr* - temperature range where at least 80% of max growth rate is achieved).

**Figure 1.**
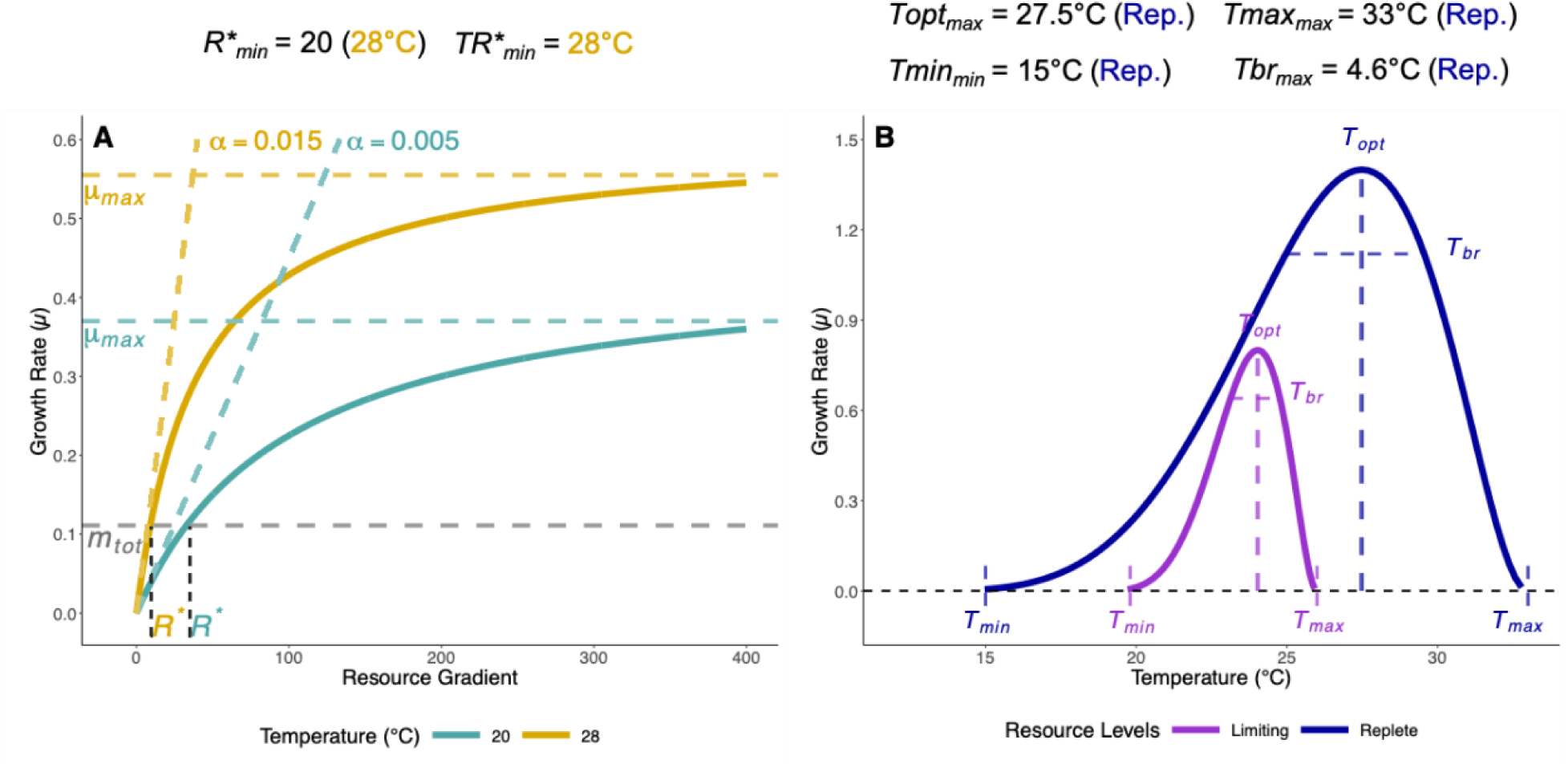
Conceptual figure of Monod curves, thermal performance curves (TPCs) and their investigated parameters. We investigated how temperature can impact Monod curve parameters (A), and how resource limitation can impact TPC parameters (B). In panel A, *R\** is the minimum resource requirement, and it reflects the resource concentration at which the population growth rate (*µ*) equals the total death/loss rate (*m_tot_*). The affinity *α* parameter of the Monod curve represents its initial slope (A). Secondary parameters were calculated for each species, such as the *R*_min_* which is the lowest resource requirement value across the temperature gradient and the *TR*_min_*, the temperature at which the *R*_min_* occurs. The thermal optimum is the temperature at which growth is maximized (*Topt*)(B). The thermal maximum (*Tmax*) is the upper temperature limit, where a population’s growth rate hits zero. The thermal (*Tmin*) minimum is the lower temperature limit. The thermal breadth (*Tbr*) represents the temperature range in which a species maintains at least 80% of its maximal growth rate at *Topt*. Additionally, for each species, we looked at the maximum values of the *Topt*, *Tmax* and *Tbr* across resource gradients (*Topt_max_*, *Tmax_max_*, *Tbr_max_*). Table S1 in the SI provides more information about all the variables introduced here.

Prior studies have investigated the temperature-dependence of Monod parameters or the resource dependence of TPC parameters. Most of these studies however, focused only on one or two species or resources (Bonilla et al., 2012; Corredor et al., 2021; Tilman et al., 1981). To our knowledge, only three studies have included up to six species, focused exclusively either on thermal (Bestion et al., 2018a) or resource parameters (or traits) (Bestion et al., 2018b; Lewington-Pearce et al., 2019). One recent study investigated how the temperature dependence of resource requirements and growth rate were related in four species, but the limited species pool prevented a more extensive analysis of trait associations (Sunday et al., 2024). Here, we investigated 19 unique freshwater phytoplankton populations (comprising 17 species, spanning three phyla), and how their specific growth rates vary in response to gradients of three different essential limiting resources (light, nitrogen, and phosphorus) crossed with temperature. By examining the temperature and resource-dependent growth parameters of an unprecedented number of species and resources within a single empirical study, we were able to assess the generality of these previous findings and ask new questions that could not be addressed by any individual prior study (see below).

Prior studies have provided valuable insights into how we may expect resource-dependent growth and competitive ability to vary with temperature. For example, resource requirements have been found to have a U-shaped relationship with temperature, declining from infinitely high values at *Tmin* and *Tmax* to a minimum *R\** value at intermediate temperatures (Lewington-Pearce et al., 2019; Sunday et al., 2024; Tilman et al., 1981), with species varying in the temperature at which they experience their lowest *R\**. We may also expect that the temperature at which species have their lowest *R*s* tends to be colder than the temperatures at which they grow fastest under replete conditions (*Topt*) (Sunday et al., 2024; Thomas et al., 2012). We may therefore expect that the impacts of warming will be dependent on whether species are growing under resource-replete or resource-limited conditions (Bestion et al., 2018b; Lewington-Pearce et al., 2019; Thomas et al., 2017). Previous work has also informed our expectations for how thermal traits may vary with resource availability. Specifically, we expect that thermal optima (*Topt*) and thermal maxima (*Tmax*) will tend to decline as resources become limiting (Bestion et al., 2018c b; Litchman & Thomas, 2023; Thomas et al., 2017), and this overall reduced thermal tolerance will shrink the thermal breadths of populations.

The broader taxonomic and environmental scope of our study with respect to previous work has allowed us to address the following, sometimes novel, questions and hypotheses:

1. How do the parameters of the Monod equation and *R*\* vary with temperature? Is there a general pattern to how *R\**s vary with temperature, or is there large variation in the shape and strength of these relationships across species and resources? H1) *R\** is generally U-shaped across the temperature gradient for all species and resources. H2) *µ_max_* is hump-shaped across the temperature gradient for all species and resources. If there is substantial variation among species, are the differences in temperature-resource interactions related to higher taxonomic grouping e.g. phyla? (No *a priori* hypotheses)
2. How do the parameters and features of TPCs vary with the availability of resources? H3) *Topt* and *Tmax* decline with resource limitation H4) *Tbr* declines with resource limitation Does this relationship vary with the identity of the limiting resource? (No *a priori* hypotheses)
3. How are the parameters of resource and temperature-dependent growth related to one another across species and resources? What do the trait covariance patterns imply for how communities will respond to environmental change? Are the differences among species structured in a way that suggests trade-offs constraining species performance, such that some species are better adapted to one stress at the cost of adaptation to another (low nutrients vs. high temperatures)? (No *a priori* hypotheses)

Finally, we investigate how trait averages in our population pool vary across thermal and resource gradients, giving an indication of expected trait values for these environments across a population or species pool.

## Methods

We isolated twenty phytoplankton populations from four Swiss lakes - Greifen, Zurich, Constance and Lucerne - between August and November 2021. We selected these lakes because they span a range of trophic states, from eutrophic to oligotrophic (Table S2). We used a combination of techniques to isolate each population from single colonies or cells. These techniques included dilution, colony-picking under a microscope and Fluorescence-Activated Cell Sorting (FACS) (Andersen, 2005). Isolates were identified morphologically to the species level whenever possible. In addition, we performed DNA barcode sequencing of multiple genetic markers to verify their identity (see Supplemental Methods for details). These approaches helped us confirm the species identity of our focal species. This process gave us a pool of 20 uni-algal isolates (“populations” hereafter), with one having been purchased from a culture collection (*Synechococcus* sp., Table S3). Our cultures were not strictly axenic, but bacteria content was kept low as we grew our cultures in a mineral medium without an organic carbon source. In addition, each experimental step was started from low density with sterile media, hence, there was no dead algal biomass for the bacteria to grow on. Of these 20 populations, 18 were different species and two were different isolates of the same species (two *Nitzschia palea* and two *Microcystis aeruginosa*, Table S3). The initial isolate pool was eventually reduced to 19 (17 species) because the filamentous species *Planktothrix agardhii* performed poorly in all experiments. Once isolated, we maintained all populations under standard conditions of 15°C and replete nutrients using COMBO medium (Kilham et al., 1998), modified nitrogen concentration -800 µmol·L^-1^ of NaNO_3_- and silica -350 µmol·L^-1^ of Na_2_SiO_3_·9H_2_O). We cultured the phytoplankton under a 14h:10h light:dark cycle at 20 μmol photons m^-2^·s^-1^ irradiance.

We investigated the effects of temperature and resource availability on growth rates for three different resources: nitrogen, phosphorus and light. Each resource type, with eight resource levels (Table 1), was crossed by six temperatures (15, 20, 24, 28, 32, 35 °C, see supplemental method for more). This resulted in 48 unique treatments for each of the three resources, run separately on each population (19) and replicated 4 times, comprising a total of 10,944 independent growth rate estimates.

**Table 1.**
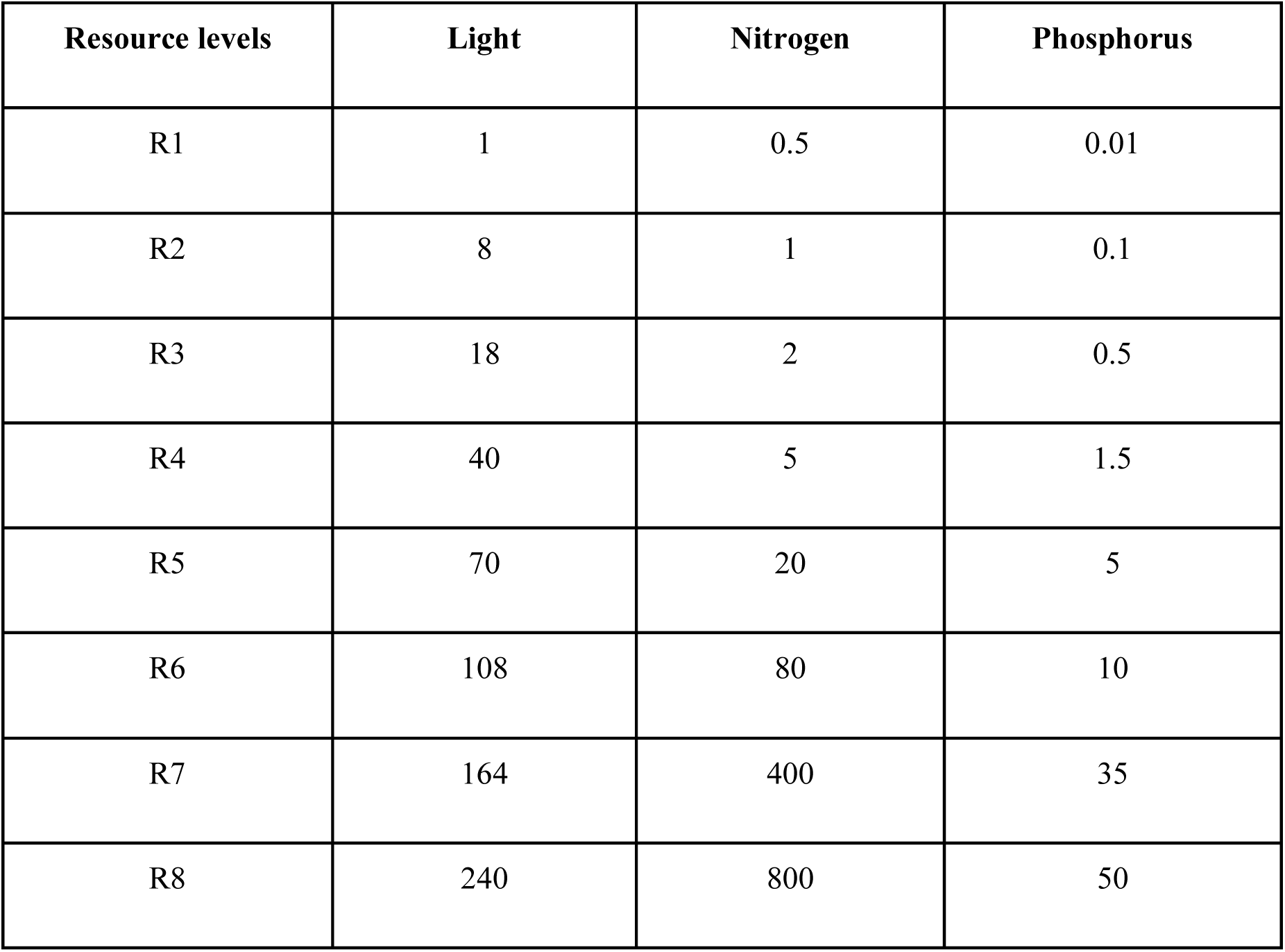
Resource levels for the resource limitation experiments. The nitrogen source was NaNO_3_, the phosphorus source was K_2_HPO_4_, and the light represented levels of photosynthetically active radiation (PAR). Light intensities are in units of μmol photons m^-2^·s^-1^ and nutrient concentrations are in µmol⋅ L^-1^. When they were not the manipulated resources, the light was 110 μmol photons m^-2^·s^-1^, nitrogen was 800 µmol⋅L^-1^ and phosphorus was 50 µmol⋅ L^-1^. All experimental treatments were silica enriched (350 µmol⋅L^-1^ of Na_2_SiO_3_⋅9H_2_O).

For all three resources, before the start of growth rate experiments, populations underwent a period of acclimation (Fig. S1). Cultures were first acclimated to given temperatures, with one new batch culture for each of the six temperatures. In a second step, temperature-acclimated batch cultures were each acclimated to one of two resource levels representing either the lowest or mid-point of the resource gradient for each resource. Acclimation was necessary to eliminate potential carry-over of nutrients from the initial batch culture and to prepare the cultures for the growth experiments i.e. to minimize lag periods caused by physiological plasticity or maternal inheritance. All details and timings of these procedures are available in the Supplemental Methods.

### Temperature x resource growth rate experiments

The growth experiments were run in 96-well plates (Cellstar, Greiner Bio-One ®), with each well filled with 200 µL, using a Freedom EVO100 TECAN automated liquid handler. Each plate comprised one resource level and temperature and contained the 19 populations with their 4 replicates. The position of each algal population within the plate was randomized, while the first and last columns on the plate contained sterile Nanopure^TM^ water to prevent evaporation of the wells, especially for the 35°C treatment. Once the plates were inoculated with the phytoplankton, they were sealed with permeable Breathe-Easy® membranes, allowing for gas exchange and preventing cross-contamination. After the acclimation, to ensure low starting densities of all acclimated cultures at the start of the growth experiment, we measured pigment fluorescence of all acclimated cultures. We used an Agilent Cytation 5 plate reader to measure pigment fluorescence as a proxy for algal biomass. We performed dilutions to achieve a target inoculation level of 20 relative fluorescence units (RFU) for the start of the growth experiment. For the Chlorophyta and Bacillariophyta, we estimated chlorophyll fluorescence (excitation wavelength was 445 nm and emission was 685 nm), while for the Cyanobacteria we measured phycocyanin fluorescence (excitation wavelength was 586 nm and emission was 647 nm).

We measured population growth rates by estimating rates of change of RFU over time. We made the first RFU measurements immediately after inoculation. Fluorescence was measured for all plates twice daily for the first four days of the experiment, with approximately 9h between readings. After day 4, we continued monitoring once per day, for a minimum of 5 to 7 days, until the cultures reached a steady state, or we had enough data in the exponential phase to estimate a growth rate.

Populations grew in Infors™ incubators and were shaken at 50 rpm with a 18h:6h light:dark cycle. The media were enriched with silica with a concentration of 350 µmol·L^-1^ of Na_2_SiO_3_·9H_2_O to ensure that the diatoms did not experience silica limitation. Nitrogen and phosphorus concentrations, unless manipulated for a certain experiment (see Table 1) were 800 µmol·L^-1^ and 50 µmol·L^-1^ respectively. Except for the light experiment (see Table 1), the light level was at 110 μmol photons·m^-2^·s^-1^.

Each resource experiment generated 3,648 time series: 19 populations x 6 temperatures x 8 resource levels x 4 replicates). All data analyses in the study were performed using R and RStudio (Versions R 4.3.1 and 2023.12.1+402). Each time series was used to estimate a population growth rate using the R package “growthTools”, version 0.1.2 (Kremer, 2020), see Supplemental Methods for details). Scripts that enable data processing from relative fluorescence units to curated population growth rates are included in our online information (*citation for open data repository will be provided upon acceptance*).

### Temperature dependence of Monod parameters

We estimated the parameters of the Monod equation for each population and temperature combination by fitting the population growth rate (*µ*) of each experimental population against the resource levels *R* using Bayesian estimation (brms package, version 2.22.0, see (*Meddle-Scor149*, n.d.) vignettes #5 & #6 for Bayesian fitting tutorials):

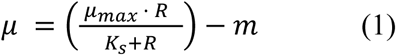

where *µ_max_* is the gross maximum specific growth rate, *Ks* is the half-saturation constant for growth, and *m* is the intrinsic loss rate. The mean and standard deviation of the priors for *µ_max_* (1, 0.9) and *m* (0, 0.7) used for the analysis were the same for each of the three resources. The priors for *Ks* were resource-specific, for light (2.5, 0.7), for nitrogen (2.5, 0.9) and for phosphorus (0.001, 0.8). We constrained all the parameters to be positive with a lower bound at 0. See Fig. S2 and Table S4 for a summary of the prior’s distribution. To estimate the posterior distribution of the Monod parameters, we ran 10,000 iterations in each of the 4 chains, then retained the last 5,000 from each chain (total of 20,000 post burn draws). From the posteriors, we then extracted each parameter’s median value and 95% credible interval (CI). Median parameter values and CI were also obtained for each population, one per resource by temperature treatment. These estimates were used to generate Monod models of each population’s growth rate across each resource gradient, with one curve per temperature (Fig. S4-S6).

We calculated three derived traits from the posteriors for each retained set of Monod parameters. The affinity *α* represents the initial slope of the Monod curve, defined as:

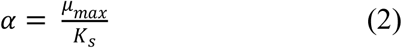

In the remainder of the paper, we focus on *α* and not *K_s_* because it reflects the slope of the initial growth versus resource curve in the Monod equation, decoupled from the value of *µ_max_*, unlike *K_s_* (Aksnes & Egge, 1991). For each fit of the Monod equation, and for each population, temperature, and resource type, we also calculated the minimum resource requirement, *R\**, or *I\** for light, *N\** for nitrogen and *P\** for phosphorus, as:

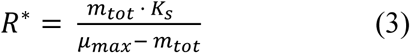

where *m_tot_* _=_ *m_ext_* _+_ *m* is the sum of the intrinsic loss rate -also defined as the species-specific density independent mortality- *m*, and the extrinsic loss rate, *m_ext_*, which is externally determined by the ecosystem (in a chemostat, it’s the dilution rate; in lakes, the sinking rate). The true value of *m_ext_* is a constant that is system-specific; here, for simplicity, we set it at a value of 0.1. *R\** is the amount of a resource a species needs to have a population growth rate (*µ;* which accounts for *m*) equal to the additional external loss rate (*m_ext_*) and serves as a measure of equilibrium competitive ability (lower is better). Any resource level above that threshold enables positive population growth (*µ* > *m_ext_*).

We correct the maximum specific population growth rate, *µ_max_*, for our estimated intrinsic loss rate, *m*. When *µ* < 0 at zero resource, *m* is positive and describes the intrinsic rate of loss at zero resource. The estimate of *µ_max_* does not account for this loss rate, and so *µ_max_* must be lowered by this amount (*net-µ_max_*, equation 4).

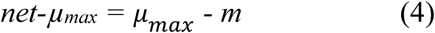

We performed the next steps of the analysis only for populations for which each temperature by resource type had at least three resource levels at which *µ* was above *m_ext_*.

### Resource dependence of thermal performance curve parameters

We fitted thermal performance curves (TPCs) for each population and resource level. There are many ways to mathematically characterize the shape and parameters of the TPC (Low-Décarie et al., 2017; Padfield et al., 2021), however not all are equally amenable to being combined with a resource-dependent growth equation. For this reason, we focus on the double exponential model (Thomas et al., 2017), which has previously been demonstrated to have biologically realistic and mathematically meaningful behavior when integrated with a Monod model. We used the same Bayesian estimation tools as for the Monod parameters to fit a re-arranged version of the double exponential model of temperature, developed in (Kremer et al., 2024):

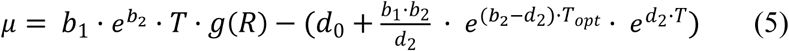

where *b_1_* is the birth rate at 0°C, g(R) is a resource-dependent term to reflect birth as a function of resource being either limited or replete, *b_2_* is the exponential change in birth rate as temperature increases, *T* is the environmental temperature, *d_0_* is the temperature-independent loss rate, *d_2_* is the exponential change in death rate as temperature increases, *Topt* is the temperature where growth rate is maximized if resource is not limiting. Our modification of the equation 5 - described in Table S5- is an algebraic rearrangement that facilitates model-fitting and improves the identifiability of parameters. This generated a TPC representing the growth rates of each population across the temperature gradient, with one curve for each level of each resource (Fig. S7-S9). For each fit, we further analyzed TPC parameters only when *µ* was above *m_ext_* for at least three temperatures. The mean and standard deviation of the prior distributions for the parameters of the modified equation 5 were the same for each of the three resources (Table S6, Fig. S3). The retained parameters from the 5,000 post-burn iterations were used to calculate three derived parameters: *Tmax*, *Tmin* and *Tbr*. *Tmax* represents the upper temperature limit for population growth rate at which the TPC crosses zero, and *Tmin* represents the lower limit. *Tbr* represents the thermal breadth, the temperature range where growth is equal or greater than 80% of the maximal growth rate at *Topt*. We used the package rootSolve (version 1.8.2.4) to estimate these derived parameters and obtained their median values and CI for each population and resource level (per resource type).

We excluded parameter estimates for which the posterior median fell outside our measured range of temperatures and resources from further analysis. Specifically, we excluded all *Topt* estimates that were below our minimum of 15 or above our maximum of 35°C. This removed 22 out of the 344 total estimates, where values of the removed estimates ranged from 4°C to 14°C and 37 to 71°C respectively. We removed *Tmax* estimates that were more than 5°C above the highest measured temperature (22 estimates removed ranging from 41 to 65°C). We excluded *Tmin* values that were more than 15°C below our lowest temperature, which eliminated all values under 0°C (254 estimates from -0.03 to -85°C). Only 98 estimated values of *Tmin* remained after this selection (Fig. S10G-I), for this reason, we eliminated *Tmin* from further analysis. Finally, we excluded *Tbr* estimates larger than our experimental temperature range of 20°C (eliminated 20 values from 20 to 47°C).

All parameter estimates and credible intervals are available in the SI.

### Fitting of generalized additive models and categorizations of their shapes

To describe how the Monod parameters vary along temperature gradients (Fig. 2) and how the TPC parameters vary along resource gradients (Fig. 3), we used Generalized Additive Models (GAMs; “mgcv” R package version 1.9-1), or linear models (LM) if there were no more than three experimental temperature or resource levels, respectively. We performed this analysis for each population to investigate how their parameters varied across the gradients. To incorporate our confidence in parameter estimates into the fitting of the models, we used a posterior bootstrapping procedure as follows: 1) We randomly subsampled 5,000 draws from the 20,000 in the posterior, for each of the levels across the gradients. 2) We fit 5,000 GAMs, one to each set of single random draws of posterior parameter estimates across the gradient. 3) We calculated the mean predicted values and 95% credible intervals from these 5,000 fits. 4) When 80% or more of the 5,000 fits showed signs of linearity (see EDF ≤ 1.6, see below), we re-fitted linear models to all 5,000 sub-sampled posteriors and estimated their slopes. We also estimated the mean predicted linear model and its 95% CI.

**Figure 2.**
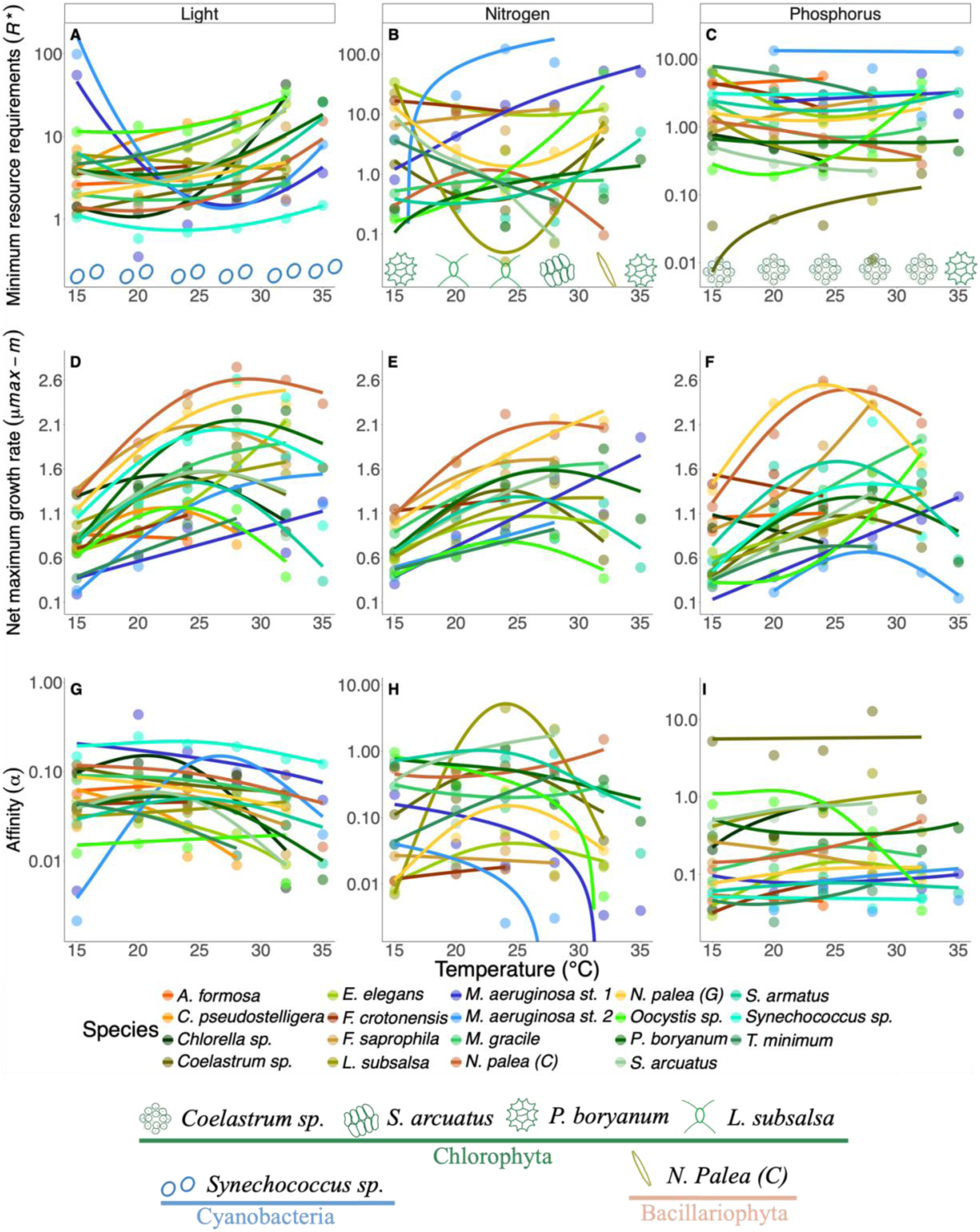
Resource-dependent growth parameters plotted as a function of temperature for each experiment (light, nitrogen and phosphorus from left to right). For each parameter, points represent the median estimated value from the Bayesian posterior distribution: minimum resource requirements (*R\**) (A-C), net maximum growth rates (*µ_max_ - m*) (D-F) and affinities (*α*) (G-I). The units for *R\** of light are μmol photons m^-2^·s^-1^ (A), while the units of *R\** for nitrogen and phosphorus (B&C) are µmol⋅ L^-1^. Phytoplankton icons across the bottom of each *R\** panel represent the population that is predicted the best competitor for that resource and temperature. The *R\** estimates incorporated an additional temperature-independent mortality term (*m_ext_* = 0.1). The middle row (D-F) shows *µ_max_ - m* (units, d^-1^) as a function of temperature. In the bottom row (G-I) points are the affinity (*α*, units are growth rate (d^-1^) per unit resource concentration). The y-axes in the *R\** panels (A-C) for affinity (G-I) are log-transformed for easier visualisation. Due to this transformation, linear models display a curvature. Bacillariophyta are indicated by hues in the yellow-brown spectrum, Chlorophyta are indicated by shades of green, and Cyanobacteria are indicated by shades of blue and turquoise.

**Figure 3.**
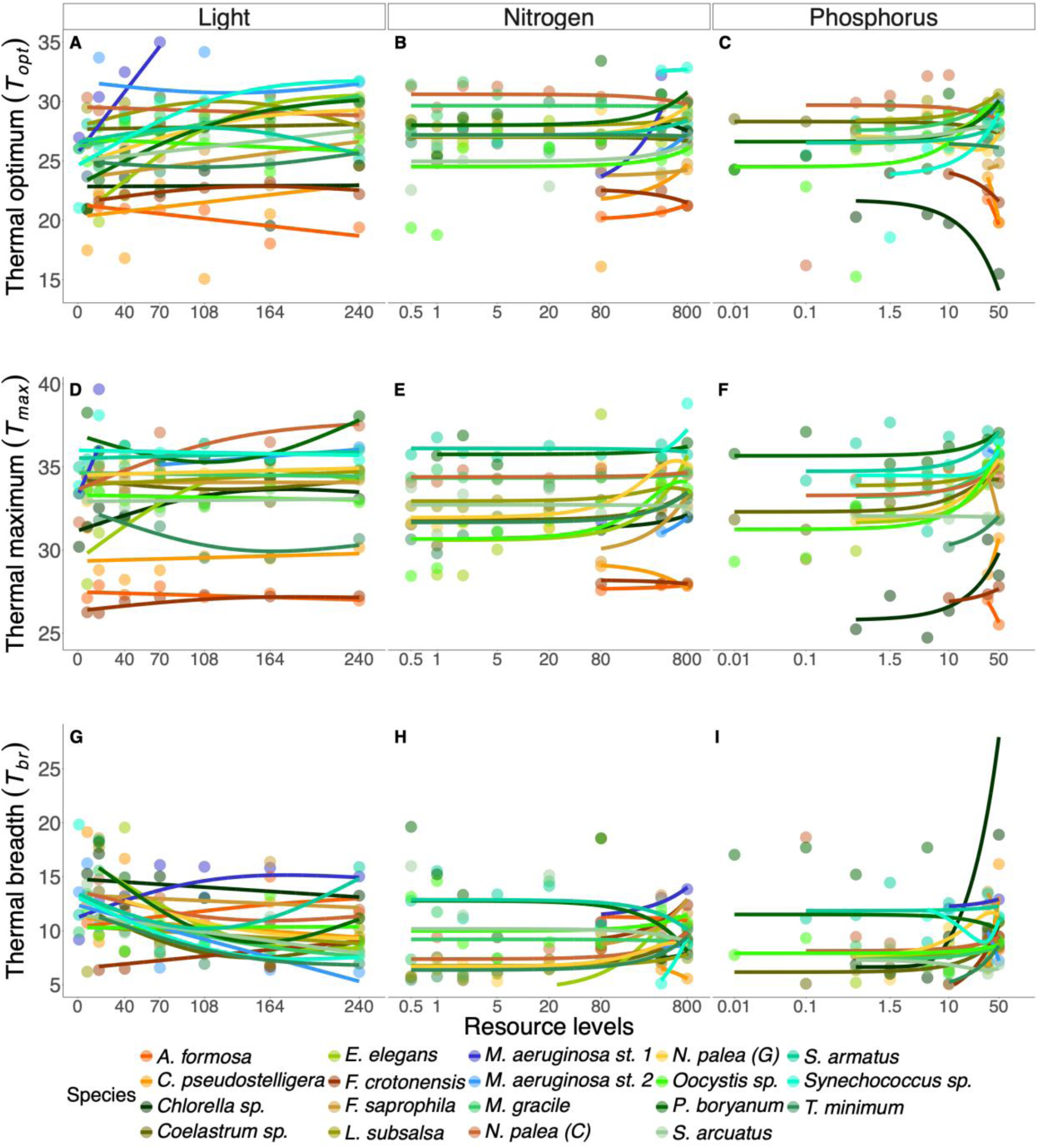
Thermal performance curve parameters as a function of resource availability for each experiment (light, nitrogen and phosphorus from left to right). For each parameter, points represent the median value of the Bayesian posterior distribution: panels in the top row (A-C) represent the thermal optimum (*Topt*); the second row (D-F) the thermal maximum (*Tmax*); the third row (G-I) the thermal breadths (*Tbr*). Temperatures are reported in °C. For the nitrogen and phosphorus panels (B&C, E&F, and H&I), the resource axes were logged, and models were fit to log-transformed values. For all panels, only significant curves are accompanied by shaded standard error ribbons. M. aeruginosa strain 1 is absent from panels for *Topt* and *Tmax* in the phosphorus experiment (C&F) (see Methods). Phytoplankton groups are shaded as in Figure 2.

To characterize the shapes of the relationships of parameters to environmental gradients, we used the estimated degrees of freedom (EDF) of the GAMs as a measure of nonlinearity. EDF values close to 1 indicate nearly linear relationships, while larger values (e.g. ≥ 2) suggest more strongly nonlinear relationships (Hunsicker et al., 2016; Zuur, 2009). To reduce the risk of overfitting (detecting a nonlinear relationship not strongly supported by the data), we set an EDF threshold of ≤ 1.6 to characterize relationships indistinguishable from a linear trend (Hunsicker et al., 2016; Zuur, 2009). For this reason, curves with an EDF ≤ 1.6, were re-fitted with linear models (golden curves in Fig. S11-14, S15-18, and increasing or decreasing linear model parameters in Tables S7-S10). EDF values > 1.6 indicative of more nonlinear, sometimes non-monotonic relationships within the observed range of the predictor, retained their GAM fits, and we further assessed their shapes. For these curves, to detect minima or maxima, we analyzed the first derivative of the GAM smoother. We checked whether the derivatives crossed zero within the observed range of the predictor, indicating a minimum or maximum (“critical points” in Tables S7and S9). When no critical point was observed, we classified the curves as monotonic increasing or decreasing. When a minimum value was identified (derivative was negative and crossed zero to become positive), the relationships were classified as U-shaped, while the presence of a maximum value indicated a hump-shaped relationship (derivative was positive and crossed zero to become negative). After the determination of the shape for a trait across a gradient (temperature or resource, see Tables S7 and S9), we designated a particular population as having a shape when ≥ 51% of the 5,000 fits had the same shape. When no shape met the criteria, we characterized the it as “uncertain”. GAMs for all thermal parameters for the nitrogen and phosphorus experiments were fit to log_10_-transformed resource levels. This improved data visualization and GAM fits due the preponderance of very low-nutrient treatments that were necessary to estimate *R\**s for these two resources (Table 1). As a result of these transformations however, we refer to linear relationships from Table S7 as “monotonic” in our Results section. We make this distinction because these linear relationships on the logged resource levels are monotonic but nonlinear, asymptotic relationships on the untransformed resource levels.

Finally, to generate an expectation of how parameters vary on average across environmental gradients (regardless of population), we fitted unweighted GAMs to the mean (of the median) parameters estimates across all populations (“Mean spp”, Tables S8 and S10). We then applied the same EDF and shape criteria as above, refitting with linear models for EDF ≤ 1.6.

All GAMs were fitted with gaussian error distributions, except for *R\** and *α* across temperatures (Fig. 2A-C & G-I), for which we used gamma distributions instead, as these parameters are bounded by zero.

### Ordination of Monod and TPC parameters

We performed a principal components analysis (PCA) to explore how the parameters of a population’s Monod and TPC curves were associated. These parameters were estimated under multiple environmental conditions, and to enable comparison, we calculated for each population and resource type, six summary statistics of this variation: 1) The lowest minimum resource requirement across all temperatures. We log-transformed and then standardized these values (i.e. we subtracted mean and divided by standard deviation) to obtain the *scaled logR*_min_*. The scaling accounts for the different units and absolute values of the requirements of the different resources. 2) The experimental temperature at which this *scaled logR*_min_* occurred (*TR*_min_*); 3) The largest net maximum growth rates (net-*µmax_max_*) across all temperatures; 4) the highest thermal optimum (*Topt_max_*) across all resource levels; 5) the highest thermal maximum (*Tmax_max_*) across all resource levels; 6) the widest thermal breadth (*Tbr_max_*) across all resource levels. We first performed the PCA on the data from all three resource experiments (Fig. 4A&B), and subsequently on each resource separately (Fig. S20). We used the packages MASS (version 7.3-61) and factoextra (version 1.0.7) for these analyses. We clustered PCA loadings by population’s phyla, as well as by experiment and tested for significant differences among clusters using an Analyses of Similarity (ANOSIM) test from the vegan package (version 2.6-4) in R. We tested for differences in *Topt_max_* and *R*_min_* among phyla using a Welch’s ANOVA and Games-Howell post-hoc test respectively (Fig. 4C&D). We explored all bivariate correlations among the summary variables using a correlation matrix of Kendall’s rank correlations (Fig. S19), we have selected a few of the significant correlations to explore further how traits relate to one another (Fig. 5).

**Figure 4.**
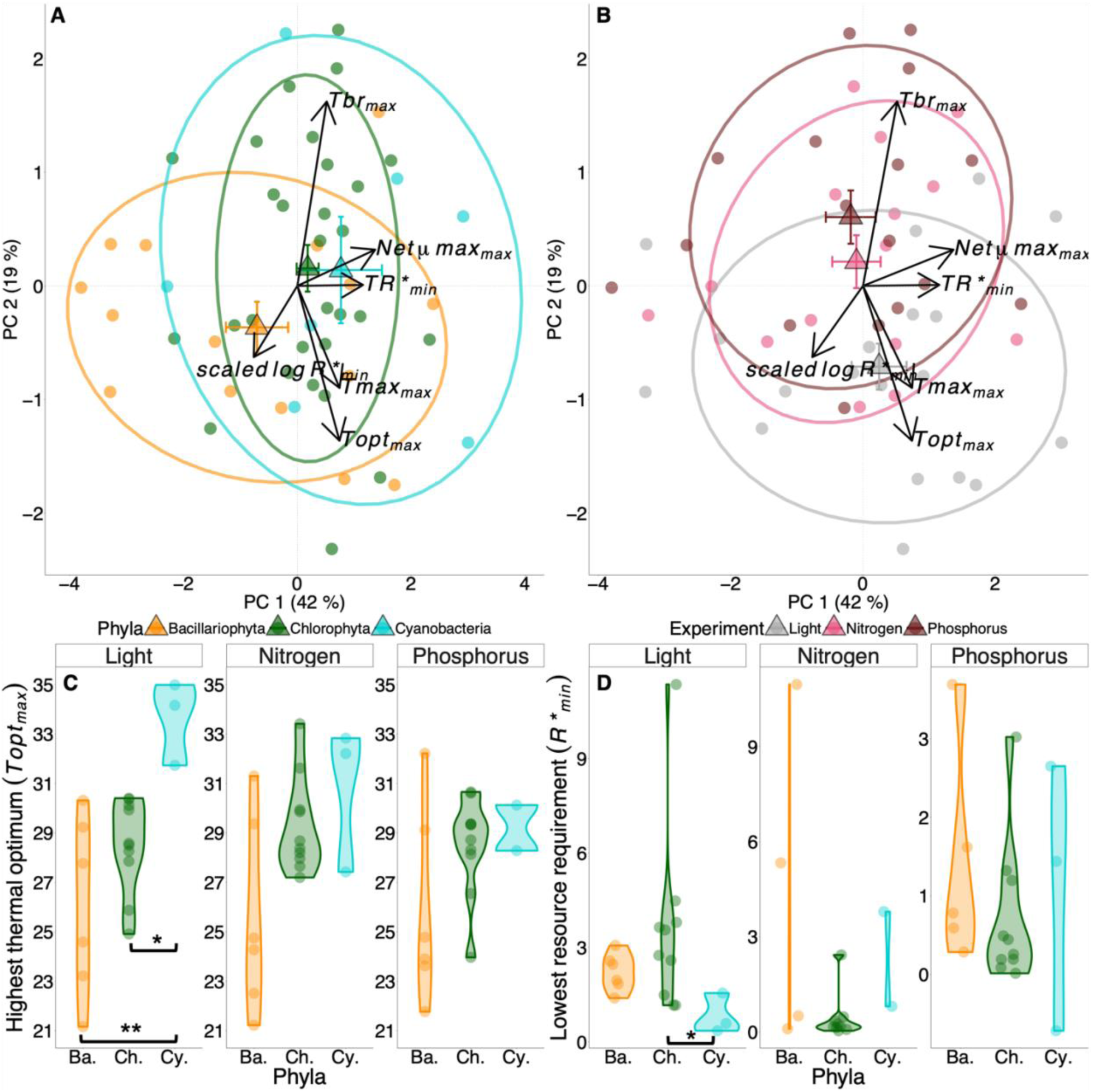
Principal components analysis (PCA, A & B) and boxplots (C & D) of the summary variables (minimum and maximum) of various Monod and TPC parameters for all three resources. The summary variables are calculated across temperature levels for each resource experiment and across resource levels for TPCs fitted for each resource. Parameters included in the PCAs are: the scaled log-transformed lowest minimum resource requirement across temperature for each resource (*scaled logR*_min_*), the temperature at which this minimum *R\** occurred (*TR*_min_*), the greatest maximum growth rate (*µmax_max_*), the greatest thermal optimum (*Topt_max_*), the greatest maximum temperature (*Tmax_max_*), and the maximum thermal breadth (*Tbr_max_*). Centroids are indicated with large triangular symbols (+/- standard error bars for PC1 and PC2) and ellipses represent one standard error of the means of the centroids (68% confidence ellipses). Box plots (C & D) show variation in the highest thermal optimum (°C), *Topt_max_* (C) and lowest minimum resource requirement (μmol photons m-2·s-1 for light, or µmol⋅L-1for nitrogen or phosphorus), *R*_min_* (D) with facets for each resource and separated by phylum. Significant differences in panels C and D indicate at least p < 0.05 (*) or < 0.01 (**) based on Welch’s ANOVA and the Games-Howell post hoc test.

**Figure 5.**
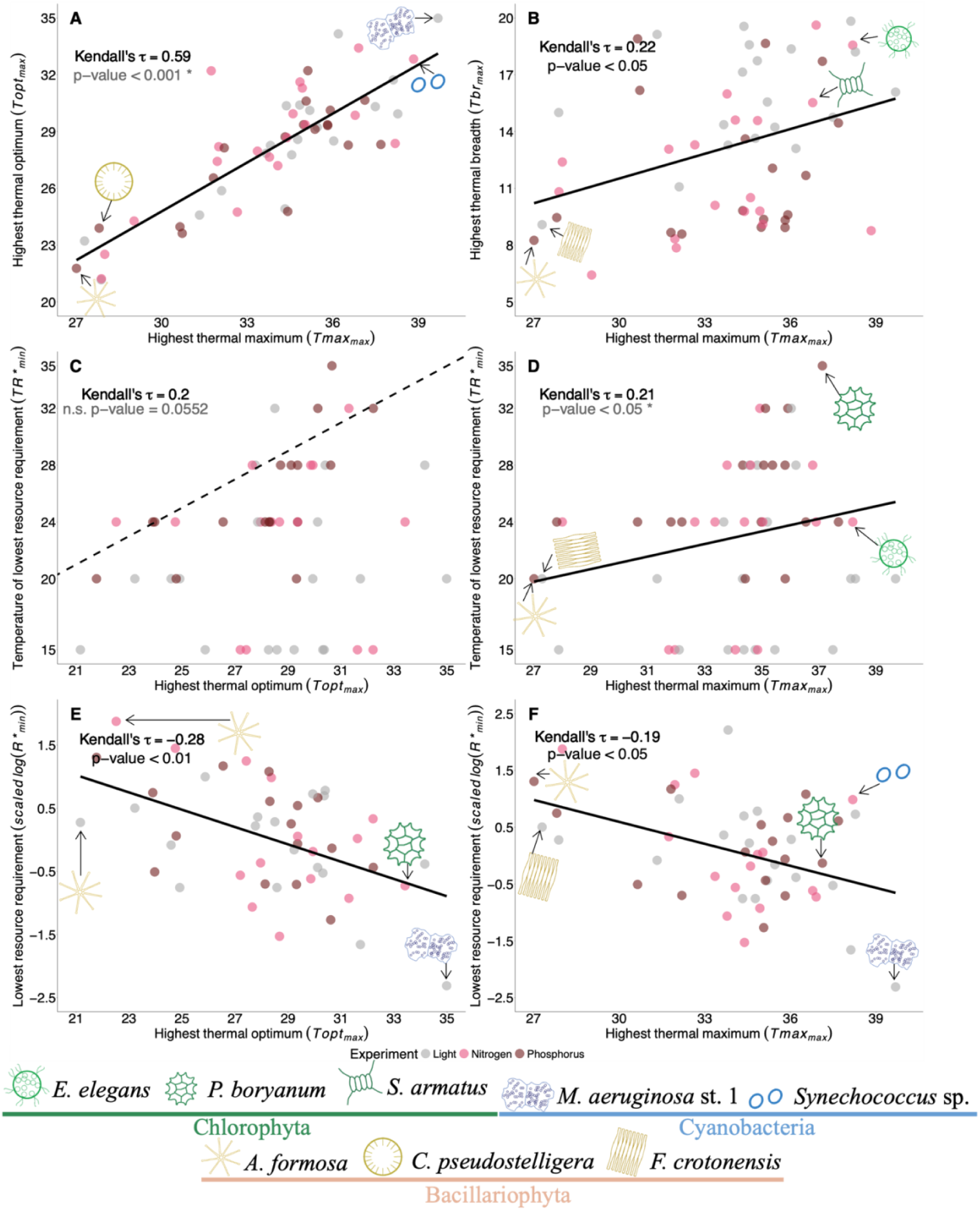
Selected significant correlation plots of summary variables (minimum or maximum) of Monod and TPC parameters from all three experiments (all plots are shown in Fig. S17). Kendall’s rank correlations are presented to assess the direction and strength of the relationship. The solid line in each panel represents a linear model. The resource identity (light, nitrogen or phosphorus) never had a significant interaction with the predictor in the model (p > 0.05), and for this reason, we provide the regression line and correlation statistics for all three resource experiments together. Panel A presents the maximum thermal optimum (*Topt_max_*) plotted against the maximum thermal maximum (*Tmax_max_*); panel B is the highest thermal breadth value (*Tbr_max_*) plotted against *Tmax_max_*; panel C is the temperature at which the minimum *R\** occurred (*TR*_min_*) plotted against *Topt_max_*; panel D is *TR*_min_* plotted against *Tmax_max_*; panel E is the lowest minimum resource requirement scaled and log (*scaled logR*_min_*) as a function of *Topt_max_*; and panel F is *scaled logR*_min_* as a function of *Tmax_max_*. In panel C, the black dashed line is a 1:1 line. In panels A and D, the rank correlation and p-value are accompanied by a “*” to highlight that these traits may be expected to have a positive relationship based on random sampling. Nevertheless, the p-value reflects a significance that is different from a slope of zero (for more see Fig S21 & 22). For panels E and F, the y-axis is scaled to account for the different units and absolute values of the requirements of the different resources. In each panel, cartoons and labels shown at particular extremes of the plot indicate which species had these trait combinations. Sample sizes for panels are: A&B: 56, C-F: 52.

## Results

### Temperature dependence of resource traits

We observed notable variation in *R\**s ranging across several orders of magnitude for all three resources: for light 0.35-98 μmol photons·m-2·s-1; for nitrogen 0.03-122.7 µmol⋅L-1 and for phosphorus 0.01-13.2 µmol⋅L-1. The data indicate that the same population remains the most competitive for light across the temperature gradient, as well for phosphorus (from 15 to 32°C). Specifically, the Cyanobacterium *Synechococcus* sp. maintained the lowest *I\** at all temperatures that we investigated, (Fig. 2A). For phosphorus, the Chlorophyta *Coelastrum* sp. had the lowest *P\** across the temperature gradient (Fig. 2C), except at 35°C (see Fig. S6) where it’s *P. boryanum*. For nitrogen, the identity of the predicted most competitive population changes with temperature (Fig. 2B), suggesting that up to four different populations would be competitively superior for nitrogen, depending on the temperature. Chlorophyta tended to be the predicted best competitors overall for nitrogen, with only one Bacillariophyta, *N. palea (C)* predicted as the winner at 32°C (Fig. 2B).

For light and phosphorus, the average *R\** across all populations in our pool displayed a U-shaped relationship with temperature (Fig. S11A&C, S12 A&C, Table S8), although non-significant. The average *N\** across all populations increased linearly (significantly) along the temperature gradient (Fig. S11B, S12 B, Table S8), indicating that warmer environments will on average favor populations with higher nitrogen requirements. The relationship of individual populations’ *R\**s with our temperature gradient varied in shape across species and resource types. For light, the majority of populations (11 of 19) showed evidence of U-shaped responses to temperature across the observed temperatures. Seven populations had *I*s* demonstrating monotonic or linear increase and one population was a monotonic decrease (Fig. S11A, S12A, Table S7). Populations displayed the most variation in their shape and direction of responses for *N\**, out of 15 populations, five showed U-shapes, three hump-shaped relationship, five linear or monotonic increase and only two with linear or monotonic decrease (Fig. S11B, S12B, Table S7). Finally, for phosphorus, out of 18, 55% had U-shape or monotonic decrease, and four had linearly decreasing, and four had linearly increasing *P\**s. Note that relationships for *R\** and affinity against temperature, that we have deemed linear on their original scales (Fig. S12&14) appear nonlinear but monotonic in Fig. 2 (A-C &G-I) due to the log_10_-transformation of the y-axis.

Individual population’s net maximum population growth rates (*net-µ_max_*) for all three resources varied extensively with temperature, from a minimum value of 0.14 (day^-1^) at 15°C and a maximum value of 2.74 (day^-1^) at 28°C, across all populations and resources. When averaged across all populations, *net-µ_max_* displayed a hump-shaped relationship with temperature (significant for light and nitrogen, Fig. S11D&E, Table S8). When considering across resources, nearly 50% of individual populations displayed a unimodal relationship of the *net-µ_max_* (Fig. 2D-F, Fig. S13, Table S7). The next two prevalent patterns were monotonic (25%) and linear increase (20%) (Fig. S13, Table S7), indicating that their thermal optima occurred at temperatures higher than we tested in our experiments (e.g. *M. aeruginosa* strain 1, Fig. 2D-F). In a minority of cases (< 6%), *net-µ_max_* decreased with increasing temperature (light once, phosphorus twice -Fig. S13, Table S7).

The affinity parameter (*α*) varied over 2 orders of magnitude for light and phosphorus, and almost 4 orders of magnitude for nitrogen (Fig. 2G-I). When averaged across all populations, for light and nitrogen, *α* tended to vary according to hump-shaped relationship, while for phosphorus it displayed a linearly decreasing trend (Fig. S11G-I, S14, Table S8). Individually, populations varied greatly in the way in which affinities for all three resources responded to temperature. Overall, there was a split of 30% with hump-shape, 35% with monotonic or linear increases, 31% displaying linear or monotonic decreases, and a rare U-shape (4%)(Fig. S14, Table S7).

### Resource dependence of thermal traits

*Topt* varied between 15℃ and 35℃ across the gradients of all three resources (Fig. 3A-C). When averaged across all populations, *Topt* showed a significant linear increase with light availability (Fig. S15A-C, Table S10) but a linear decrease for nitrogen and a hump-shape for phosphorus (both non-significant). *Topt*s of individual populations increased with light -linearly or monotonically- for 57.9% of the populations, and for another three populations, *Topt* had a hump-shaped relationship (Fig. S16A, Table S9. Such a hump-shaped relationship has previously been observed by (Edwards et al., 2016), and is supported by theory (Kremer et al., 2024). More than 60% of the populations showed strong support for linear increases in *Topt* with nitrogen and the remaining populations showed support for linear decreases in *Topt* over our nitrogen levels. For phosphorus, six populations showed decreasing *Topt* with increasing phosphorus availability, while 10 populations showed increasing *Topts* (7 linear, 3 monotonic, Fig. S16B&C, Table S9). Across all three experiments, there were four cases in which *Topt* varied in a hump-shape and one U-shape (Fig. S16, Table S9).

*Tmax* estimates varied between a minimum of 24℃ and 40℃ across the populations and resource levels. There was a significant linear increase between *Tmax* and phosphorus when averaged across all populations (Fig. S15D-F, Table S10). Individually, over 60% of all populations (across resource types) displayed monotonically or linearly increasing *Tmax* values (Fig. S17, Table S9). Three populations displayed *Tmax* varied in a U-shaped for light (Fig. S17A, Table S9). By contrast, six populations varied in a hump-shaped across the three resources (Fig. S17, Table S9).

Finally, *Tbr* varied between a minimum of 2℃ and a maximum of 20℃ (Fig. 3G-I). Overall, when averaged across all populations, *Tbr*s displayed a significatn U-shape with light availability, while nitrogen and phosphorus had linear increases (non-significant, Fig. S15H-I, Table S10). Individually, populations varied in the response of *Tbr* depending on the resource type. For over 52% of the populations, increasing light tended to reduce thermal breadth, linearly or monotonically (Fig. S15, Fig. S18). Six populations showed evidence of U-shape in *Tbr* across the light gradient (4 U-shape with critical points > 150 μmol photons·m-2·s-1). This is because light limitation tended to flatten the TPCs (Fig. S7), thereby increasing the range of temperatures at which a population achieved 80% of the maximal growth rate (*Tbr*). For nitrogen and phosphorus, more than 63% of all populations demonstrated increases in *Tbr* - either linear or monotonic (Fig. S15, Table S9). When TPCs were not flattened by resource limitation, but maintained similar curvature, then increasing resource availability tended to increase *Tbr*, e.g. for *Coelastrum* sp. -phosphorus-, *N. palea* (C&G) -nitrogen- (Figs. S8 & 9, Table S9).

### Associations of thermal and resource traits across species

The first two axes of the principal component analysis explained 61% of the variance of the data (Fig. 4A&B). The phylum to which phytoplankton population belonged was a significant predictor of the variance in the data (Fig. 4A, ANOSIM R = 0.1987, p-value < 0.005, Fig. S20). The resource type was, however, not a significant predictor of PCA scores, and explained less of the variation in the data than phylum (Fig. 4B, ANOSIM p-value=0.0541). Across all experiments, but particularly for light, Cyanobacteria had significantly higher maximum thermal optima (*Topt_max_*) than the other two phyla (Fig. 4C). Cyanobacteria also tended to have the lowest *R*_min_* for light (p-value < 0.05, Fig. 4D).

Populations’ *Topt_max_* were strongly positively associated with their *Tmax_max_* (Fig. 4A&B, Fig. 5A, Fig. S19). The *Tbr_max_* were positively associated with the *Tmax_max_*, indicating that the most warm-adapted populations could also be more generalist by having a wider thermal niche (Fig. 5B, Fig. S19). Populations tended to have *TR*_min_* that were ∼5 °C lower than the *Topt_max_*, i.e. *TR*_min_* values below the 1:1 dashed line Fig. 5C. *TR*_min_* was positively correlated with *Tmax_max_* (Fig. 4A&B, Fig. 5D, Fig. S19). This suggests that populations that maximize their growth rates at high temperatures and have high thermal tolerances also tend to be populations that have their lowest resource requirements at high temperatures. It should be noted, however, that as *Tmax* increases, a random sampling of temperatures between *Tmin* and *Tmax* at which *TR*_min_* could occur is likely to generate a positive relationship. The same applies for the relationship between *Tmax_max_*, and *Topt_max_* (Fig 5A). For these correlations with *Tmax_max_*, we have generated random expectations of distribution of Kendall’s Tau correlations for our data (see data package and Figs. S21, S22). The observed relationship between *Topt_max_* and *Tmax_max_*, is stronger than expected based on a random sampling of *Topt_max_*, constrained by *Tmax_max_* (Fig 5A, Fig. S21). However, the strength of the relationship between *TR*_min_* and *Tmax_max_* was within the distribution of the random expectations ( Fig. 5D, Fig. S22). Lastly, populations’ *Topt_max_* and *Tmax_max_* tended to be negatively associated with the lowest minimum *R\** values (*scaled log R*_min_*, Fig. 4A&B, Fig. 5E&F, Fig. S19). This suggests that the most warm-adapted taxa may also be the best resource competitors under limiting conditions. Overall, our analysis reveals that some populations in our pool have trait combinations that are beneficial under warm and resource-limited conditions: association high *TR*_min_* & high *Tmax_max_*; low *R*_min_* & high *Topt_max_*, or low *R*_min_* & high *Tmax_max_*. These can then be described as trait combinations adapted for future climate. The experimental resource manipulation did not interact significantly with any of the predictors of bivariate trait associations (Fig. 5, ANOVA interaction p > 0.05), and so all correlations were reported using data from all three experiments.

## Discussion

As lakes and oceans warm, resource availability in surface waters is changing concurrently due to extended stratification (Kraemer et al., 2021a; Li et al., 2020; Woolway & Merchant, 2019). Understanding which species will benefit or suffer from these combined effects of climate change is central to understanding and predicting how communities will reorganize under future climate. Our work demonstrates that minimum requirements of three essential resources for 19 populations of phytoplankton often vary with temperature. Warming may therefore alter the identity of the winner of resource competition. We also found that parameters of individual populations’ thermal performance curves often changed with reduced resource availability.

Our results suggest that a combination of resource limitation and high temperatures will tend to reduce the performance of individual populations relative to the conditions of optimal performance. Nevertheless, across all resources, we found that some populations with high thermal optima and high thermal maxima also have low minimum resource requirements. Furthermore, these warm-adapted populations also have their lowest resource requirements at high temperatures (Fig. 4A&B, Fig. 5E&F). These trait correlations, observed across our 19 populations, therefore indicate that some warm-adapted taxa also have traits that support their performance under resource limitation, combinations that will be beneficial under future climate.

### Resource niche variation with temperature

Our analysis indicates that the identity of the best competitor for light and phosphorus was stable across the measured temperature range (respectively *Synechococcus* sp. and *Coelastrum* sp.). However, competitive shifts were more common when individual populations were competing for nitrogen. Previous work has shown that competitive hierarchies among populations may change with temperature (Bestion et al., 2018b; Lewington-Pearce et al., 2019; Sunday et al., 2024; Tilman et al., 1981), but data limitations did not allow comparisons of the impacts of temperature under different limiting resources.

While temperature impacted *R\**s of most populations for all three resources, the way in which it influenced *R\**s, both when averaged across all populations, and when considering response variability among populations, differed depending on the identity of the limiting resource. Specifically, *I\**s and *P\**s tended most often to vary in a non-monotonic, U-shaped fashion with temperature (lending strong support for H1), while requirements for nitrogen showed greater variability among populations in their dependence on temperature (Fig. 2A-C, Fig. S11A-C, Table S7-S8). Biologically, there is no reason to expect linear relationships for any of the traits we explored, which is why in the majority of cases where a U-shape was not supported, it is still possible that a U-shape would have been observed if a larger temperature gradient would have been explored (e.g. monotonic or linear increasing or decreasing relationships).

Multiple studies on the ecological effects of warming have focused on estimates of population growth under non-limiting resource conditions (Bennett et al., 2018; Butterwick et al., 2004; Dell et al., 2011; Lürling et al., 2013; Savage et al., 2004). This corresponds to the parameter *µ_max_* in the standard Monod equation. The equivalent parameter in our study, *net-µ_max_*, tended to show a hump-shaped relationship with temperature, confirming that our methodology can recover the well-established shape of a TPC, and supporting H2. This finding was consistent both among populations and resources. Almost all exceptions to a hump-shaped relationship were positive linear or monotonic, suggesting that *Topt* occurred above our range of temperatures. Cases in which a linear decrease were inferred had high parameter uncertainty (wide credible intervals, Fig. S13). The impact of temperature on *R\** showed greater variation in shape than the impact of temperature on *net-µ_max_* (Fig. 2A-F, Fig. S11A-F). Temperature also significantly affected affinity (*α*), with large variations in both shape and direction, especially for phosphorus (Fig. 2I, Fig. S11I), suggesting that affinity is a parameter that is extremely hard to constrain well, and variation should be interpreted with caution. It is also possible that the difference among resources in how temperature influences *R\** may be largely determined by how temperature affects affinity. Prior studies have also demonstrated that temperature impacts low resource-sensitivity (e.g. *Ks* (Bestion et al., 2018c) and *α* (Lewington-Pearce et al., 2019)). Future physiological studies that specify the mechanistic underpinnings of differences in how *α* varies with temperature for resources would be valuable.

### Temperature niche variation with resources

Resource limitation tended to reduce the thermal optimum and maximum of populations, significantly for light and phosphorus (*Topt* & *Tmax* respectively), lending support to H3 and the ‘Metabolic Meltdown Hypothesis’. This hypothesis states that due to the higher metabolic costs and concurrent reduced access to resources caused by warming -for ectotherms-population’s optimal temperature for performance and fitness are expected to decline. Overall, this means that reduced resource availability may result in a reduction in population’s thermal niche size, potentially causing extirpation (Huey & Kingsolver, 2019). This hypothesis pertains to ectotherms, and similar findings have been documented in a marine copepod (Rueda Moreno & Sasaki, 2023), a mosquito (Huxley et al., 2021), and marine phytoplankton (Thomas et al., 2017). Furthermore, this suggests that resource limitation is likely to reduce the pool of populations that can withstand the increased strength and prevalence of extreme heat events projected to occur under all future climate change scenarios (Woolway et al., 2021). Thermal breadths varied greatly among populations and resources in how they responded to resource availability. Often thermal performance curves were flattened with lower resource availability, particularly for light, as found in previous data synthesis (Edwards et al., 2016), which had the consequence of increasing the thermal breadth. At least one previous study found that light limitation reduces the sensitivity (*E_a_* or activation energy) of phytoplankton growth rates to temperature more often than nitrogen or phosphorus limitation, which is consistent with our finding (Weber de Melo et al., 2025). This finding counters our initial hypothesis that thermal breadth would decline with resource limitation (H4) but may also be a consequence of how we defined thermal breadth. We recommend further exploration of how various definitions of thermal breadth (e.g. thermal range

= *Tmax*-*Tmin*, among others) respond to resource limitation, when possible. There were, however, several cases for nitrogen where thermal breadths increased with resource availability, in particular when the curvature of the TPC remained similar across resource levels, supporting H4. In such cases, limiting resource concentrations make populations more sensitive to environmental temperature (i.e. more thermally specialist), and limiting resources may lead to more rapid turnover of populations along thermal gradients than under non-limiting conditions. Such consequences have been documented in controlled experiments with ciliates and rotifers, where resource competition reduced the temperature at which focal populations went locally extinct, as well as the time to local extinction (Walberg, 2024).

### Resource and thermal trait correlations and implications for future climate

Our findings suggest that individual populations may suffer from greater resource requirements with warming and reduced thermal optima and maxima with limiting resources. Nevertheless, some populations have trait combinations conferring a competitive advantage in a warmer and more resource-limited future. For example, the Cyanobacteria *M. aeruginosa* strain 1 and the Chlorophyta *P. boryanum* both had high maximum thermal optima and among the lowest minimum requirements for light and nitrogen, respectively (Fig. 5E). Furthermore, we confirmed previous findings suggesting that the temperature at which populations have their minimum *R\**s (*TR*_min_*) are “cold-shifted” relative to the temperature at which populations grow fastest under non-limiting conditions (*Topt_max_*, Fig. 5C, most points below the 1:1 line, (Sunday et al., 2024). So, while combined warming and resource limitation are likely to limit the distribution and abundance of many phytoplankton species, some populations will suffer less from the combination of warming and resource limitation. Our data suggest that Cyanobacteria will be selected both by light limitation and warming, whereas green Chlorophyta may generally be stronger competitors under nitrogen and phosphorus limitation (Fig. 2B-C, Fig. 4D). Our pool of cyanobacterial populations is rather small (3 populations, two from the genus *Microcystis*), and further validation is required to confirm our results, however, our inferences are still supported by literature (Paerl & Huisman, 2008). Cyanobacteria are generally of poor nutritional quality for grazers (Ahlgren et al., 1990; Lampert, 1987), and can form toxic blooms at great cost for the provisioning of ecosystem services (Cheung et al., 2013; Watson et al., 2015). Characterizing the environmental conditions that select for these populations is therefore important for management actions (e.g. reduction of nutrient loads to lakes) aimed at maintaining the provisioning of important ecosystem services (e.g. drinking water (Jing et al., 2018)) under future climate (Doubek et al., 2015; Schampera et al., 2024; Schampera & Hellweger, 2024).

### Limitations and future directions

This new dataset provides several novel insights, but our inferences are limited by the scope and parameters that we investigated. Most importantly, our findings are determined by the populations in our pool, and may change if, for example, more tropical or polar populations would be included. Specifically, we recommend that future studies investigate a greater number of cyanobacterial species due to the potentially important role they may play in warmed waters. In particular, broader species pools would enable the confirmation of whether there is generally less turnover in the identity of the best competitor for light and phosphorus across thermal gradients than there is for nitrogen. We also recommend that future studies investigate wider experimental thermal ranges to more completely describe TPCs and provide accurate estimates of *Tmin*. Additional temperatures would also enable more accurate inferences with GAMs; hence reducing the number of linear patterns described.

Our analysis of parameters and their variation across environmental gradients represents multiple levels of analysis through which error can be propagated. In the future, it may be possible to more directly characterize the impacts of temperature and resources on growth rates using either parametric or non-parametric response surfaces (van Moorsel et al., 2023), rather than analyzing the variation of parameters across environmental gradients. It may be possible to analyze density directly, rather than growth rates, to extract population parameters (Palamara et al., 2014). Further development of statistical methods to reduce or quantify the propagation of errors would be commendable. Nevertheless, fundamentally reducing error to improve statistical inference may require sample sizes that are prohibitive (we analyzed > 10,000 growth rate estimates). We expect that patterns that are repeated across large species pools, and supported by theory, are likely to be robust, despite statistical uncertainty.

Our high-throughput approaches, came at the cost of the optimization of the experimental treatments. Nevertheless, reducing resources and increasing temperature may both concurrently impact density-independent growth rates (our focus), but also the carrying capacity of populations (Bernhardt et al., 2018a). This concurrent change may constrain our ability to isolate the response of the growth rates at increasingly limiting resource levels and elevated temperatures. As resources become more limiting and carrying capacities drop, estimates of growth rates in batch culture experiments such as ours should be performed on single-individuals in large volumes and at infinitesimally small time-intervals to avoid the impact of density dependence. Given that this may not be feasible, we recommend the further development of methods that enable the separation of density-independent and density-dependent effects of both temperature and resource availability.

Finally, this dataset could be used to parameterize community models and to make predictions of shifting species composition (Bestion et al., 2018a). This was beyond the scope of the current study, but our data could be used to generate model predictions to be validated with empirical tests. *R\**s alone make predictions about the winner of competition under a single limiting resource, but information on resource consumption ratios is also needed when multiple resources are limiting. Studies investigating how species’ resource consumption rates vary with temperature are required for this purpose (Yvon-Durocher et al., 2017). Finally, Resource Competition Theory is focused on making predictions under equilibrium conditions, but this is an oversimplification for making predictions for natural systems. It will be important to determine to what extent the traits we investigated here can be used to make predictions under fluctuating resource and thermal environments (Bernhardt et al., 2018b; Litchman et al., 2004; Thompson et al., 2018).

### Conclusion

We present an analysis of temperature-resource interactions on phytoplankton growth rates for the greatest number of populations and resources studied within a single empirical workflow to-date. This allowed an unprecedented investigation of how resources impact thermal traits, and, vice versa, how temperature impacts resource traits. Most importantly, our analysis has shown that across 19 populations, the lowest minimum resource requirements (*R\**) tend to be associated with warm-adapted populations (highest *Topt*). We infer that these are also the taxa that are most likely to succeed in future warm and resource-limited surface waters. A future challenge will be to integrate these trait data into predictive community and ecosystem models to make quantitative predictions of shifts in community composition, as well as the resulting changes in ecosystem functioning, including biomass production, nutrient cycling and food quality for consumers (Allen & Gillooly, 2009; Hutchins & Tagliabue, 2024). Physiological cell models may also provide promising mechanistic insight into the basis of our observed interactions (Armin & Inomura, 2021; Toseland et al., 2013), and may provide opportunities for constraining predictions of community and ecosystem-level models (Chien et al., 2023). Our understanding of observed temperature-resource interactions would also ideally be incorporated into Earth System Models, which currently oversimply the responses and roles of phytoplankton. This would enable climate projections with realistic scenarios of environmental change to incorporate the most important nonlinear responses of phytoplankton to these gradients (e.g. Benedetti et al., 2021; Henson et al., 2021; Holder & Gnanadesikan, 2023). Some steps have been taken in this direction, though based on the limited data available to date (e.g. (Thomas et al., 2017)), and it is clear that temperature-resource interactions can have major impacts on species distributions and therefore also to contributions to ecosystem functions. Much of this work has focused on marine phytoplankton, but it is important to apply these findings to lake ecosystems as well. Unlike marine systems, lake ecosystems may be more constrained by dispersal limitation, reducing their ability to track optimal environmental conditions. Lake communities may therefore be expected to respond more slowly to changing climate (Khaliq et al., 2024; McFadden et al., 2023; Pinsky et al., 2019).

## Acknowledgements

We thank Gabriella Mège for her work in isolating phytoplankton species from four Swiss lakes, Raphael Bossart for his work in confirming taxonomic identities with DNA sequencing and Pinelopi Ntetsika for running our scripts on the ETH Zurich High Performance Computing cluster. The project was financially supported by a Project Grant from the Swiss National Science Foundation (SNF) to AN (310030_197812/1), Flexibility Grants to VWM (310030_197812/2, 4) and a Mobility Grant to SL (310030_197812/3).

