## Supplemental Info - Figures & Tables for "Warm-loving species perform well under limiting resources: trait combinations for future climate"

**Title**

### Supplemental Methods:

*Species identification using barcoding*

The algal DNA of each population was extracted with a Xanthogenate DNA isolation protocol (Tillett & Neilan, 2000). Genotyping was done using 16S, 18S or ITS sequences, depending on the species (Table S2). These markers were amplified using a QIAGEN Multiplex PCR KIT, and the PCR products were sequenced by Microsynth using the Sanger method. The output sequences were compared to existing identifications using GenBank and NCBI BLAST.

*Acclimation to temperature and resource conditions*

Over ten days, we gradually increased the light from 40 to 110 µmol photons $m^{-2} \cdot s^{-1}$ (~20 µmol photons $m^{-2} \cdot s^{-1}$ every two days). We changed the temperature simultaneously by a maximum of 2°C per day until the 6 target temperatures were achieved. Once the light and temperature targets were reached, we waited 48 hours before starting the resource acclimation.

For the acclimation to nitrogen and phosphorus, the temperature-acclimated cultures were divided in two, and centrifuged at 2,000 rpm for 4 min, the supernatant was discarded and replaced by the corresponding medium (level R1 or R5) (Fig. S1). For the light experiment, the temperature-acclimated cultures were split in two, the light acclimation was done by setting the incubator light at the R5 level and the R1 level was achieved with the use of neutral density filters (Solar GraphicsTM, Clearwater, Florida). The acclimation to R1 was 48h and to R5 was 72h. After all the populations were temperature- and resource-acclimated, the R1 cultures were used to inoculate 96-well plates of levels R1, R2, R3 and R4, and the R5 cultures were used to inoculate plates of levels R5, R6, R7 and R8 (Fig. S1). In the light experiment, the incubators were set at R8 (Table 1), and the other levels were achieved through the use of custom-made 96-well plate neutral density filter covers (Solar Graphics$^{TM}$, Clearwater, Florida).

*Temperature x resource growth rate experiments*

The temperature range we chose to perform the experiment, aimed to represent as best as possible the different thermal niches of our 19 phytoplankton populations. The 4°C interval between 20-32°C aimed to target the core of the thermal range, while 15 and 35°C were considered, for most of our species, to be at or close to the thermal minimum and maximum.

Once the temperature and resource acclimation were completed, on the day of the start of the experiment, each population-temperature-resource level culture, was subsampled (200 µL) in order to measure the culture's relative fluorescence unit (RFU). We used a plate reader (Agilent Cytation 5) to estimate pigment fluorescence (measurement from the bottom of the wells), chlorophyll-a for the Chlorophyta and the Bacillariophyta (excitation wavelength was 445 nm and emission was 685 nm) and phycocyanin for the Cyanobacteria (excitation wavelength was 586 nm and emission was 647 nm). These RFU values from our temperature and resource acclimated populations, allowed us to create dilution tubes that served for the inoculation of the 96-well culture plate via the use of the TECAN robot (automatic liquid handler).

*Estimation of population growth rates*

We used the 'get.growth.rate' function (Kremer, 2020) on the whole fluorescence time series (with few exceptions) to extract the exponential growth rate using one of five models: a simple exponential model, a model with a lag phase, a model with a saturation phase, a model incorporating both lag and saturation phases, and a logistic growth model. These models allow for flexibility in accounting for different growth dynamics, including delayed onset of growth (lag phase) and potential limitations as the system approaches carrying capacity (saturation phase). The function chooses the best model to fit the data and provide a growth rate via comparison of the AIC values. For time series where a population grew quickly and then declined, time series were truncated using a custom R script, to eliminate the declining portion of the curve because the get.growth.rate function does not handle such growth dynamics. Specifically, for time series where there were 4 or more decreasing data points after maximum fluorescence, growth curves were fit only until the maximum fluorescence was achieved. Time series truncation was required for fewer than 2% of the growth curves. All time-series retained at least 5 time points. The target fluorescence to start the experiment was 20 RFU. High starting densities limit our ability to observe exponential growth and can also cause bias in the estimated population growth rates (Pylvänäinen, 2005). For this reason, when the inoculation density for any replicate was above 40 RFU, the time series was removed from the analysis. Across the three experiments, this last filter eliminated 1.6 % of the time series. Note that growth rates at low nutrient levels may be underestimated if the inoculum density is a significant fraction of the carrying capacity, particularly if uptake rates exceed growth rates and nutrient storage is possible. Consequently, for each experiment, following the estimation of growth rates, we determined the temperature at which each population exhibited the greatest number of resource levels with consistent positive growth across all four replicates ($\mu > 0$). Subsequently, for each population, we excluded resource levels at temperatures that did not fulfill this criterion, these filtered out negative growth rates could be due to either inoculation above a lowered carrying capacity or because the resource level doesn't allow cell survival. For example, consider the nitrogen experiment and the Chlorophyta species *Eudorina elegans*. Suppose 24°C was identified as the temperature at which the population exhibited the highest number of nitrogen levels where all four replicates showed positive growth ($\mu > 0$). In this scenario, nitrogen levels N2 through N8 met the criterion, therefore, we retained nitrogen levels N2 to N8 and applied this selection consistently across all temperature treatments. This consistent filtering across temperatures ensured temperature-dependent growth limitations (i.e., negative growth at certain temperatures) were not inadvertently excluded from the analysis.

*Estimation of Monod parameters*

For the Monod fits the filter described above led to the loss of 4 population for the nitrogen experiment (*A. formosa, Chlorella* sp.*, C. pseudostelligera and Synechococcus* sp.) because they were left with only 2-3 resources levels, which was not enough data to allow the fit of a Monod model. For the phosphorus fits, only one population was filtered out due to this rule (*C. pseudostelligera*), no change for the light experiment. An additional filter was necessary, for each experiment-species combo, we removed any temperature treatments where there were not at least 4 resource levels where the growth rate was positive.

**Table S1.** Summary table of all the parameters used and traits investigated in our study and their definitions.

| Parameter: | Parameter name: | Parameter definition: | Equation: |
|---|---|---|---|
| $\mu_{max}$ | Gross maximum specific growth rate | Maximum value for growth as the resource increases to infinity | lim $\mu \rightarrow$ R $\infty$ |
| $\mu$ | Population growth rate | Population growth as a function of resource availability. | $\left(\frac{\mu max * R}{Ks + R}\right) - m$ |
| $\alpha$ | Affinity | The initial slope of the Monod curve | $\alpha = \frac{\mu max}{Ks}$ |
| $R*$ | Minimum resource requirement | Amount of a resource to have population growth rate equal to external loss rate ($m_{ext}$) | R $\rightarrow$ $\mu = m_{ext}$ also, $R* = \frac{mtot * Ks}{\mu max - mtot}$ |
| net-$\mu_{max}$ | Net maximum specific growth rate | Gross maximum specific growth rate corrected for intrinsic loss rate ($m$) | net-$\mu_{max}$ = $\mu_{max}$ - $m$ |
| $T_{opt}$ | Thermal optimum | Temperature where growth rate is maximized if resource is not limiting | $\mu_{max}$ (T) |
| $T_{min}$ | Thermal minimum | Lower temperature limit for population growth rate at which growth crosses zero | $\mu$ ($T_{min}$) = 0 |
| $T_{max}$ | Thermal maximum | Upper temperature limit for population growth rate at which growth crosses zero | $\mu$ ($T_{max}$) = 0 |
| $T_{br}$ | Thermal breadth | Temperature range where growth is equal or greater than 80% of the maximal growth rate at *Topt.* | $\mu$ (T) $\geq 0.8 \times \mu_{max}$ for T $\in$ [$T_{min}$*, $T_{max}$*] $T_{min}$*, $T_{max}$* temperature at which growth = 80% $\mu_{max}$. |
| $R*_{min}$ | Lowest $R*$ | Lowest minimum resource requirement across all temperatures | min ($R*$) |
| *Scaled log* ($R*_{min}$) | Scaled logged lowest $R*$ | Log-transformed and standardized $R*_{min}$ | log($R*_{min}$) and |
| $TR*_{min}$ | Temperature of the lowest $R*$ | Temperature at which $R*_{min}$ occurred | $R*_{min}$ (T) |
| net- $\mu max_{max}$ | Highest net maximum growth rates | Highest net maximum growth rates | max (net- $\mu_{max}$) |
| $Topt_{max}$ | Highest thermal optimum | Across resource levels, the maximum value for thermal optimum. | max ($T_{opt}$) |
| $Tmax_{max}$ | Highest thermal maximum | Across resource levels, the maximum value for thermal maximum. | max ($T_{max}$) |
| $Tmin_{min}$ | Lowest thermal minimum | Across resource levels, the minimum value for thermal minima. | min ($T_{min}$) |
| $Tbr_{max}$ | Highest thermal breadth | Across resource levels, the maximum value for thermal breadth. | max ($Tbr$) |

**Table S2**. Summary of sampled lakes, their trophic status and phosphorus load. Values are from literature.

| Lake | Trophic status | Phosphorus load | Reference |
|---|---|---|---|
| Greifen | eutrophic | 96 µg $L^{-1}$ | (Pomati et al., 2019) |
| Lower Lake Zurich | mesotrophic | 33 µg $L^{-1}$ | (Pomati et al., 2019) |
| Lucerne | meso-oligotrophic | 9 µg $L^{-1}$ | (Pomati et al., 2019) |
| Constance | oligotrophic | 7.5 µg $L^{-1}$ | (Saboret et al., 2023; Wörner & Pester, 2019) |

**Table S3**. Species list used during the 3 experiments, with their phylum lake origin and marker for identification.

| Species | Short name | Phylum | Lake | Marker |
|---|---|---|---|---|
| *Tetraedron minimum* | *T. minimum* | Chlorophyta | Lake Greifen | 18S |
| *Oocystis* sp. | *Oocystis* sp. | Chlorophyta | Lake Greifen | ITS1-5.8S-IST2 |
| *Scenedesmus armatus* | *S. armatus* | Chlorophyta | Lake Zurich | ITS1-5.8S-IST2 |
| *Coelastrum* sp. | *Coelastrum* sp. | Chlorophyta | Lake Zurich | 18S |
| *Messastrum gracile* | *M. gracile* | Chlorophyta | Lake Zurich | ITS1-5.8S-IST2 |
| *Lagerheimia subsalsa* | *L. subsalsa* | Chlorophyta | Lake Constance | rcbl |
| *Pediastrum boryanum* | *P. boryanum* | Chlorophyta | Lake Constance | ITS1-5.8S-IST2 |
| *Scenedesmus arcuatus var. platydiscus* | *S. arcuatus* | Chlorophyta | Lake Constance | 18S |
| *Chlorella* sp. | *Chlorella* sp. | Chlorophyta | Lake Lucern | ITS1-5.8S-IST2 |
| *Eudorina elegans* | *E. elegans* | Chlorophyta | Lake Lucern | ITS1-5.8S-IST2 |
| *Fistulifera saprophila* | *F. saprophila* | Bacillariophyta | Lake Greifen | 18S |
| *Cyclotella pseudostelligera* | *C. pseudostelligera* | Bacillariophyta | Lake Greifen | 18S |
| *Nitzschia palea* | *N. palea (G)* | Bacillariophyta | Lake Greifen | 18S |
| *Fragilaria crotonensis* | *F. crotonensis* | Bacillariophyta | Lake Constance | ITS1-5.8S-IST2 |
| *Asterionella formosa* | *A. formosa* | Bacillariophyta | Lake Constance | 18S |
| *Nitzschia palea* | *N. palea (C)* | Bacillariophyta | Lake Constance | 18S |
| *Microcystis aeruginosa* Strain 1 | *M. aeruginosa* st. 1 | Cyanobacteria | Lake Greifen | 16S |
| *Microcystis aeruginosa* Strain 2 | *M. aeruginosa* st. 2 | Cyanobacteria | Lake Greifen | 16S+PCbetaIGS |
| *Planktothrix agardhii* | *P. agardhii* | Cyanobacteria | Lake Greifen | 16S |
| *Synechococcus* sp. CCAP 1479/11 | *Synechococcus* sp. | Cyanobacteria | Culture Collection | NA |

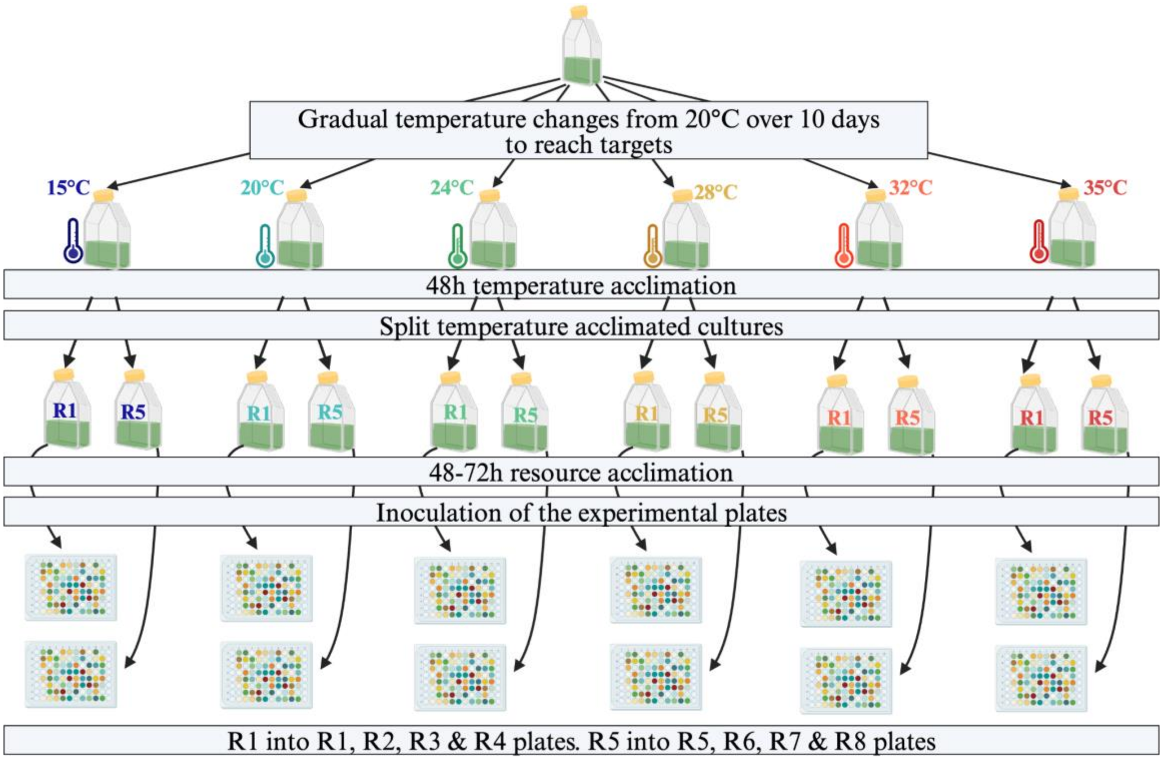

**Figure S1**. Experimental design and steps of the growth experiments, from batch cultures through temperature and resource acclimations and experimental inoculations. Each resource was investigated separately, so this design was repeated three times for all the species. From the start of the temperature change until the start of the experiment, the process took 15 days. For the acclimation to nitrogen and phosphorus, the temperature-acclimated cultures were divided in two, and centrifuged at 2,000 rpm for 4 min, the supernatant was discarded and replaced by the corresponding media level (R1 or R5).

**Table S4:** List of priors for the Monod fits with Bayesian statistics. The values for the gross maximum specific growth rate ($\mu_{max}$) and the parameter indicating the rate of death (*m*) are the same across resources, however, the half-saturation constant for growth (*Ks*) are resource specific.

| Resource | Parameter | Distribution | Mean | Standard deviation | Lower bound |
|---|---|---|---|---|---|
| Light | $\mu max$ | normal | 1 | 0.9 | 0 |
| Light | *Ks* | lognormal | 2.5 | 0.7 | 0 |
| Light | *m* | normal | 0 | 0.7 | 0 |
| Nitrogen | $\mu max$ | normal | 1 | 0.9 | 0 |
| Nitrogen | *Ks* | lognormal | 2.5 | 0.9 | 0 |
| Nitrogen | *m* | normal | 0 | 0.7 | 0 |
| Phosphorus | $\mu max$ | normal | 1 | 0.9 | 0 |
| Phosphorus | *Ks* | lognormal | 0.001 | 0.8 | 0 |
| Phosphorus | *m* | normal | 0 | 0.7 | 0 |

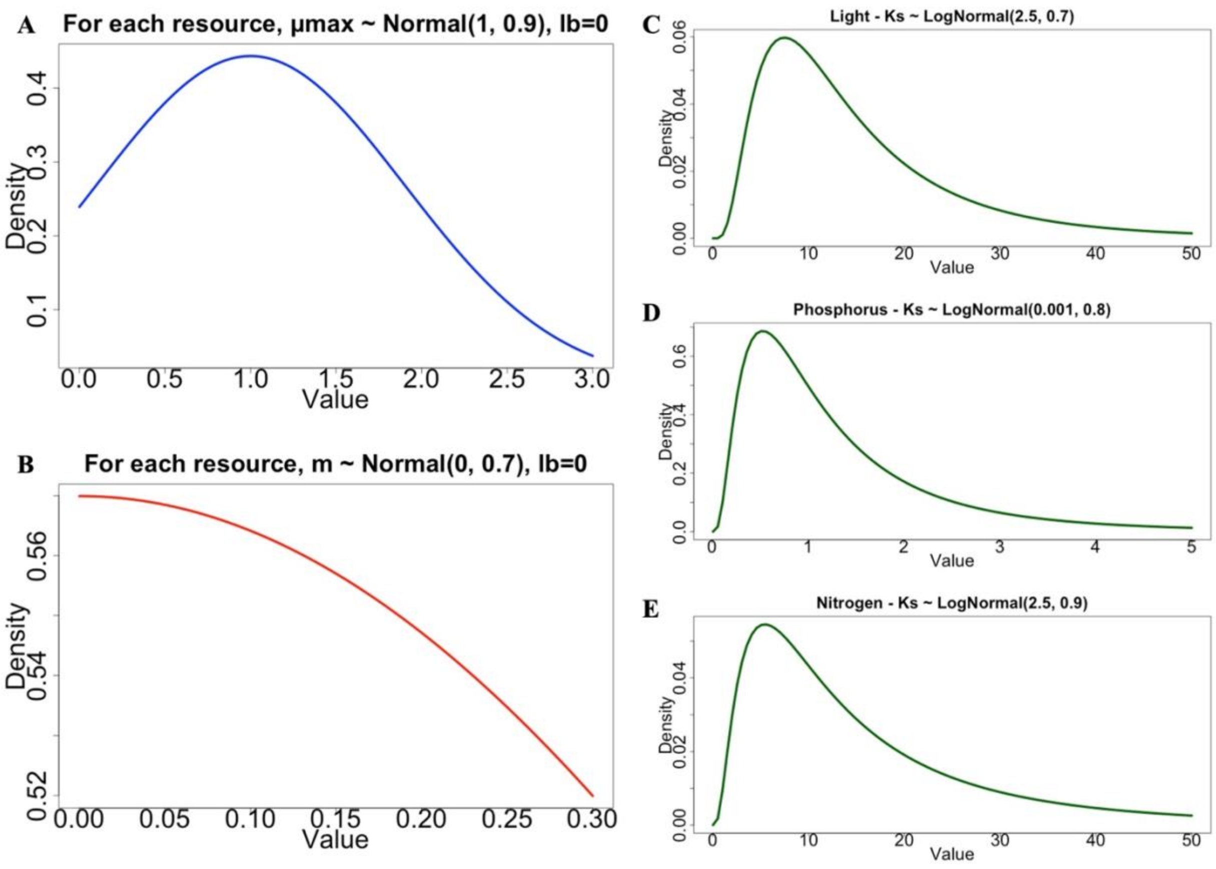

**Figure S2**. The priors' distribution of the Monod parameters for the Bayesian fitting. The priors mean and standard error values for the gross maximum specific growth rate ($\mu_{max}$, A) and the intrinsic loss rate (*m*, B) are the same across the three resources. The priors mean and standard error values for the half-saturation constant for growth (*Ks*) differ for each experiment, (C) for the light, (D) for the phosphorus and (E) for the nitrogen.

**Table S5.** Modified equation 5 used for for Bayesian fitting of the thermal performance curve (TPC).

| Model name | Equation | Definition |
|---|---|---|
| Modified Double exponential model from (Kremer et al., 2024) | $\mu \sim -d_0 + ((d_0 + \mu max) * (-((b_2 * exp((T - Topt) * (b_2 + \Phi)))/\Phi) + ((exp(b_2 * (T - Topt)) * (b_2 + \Phi))/\Phi)))$ | $d_0$ is the temperature-independent loss rate; $\mu max$ gross maximum specific growth rate; $b_2$ is the exponential change in birth rate as temperature increases; $T$ is the environmental temperature; $Topt$ the temperature where the growth rate is maximized; $\Phi$ (phi) = $b_2$- $d_2$; $d_2$ is the exponential change in death rate as temperature increases. |

**Table S6.** List of priors for the thermal performance curves fits with Bayesian statistics. The priors were the same for the three resources (light, nitrogen, phosphorus). The parameters are the following: *d0*, the temperature-independent death rate; $\mu max$, the gross maximum specific growth rate; *b2,* the exponential change in birth rate as temperature increases; *Topt*, the species' thermal optimum; $\Phi$ (phi) is *b2*- *d2*, with *d2* being the exponential change in death rate as temperature increases; finally σ, sigma, is the error term.

| Parameter | Distribution | Mean | Standard deviation | Lower bound |
|---|---|---|---|---|
| *d0* | lognormal | 1.41 | 1.58 | 0 |
| $\mu_{max}$ | lognormal | 0.95 | 1.41 | 0 |
| *Topt* | normal | 24 | 4.5 | -3.8 |
| *b2* | lognormal | 1.22 | 1.29 | 0 |
| $\Phi$ | lognormal | 1.22 | 1.65 | 0 |
| σ | lognormal | -2.7 | 0.6 | 0 |

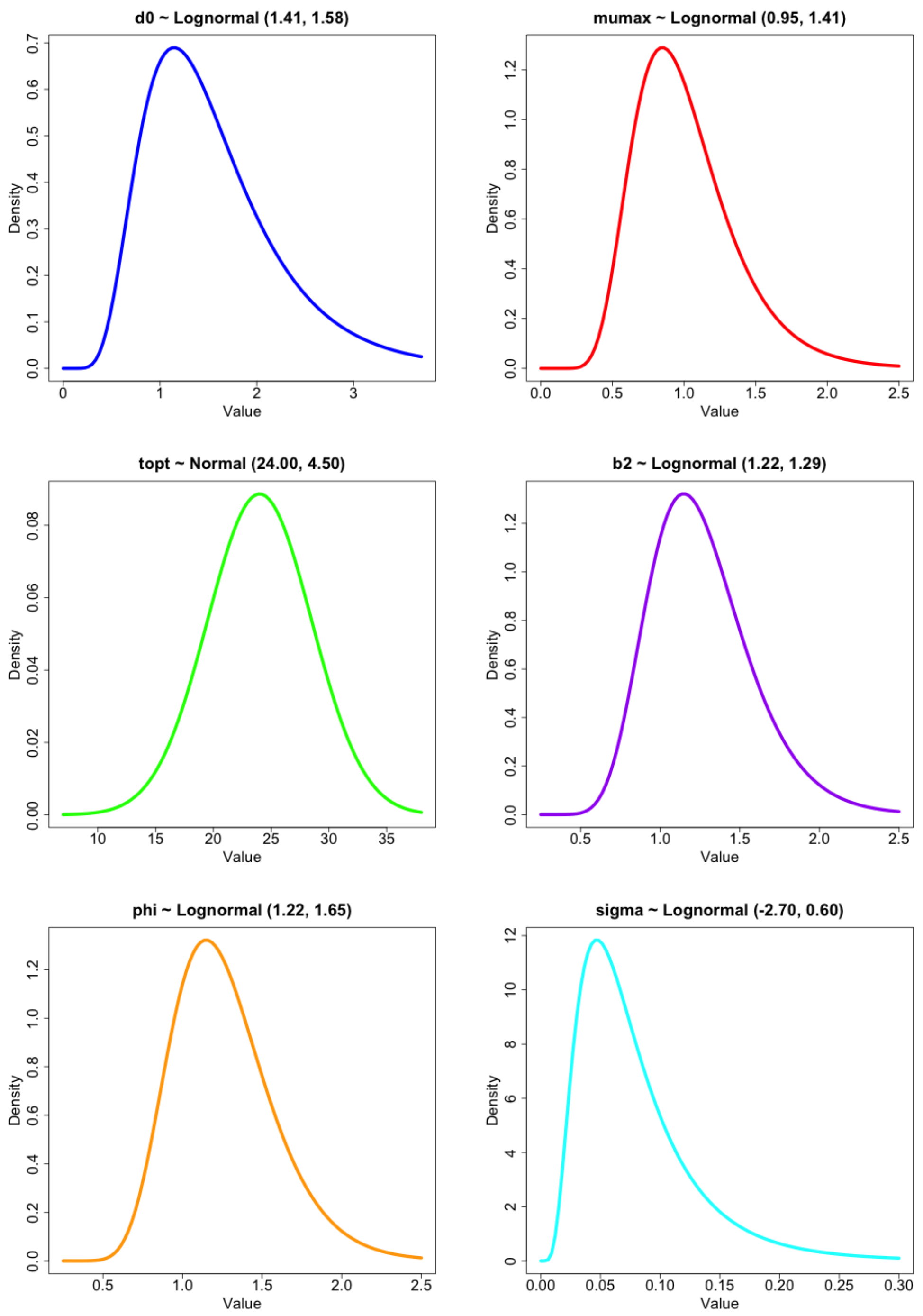

**<u>Figure S3</u>**. The priors' distribution of the Thermal Performance Curves (TPC) parameters for the Bayesian fitting. All the prior mean and standard error values are the same across light, nitrogen and phosphorus.

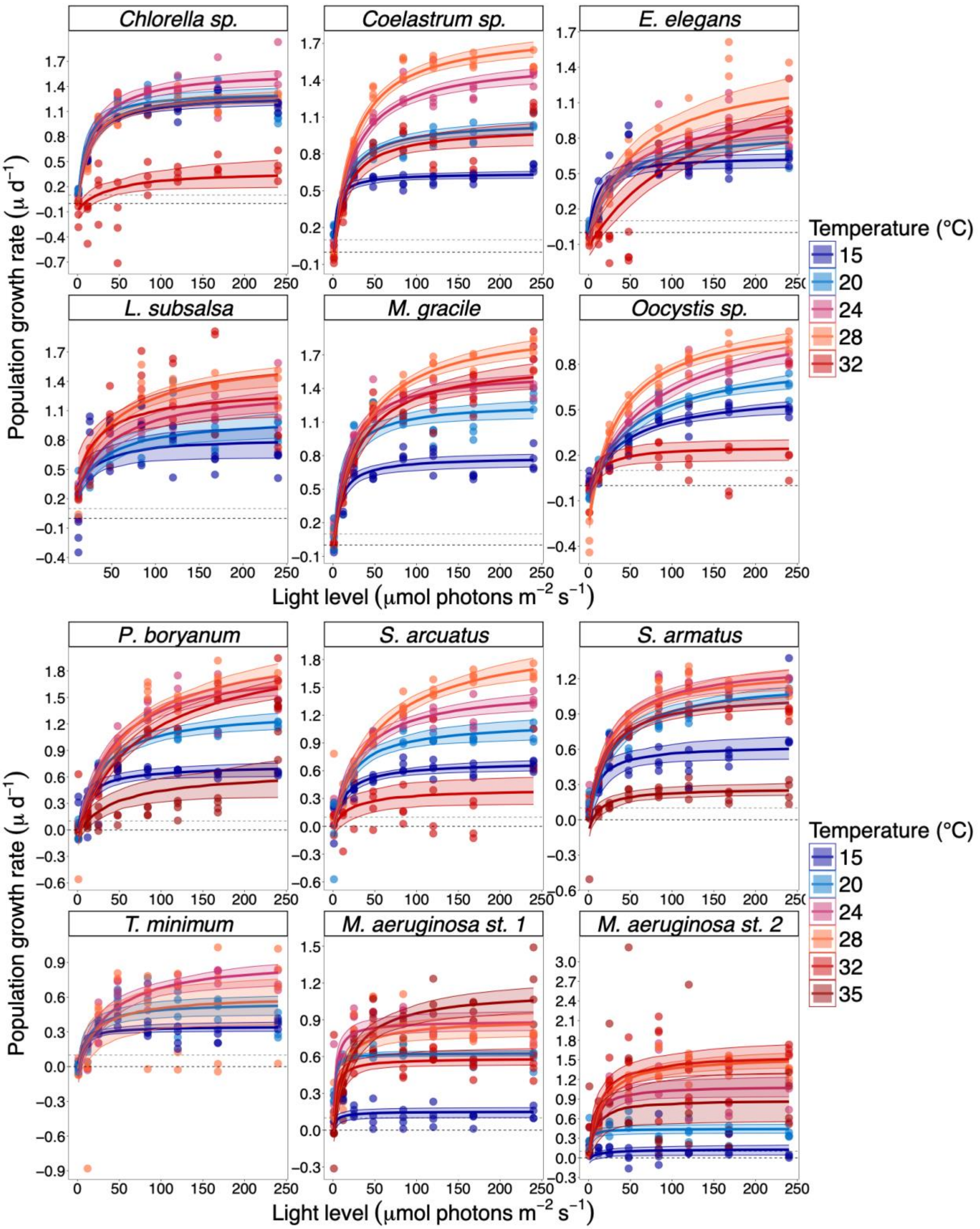
Chlorella sp.
Coelastrum sp.
E. elegans
L. subsalsa
M. gracile
Oocystis sp.
Population growth rate (μ d⁻¹)
Light level (μmol photons m⁻² s⁻¹)
Temperature (°C)
15
20
24
28
32
P. boryanum
S. arcuatus
S. armatus
T. minimum
M. aeruginosa st. 1
M. aeruginosa st. 2
Population growth rate (μ d⁻¹)
Light level (μmol photons m⁻² s⁻¹)
Temperature (°C)
15
20
24
28
32
35

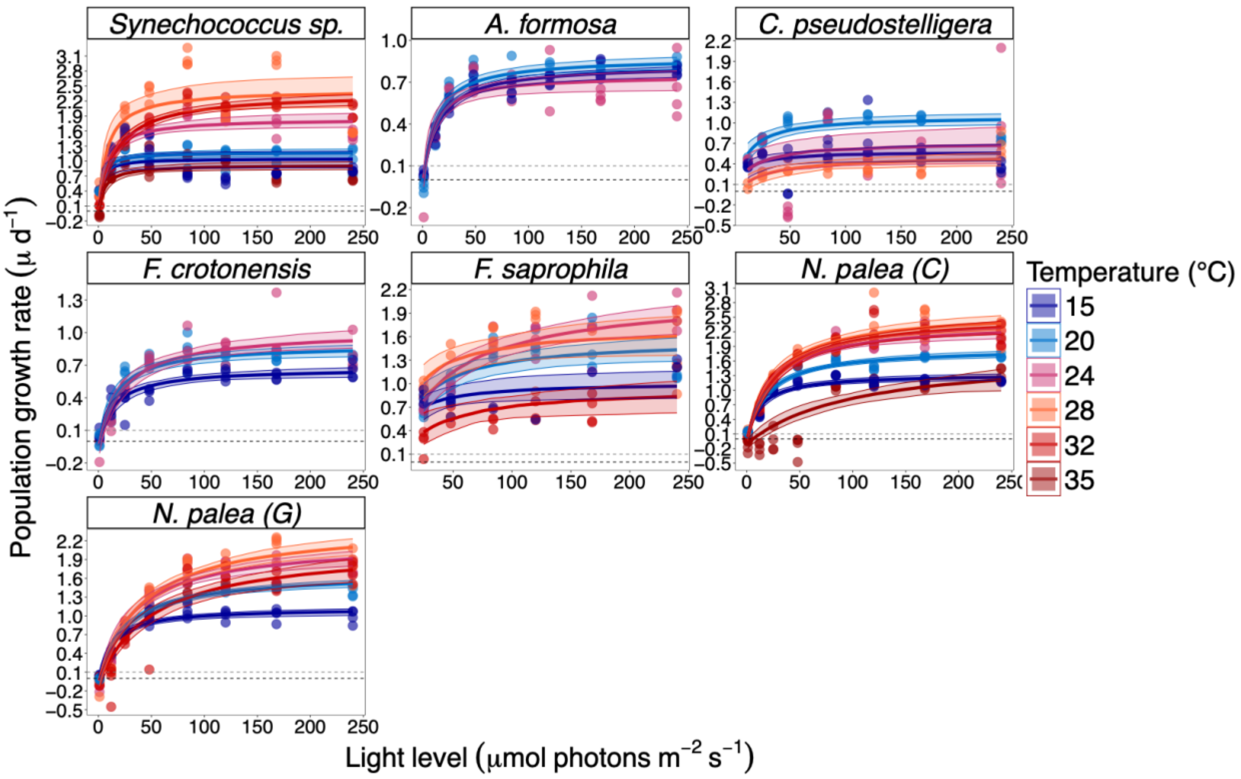

**Figure S4**. Population growth rates (day$^{-1}$) as a function of light availability for 19 populations of phytoplankton. Monod curves are fitted for each of the six different temperatures separately. Shaded ribbons indicate Bayesian 95% credible intervals. The black dotted line represents the $\mu = 0$ line, and the grey dotted line is the $m_{ext} = 0.1$ line.

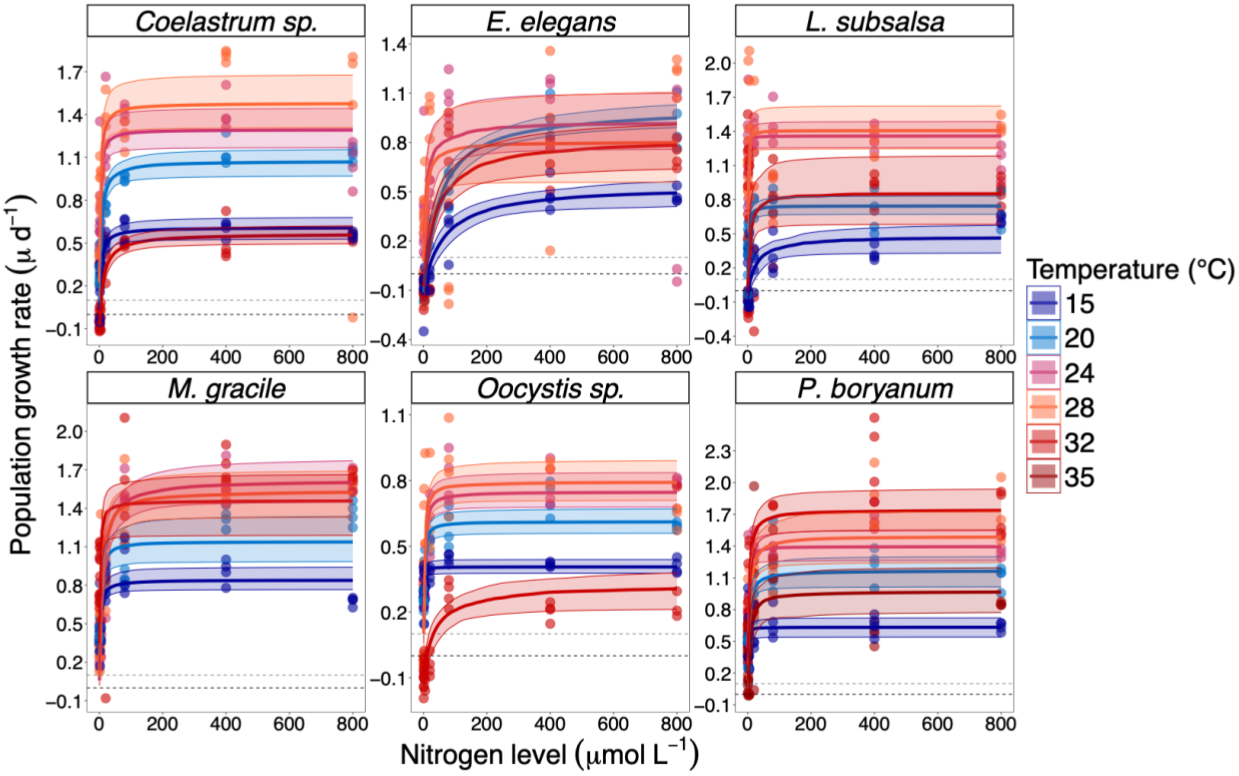
Coelastrum sp.
E. elegans
L. subsalsa
M. gracile
Oocystis sp.
P. boryanum
Population growth rate (μ d$^{-1}$)
Nitrogen level (μmol L$^{-1}$)
Temperature (°C)
15
20
24
28
32
35

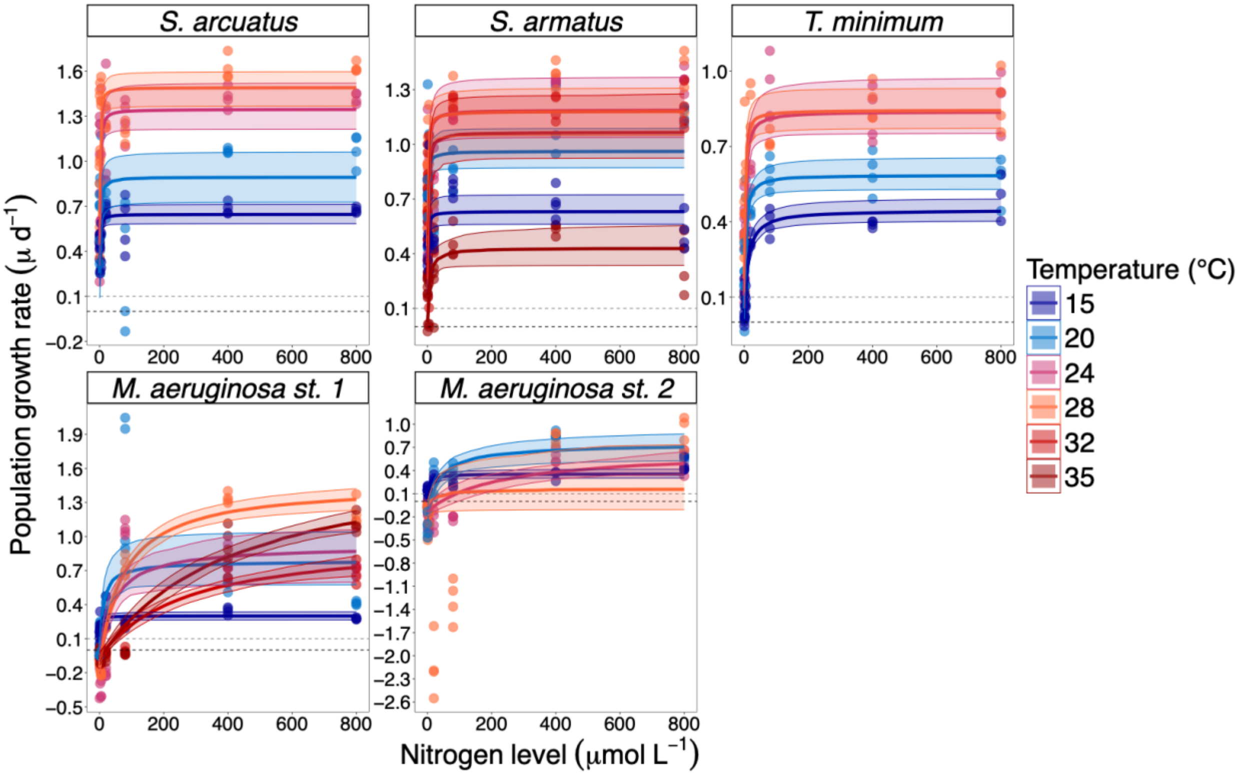
S. arcuatus
S. armatus
T. minimum
M. aeruginosa st. 1
M. aeruginosa st. 2
Population growth rate (μ d$^{-1}$)
Nitrogen level (μmol L$^{-1}$)
Temperature (°C)
15
20
24
28
32
35

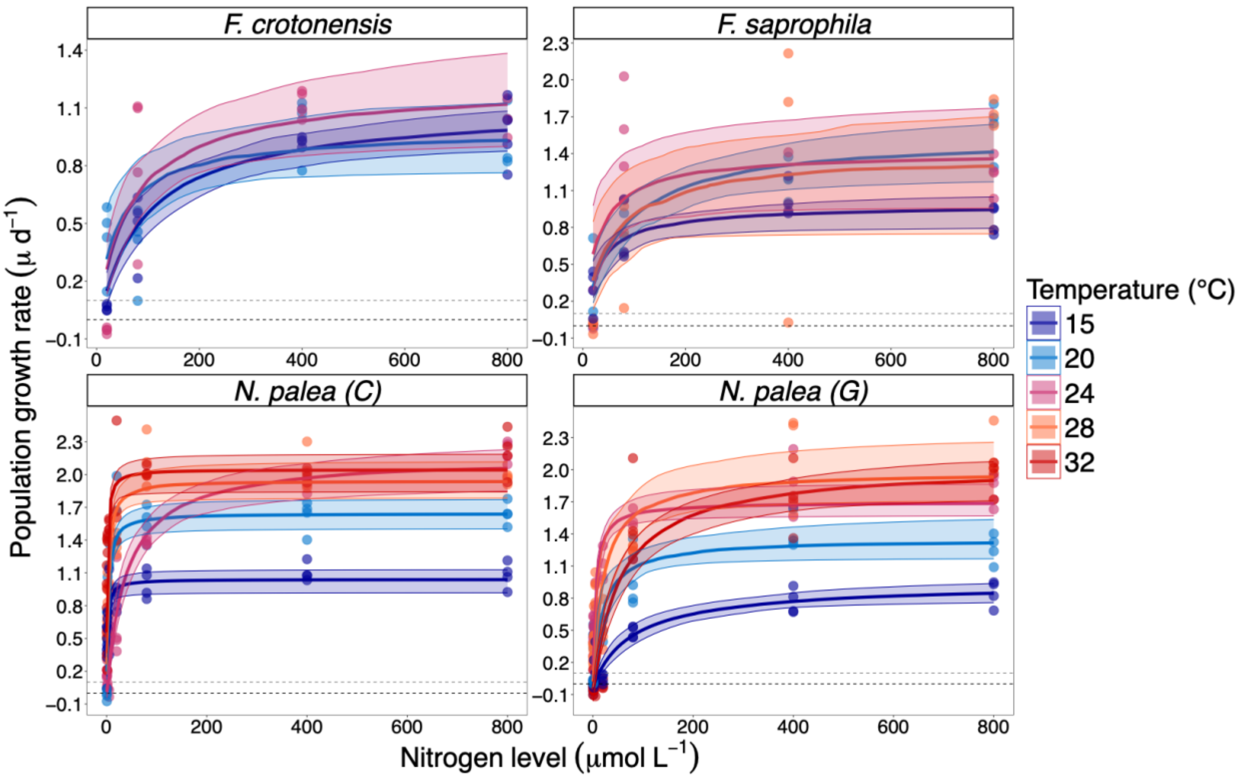

**Figure S5**. Population growth rates (day$^{-1}$) as a function of nitrogen availability for 19 populations of phytoplankton. Monod curves are fitted for each of the six different temperatures separately. Shaded ribbons indicate Bayesian 95% credible intervals. The black dotted line represents the $\mu = 0$ line, and the grey dotted line is the $m_{ext} = 0.1$ line.

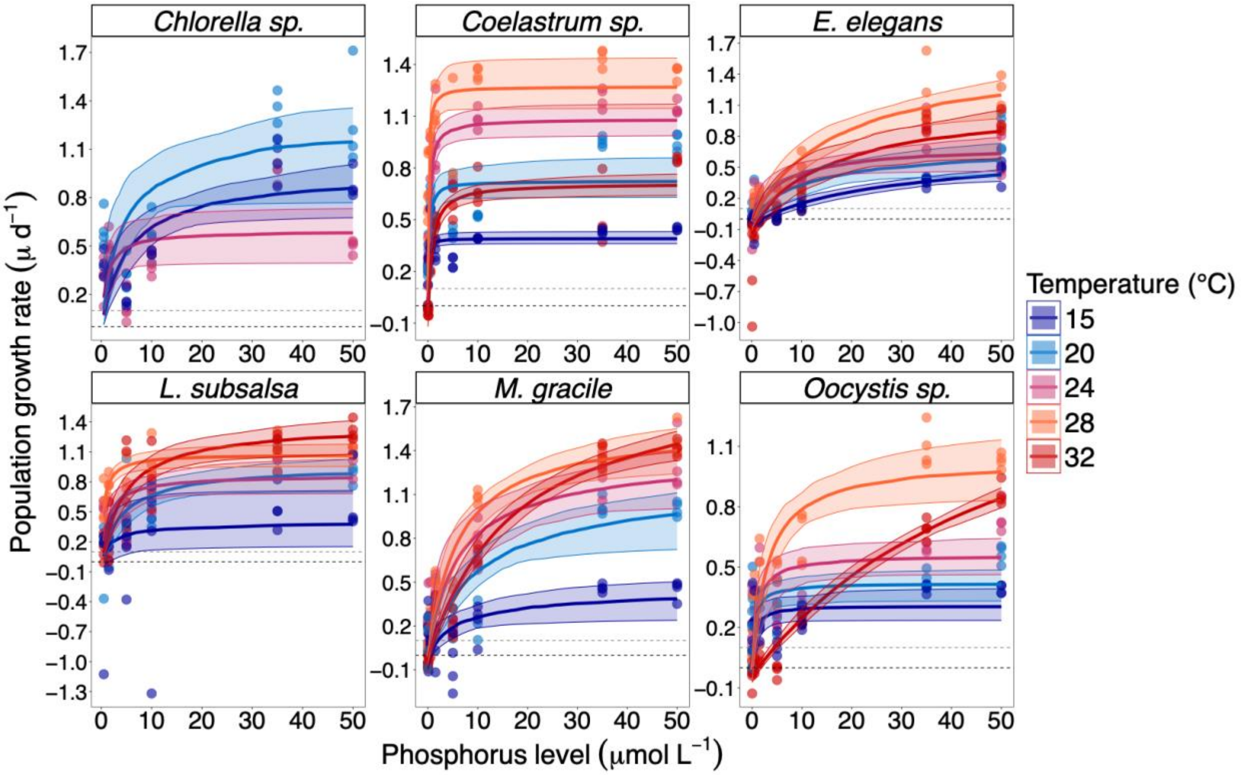
Chlorella sp.
Coelastrum sp.
E. elegans
L. subsalsa
M. gracile
Oocystis sp.
Population growth rate (μ d$^{-1}$)
Phosphorus level (μmol L$^{-1}$)
Temperature (°C)
15
20
24
28
32

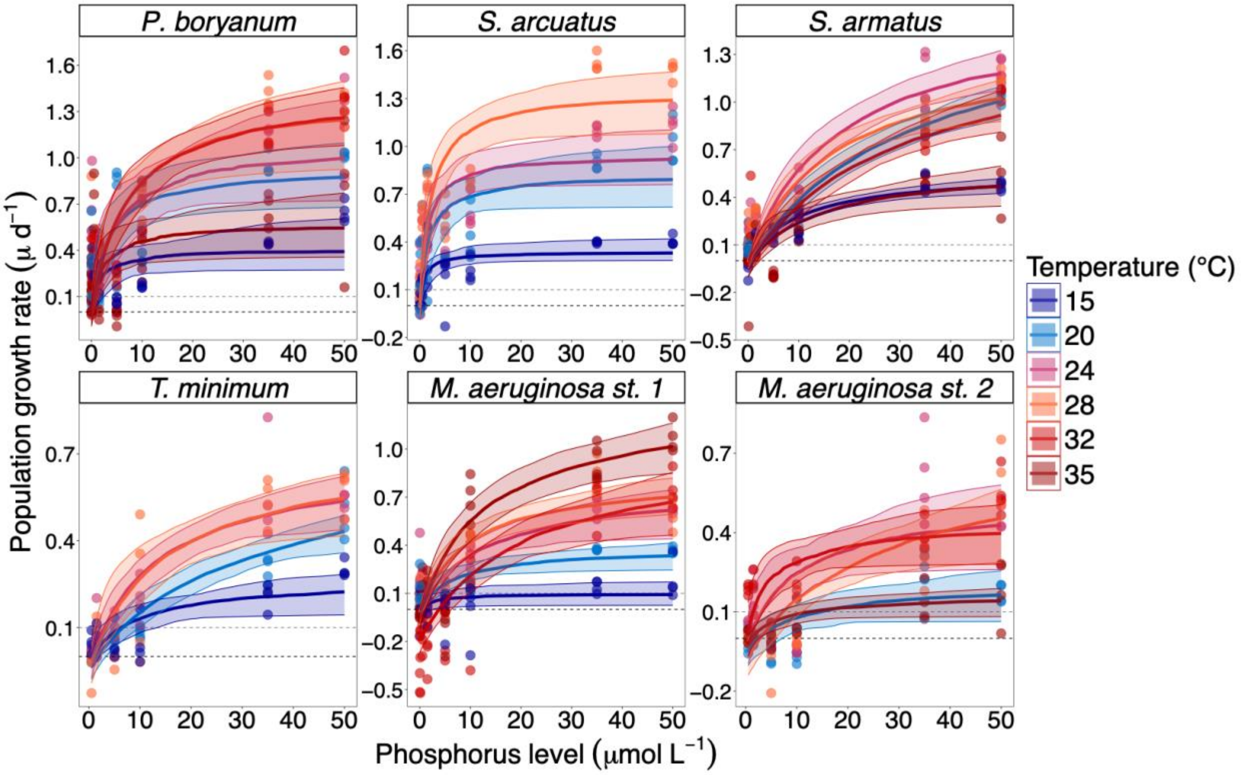
P. boryanum
S. arcuatus
S. armatus
T. minimum
M. aeruginosa st. 1
M. aeruginosa st. 2
Population growth rate (μ d$^{-1}$)
Phosphorus level (μmol L$^{-1}$)
Temperature (°C)
15
20
24
28
32
35

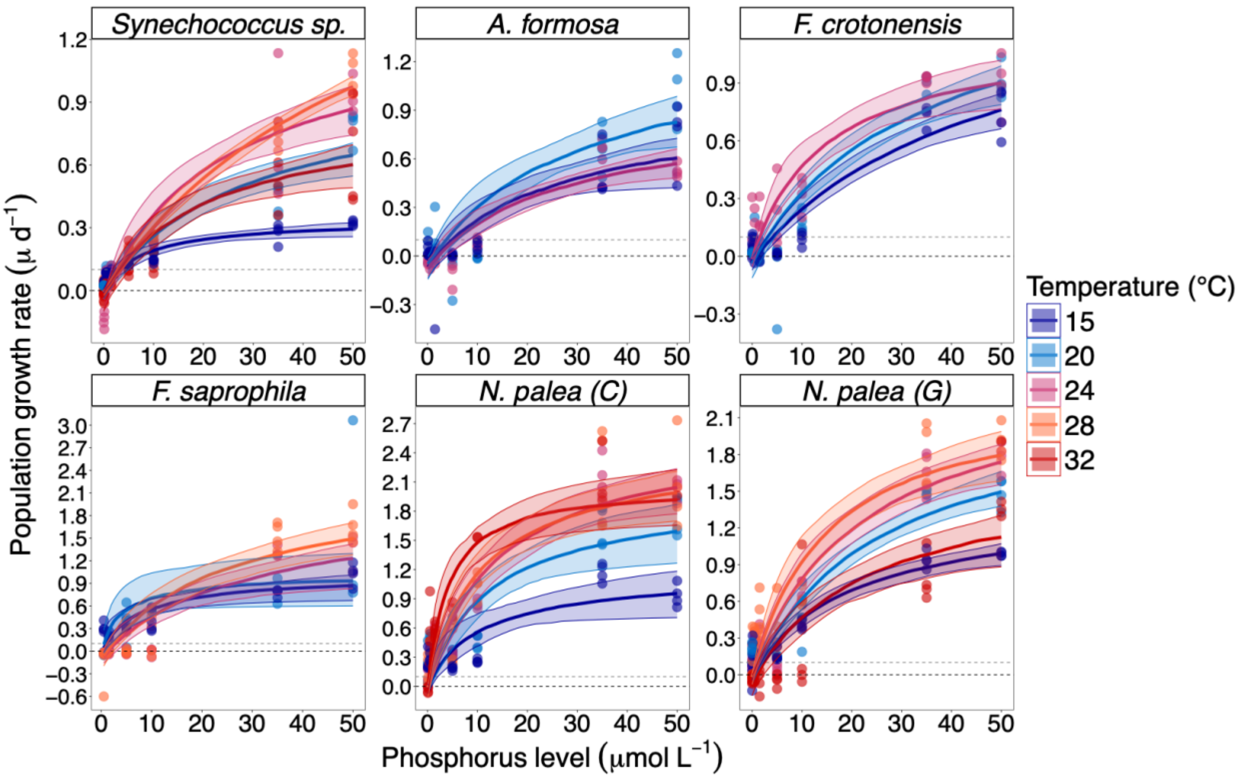

**Figure S6**. Population growth rates (day$^{-1}$) as a function of phosphorus availability for 19 populations of phytoplankton. Monod curves are fitted for each of the six different temperatures separately. Shaded ribbons indicate Bayesian 95% credible intervals. The black dotted line represents the $\mu = 0$ line, and the grey dotted line is the $m_{ext} = 0.1$ line.

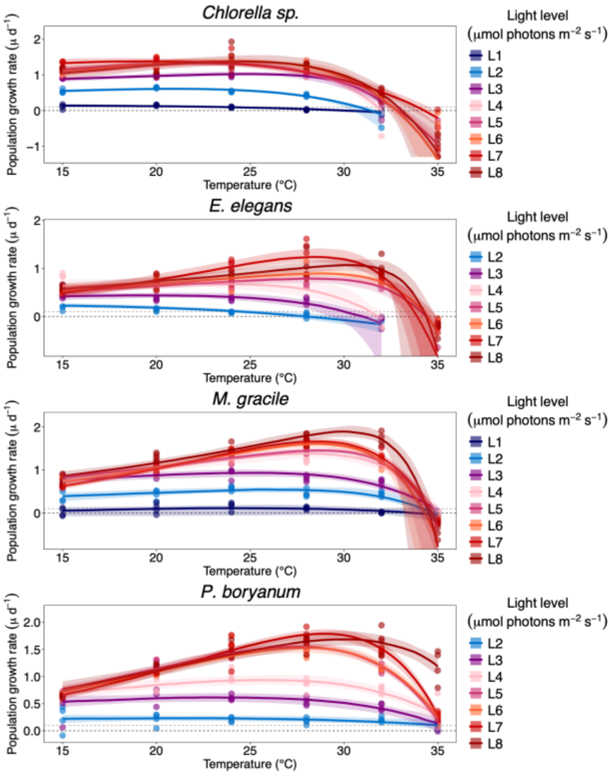
Chlorella sp.
Coelastrum sp.
E. elegans
L. subsalsa
M. gracile
Oocystis sp.
P. boryanum
Population growth rate (µ d⁻¹)
Temperature (°C)
Light level
(µmol photons m⁻² s⁻¹)
L1
L2
L3
L4
L5
L6
L7
L8

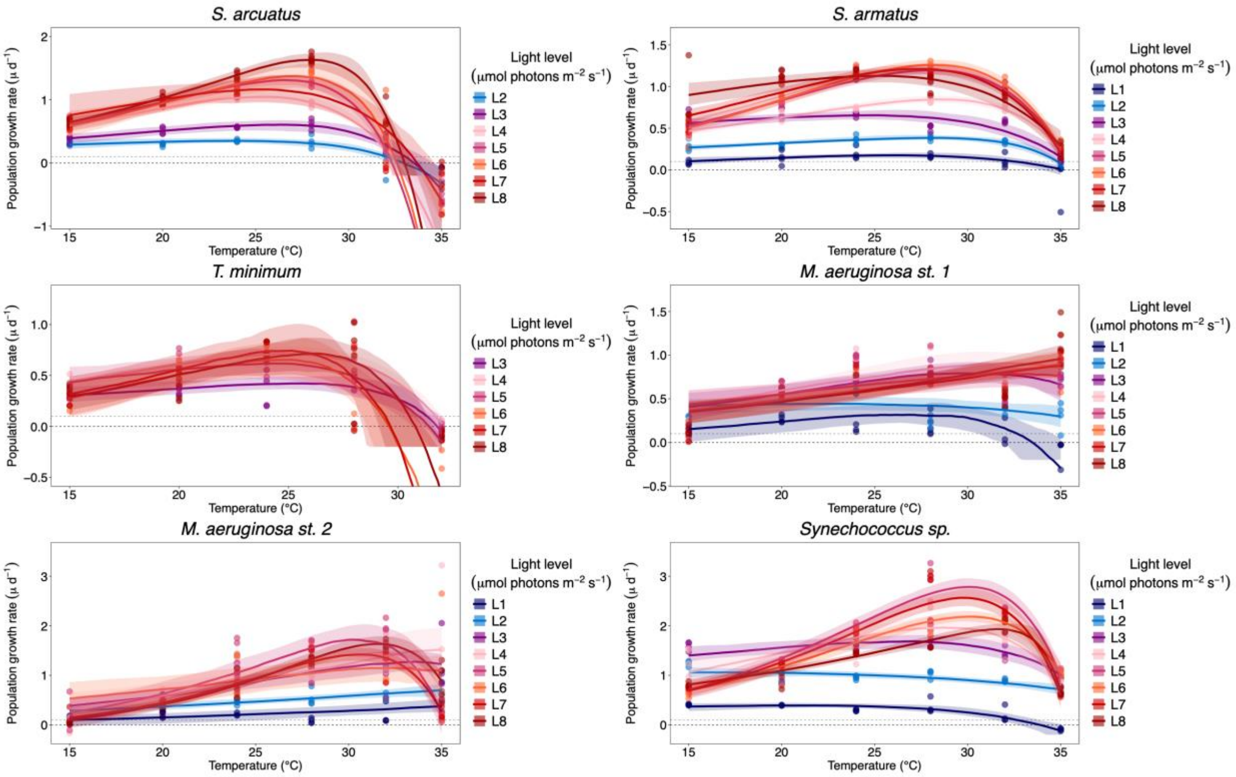
S. arcuatus
S. armatus
T. minimum
M. aeruginosa st. 1
M. aeruginosa st. 2
Synechococcus sp.
Population growth rate (µ d⁻¹)
Temperature (°C)
Light level
(µmol photons m⁻² s⁻¹)
L1
L2
L3
L4
L5
L6
L7
L8

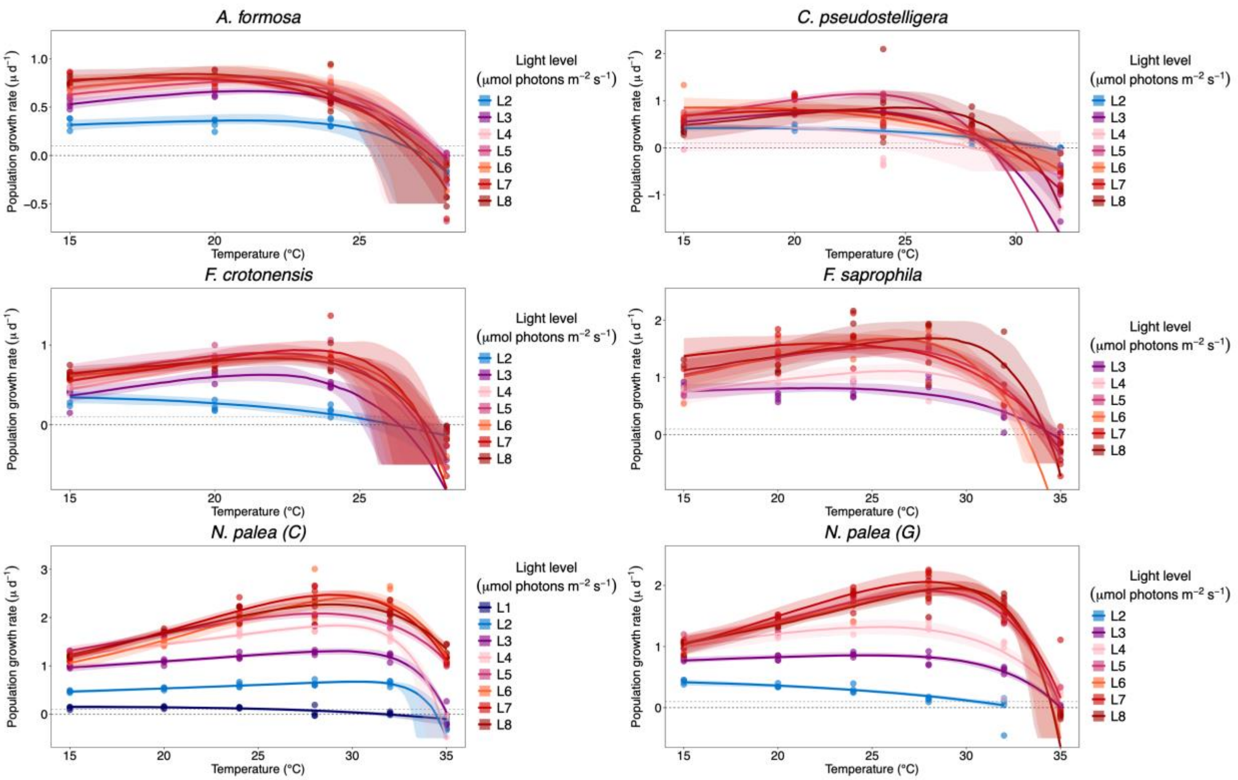

**Figure S7**. Population growth rates (day$^{-1}$) as a function of temperature for 19 populations of phytoplankton. Thermal performance curves are fitted for each of the eight different light levels separately. Shaded ribbons indicate Bayesian 95% credible intervals. The black dotted line represents the $\mu = 0$ line, and the grey dotted line is the $m_{ext} = 0.1$ line.

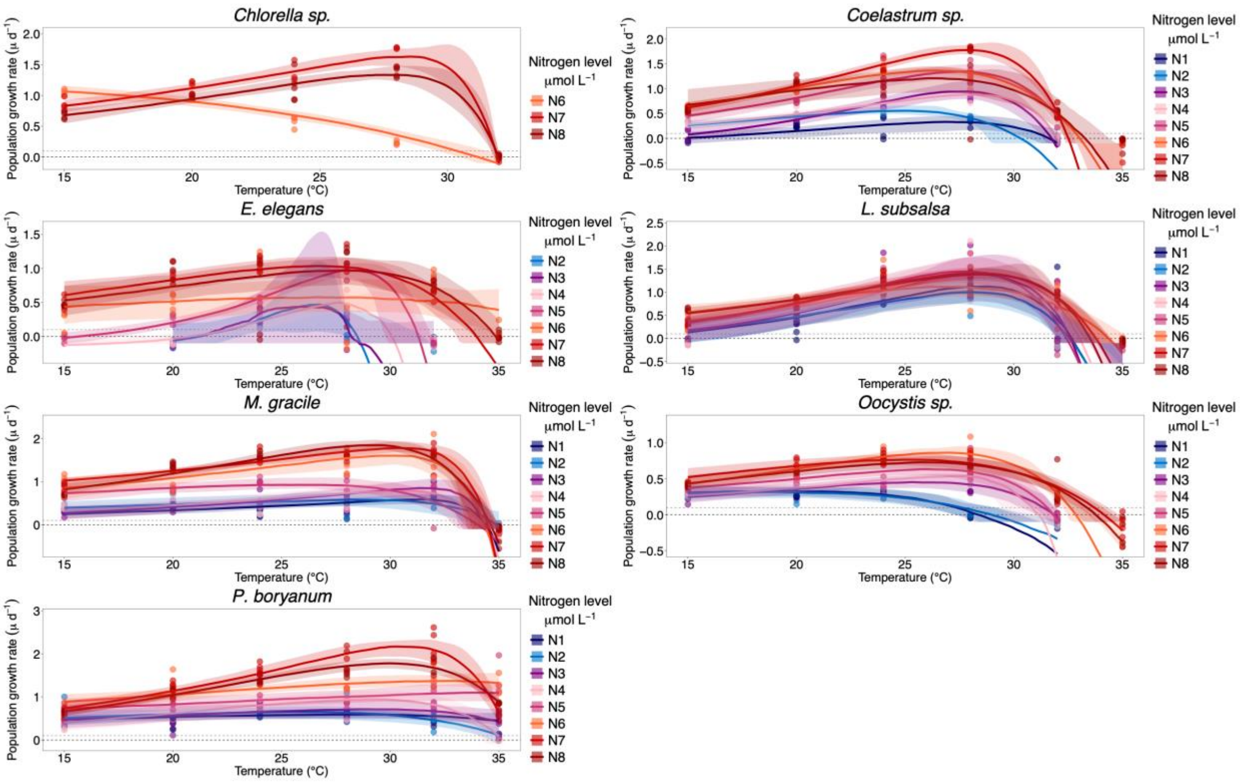
Chlorella sp.
Coelastrum sp.
E. elegans
L. subsalsa
M. gracile
Oocystis sp.
P. boryanum
Population growth rate (µ d⁻¹)
Temperature (°C)
Nitrogen level µmol L⁻¹
N1
N2
N3
N4
N5
N6
N7
N8

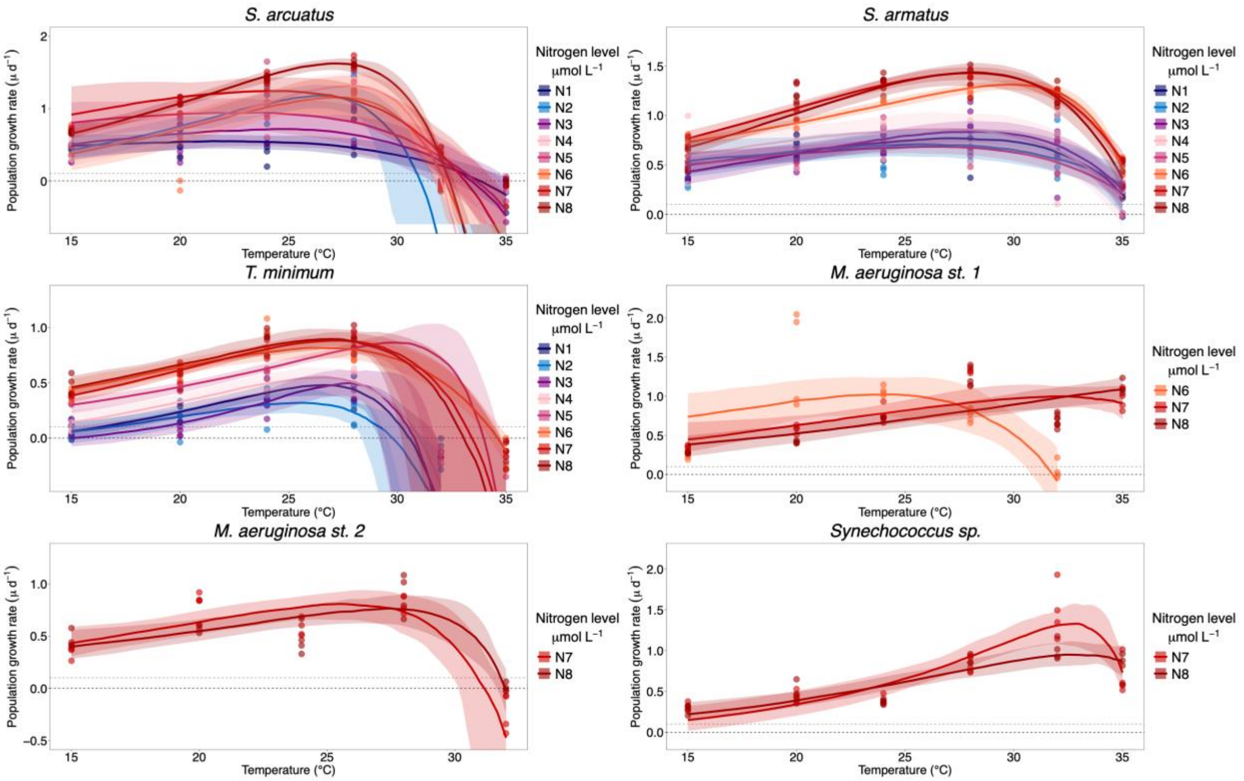
S. arcuatus
S. armatus
T. minimum
M. aeruginosa st. 1
M. aeruginosa st. 2
Synechococcus sp.
Population growth rate (µ d⁻¹)
Temperature (°C)
Nitrogen level µmol L⁻¹
N1
N2
N3
N4
N5
N6
N7
N8

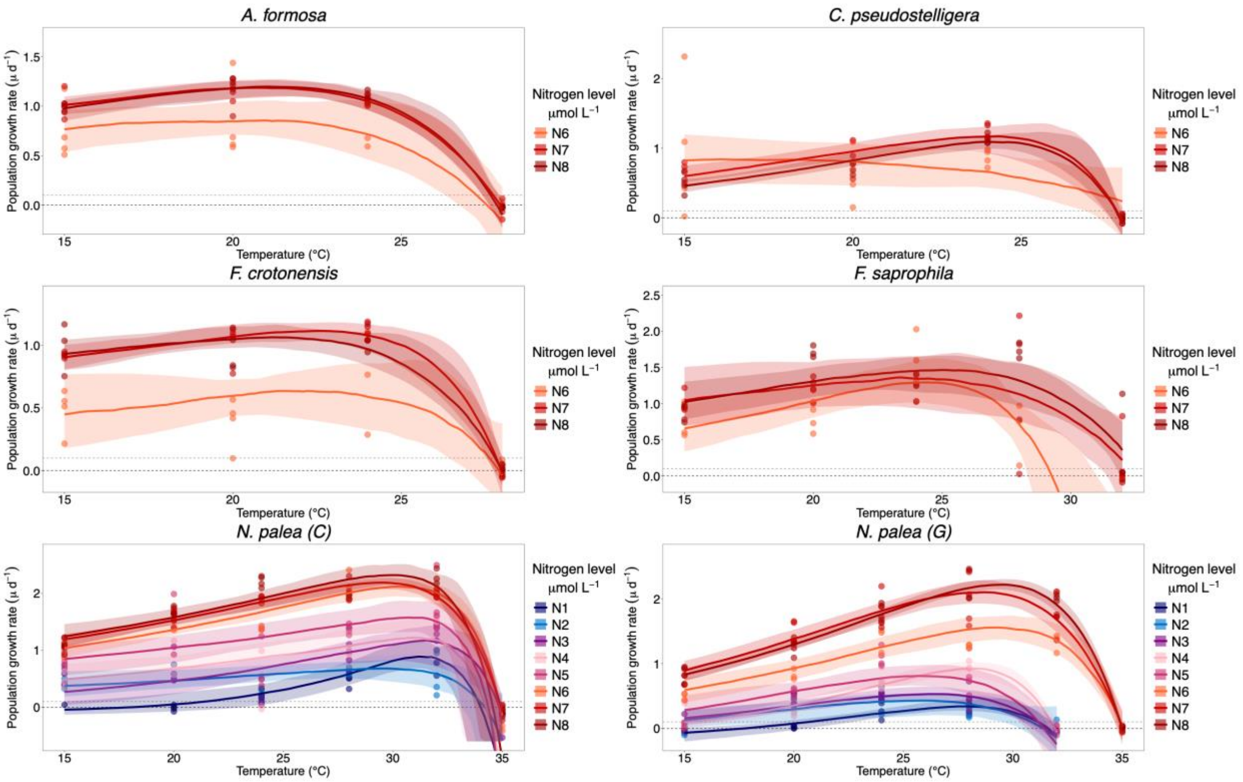

**<u>Figure S8</u>**. Population growth rates (day$^{-1}$) as a function of temperature for 19 populations of phytoplankton. Thermal performance curves are fitted for each of the eight different nitrogen levels separately. Shaded ribbons indicate Bayesian 95% credible intervals. The black dotted line represents the $\mu = 0$ line, and the grey dotted line is the $m_{ext} = 0.1$ line.

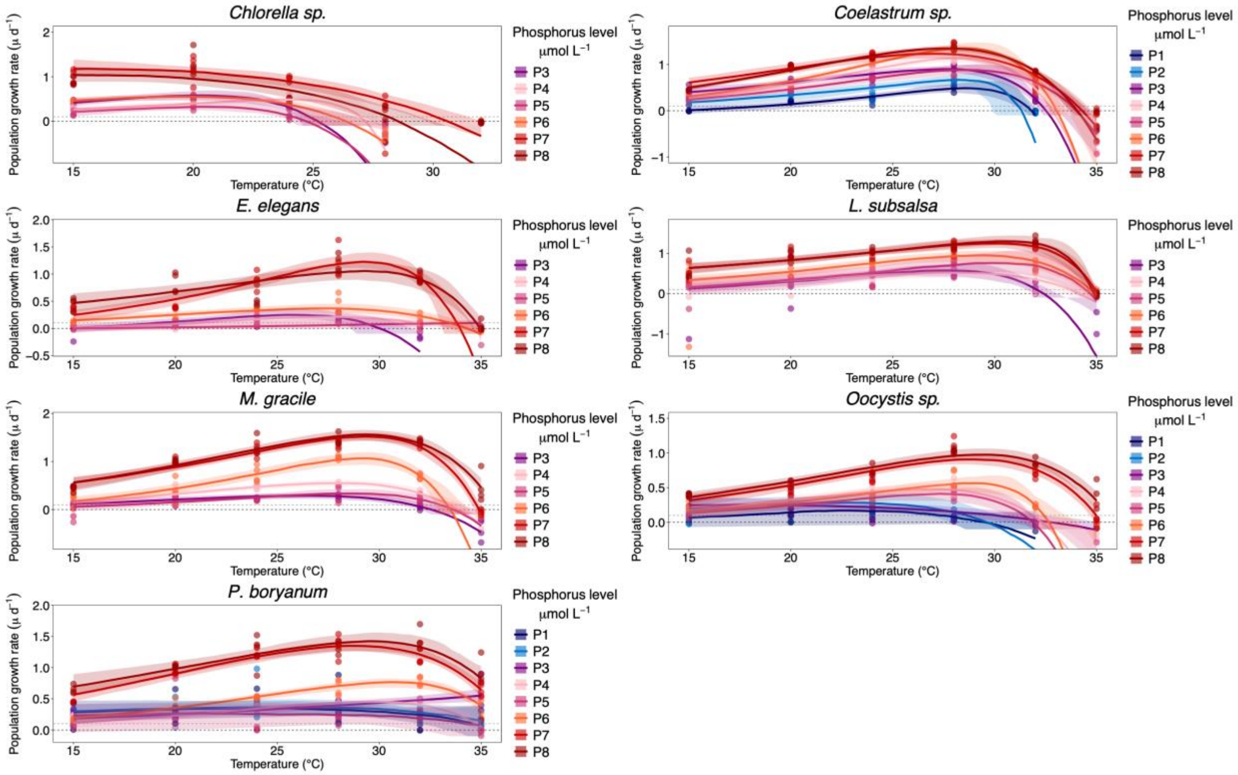
Chlorella sp.
Coelastrum sp.
E. elegans
L. subsalsa
M. gracile
Oocystis sp.
P. boryanum
Population growth rate (μ d−1)
Temperature (°C)
Phosphorus level μmol L−1
P1
P2
P3
P4
P5
P6
P7
P8

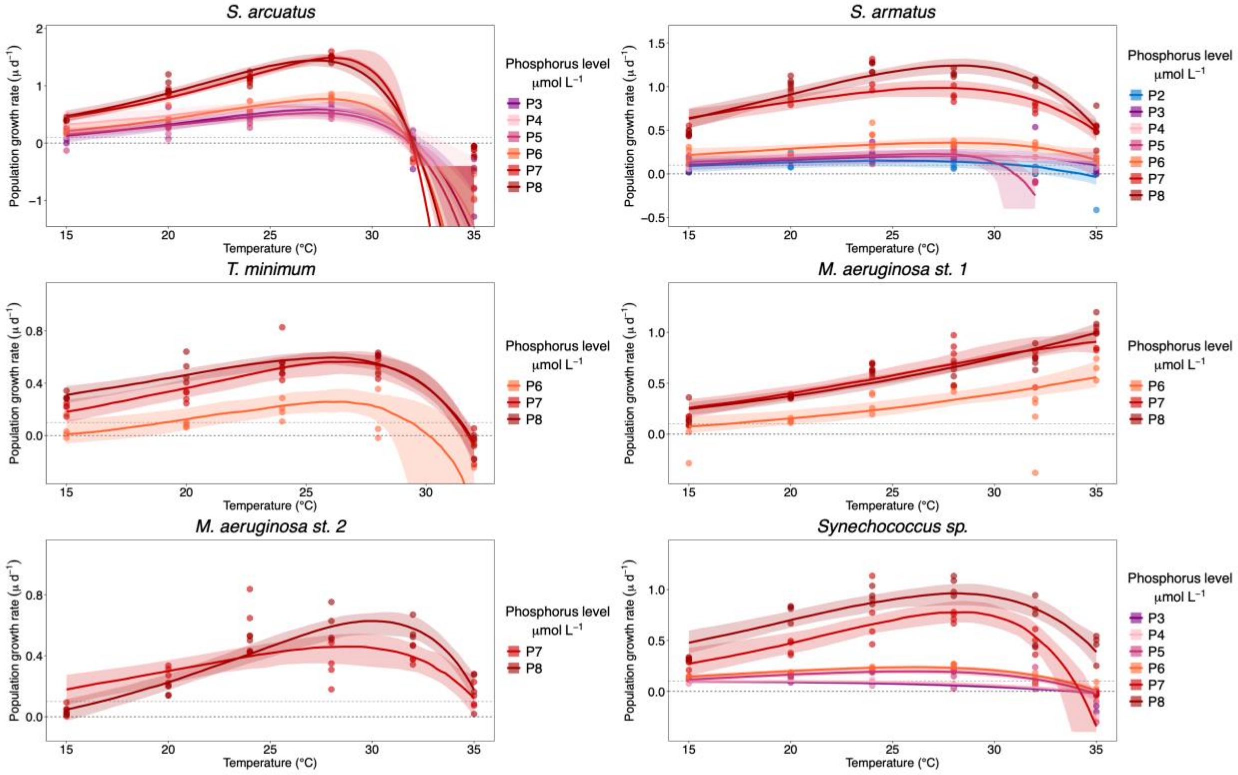
S. arcuatus
S. armatus
T. minimum
M. aeruginosa st. 1
M. aeruginosa st. 2
Synechococcus sp.
Population growth rate (μ d−1)
Temperature (°C)
Phosphorus level μmol L−1
P2
P3
P4
P5
P6
P7
P8

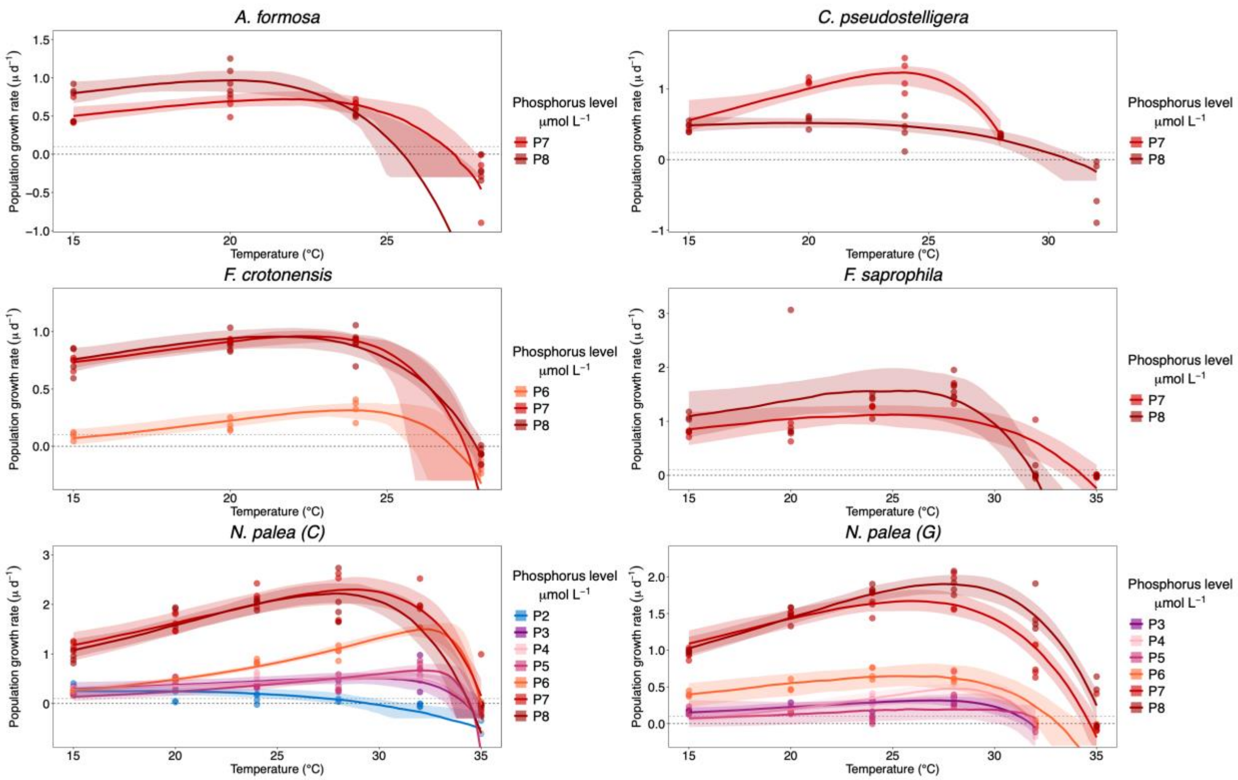

**Figure S9**. Population growth rates (day$^{-1}$) as a function of temperature for 19 populations of phytoplankton. Thermal performance curves are fitted for each of the eight different phosphorus levels separately. Shaded ribbons indicate Bayesian 95% credible intervals. The black dotted line represents the $\mu = 0$ line, and the grey dotted line is the $m_{ext} = 0.1$ line.

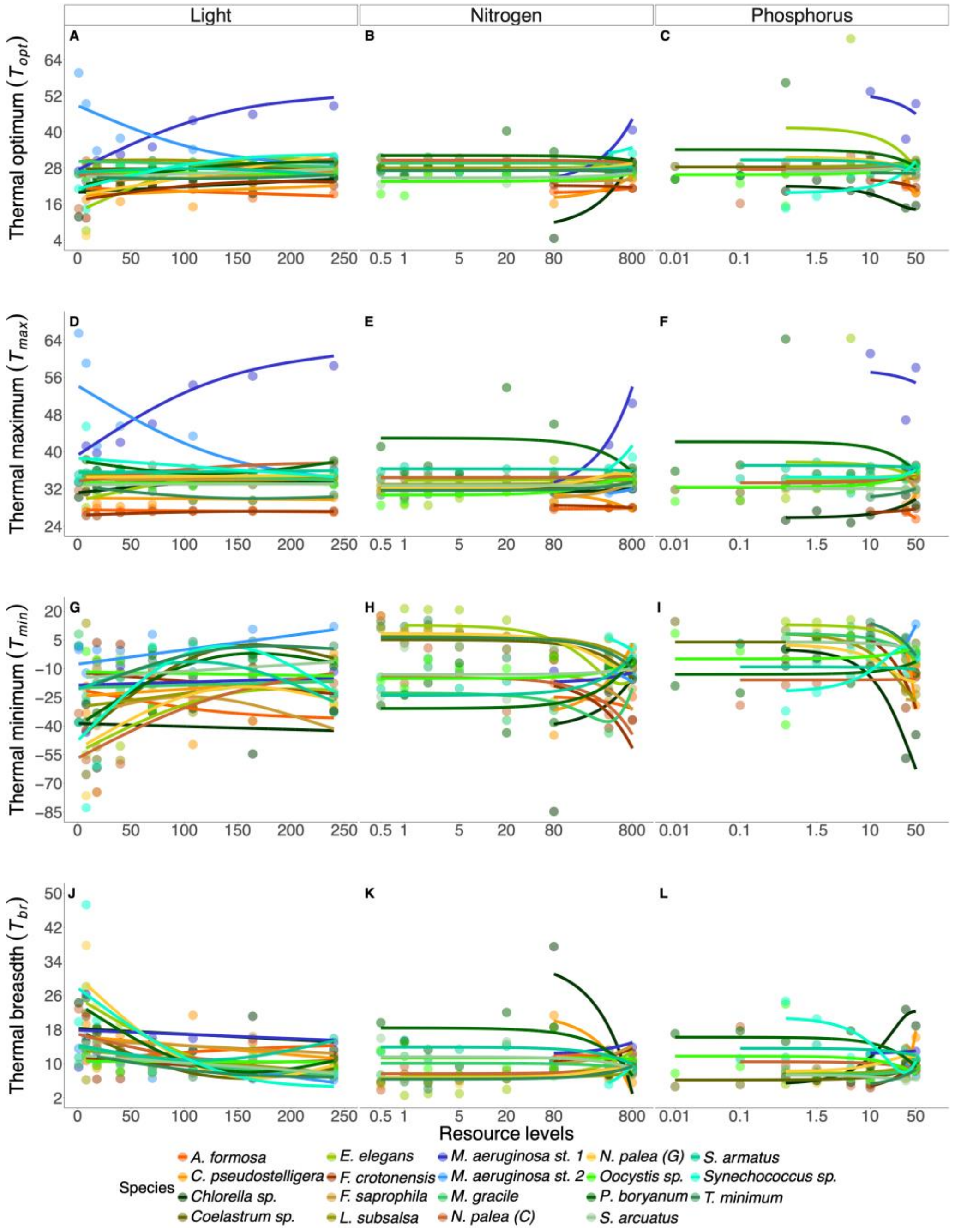

**Figure S10.** Unfiltered thermal performance curve parameters as a function of resource availability for each experiment (light, nitrogen and phosphorus from left to right). Panels in the top row (A-C) represent the thermal optimum (*Topt*); the second row (D-F) the thermal maximum (*Tmax*); the third row (G-I) the thermal minimum (*Tmin*), and the fourth row (J-L) the thermal breadths (*Tbr*). Temperatures are reported in °C. For easier visual representation, the resource axes were logged for the nitrogen and phosphorus panels (B&C, E&F, H&I and K&L), and models were fit to log-transformed values.

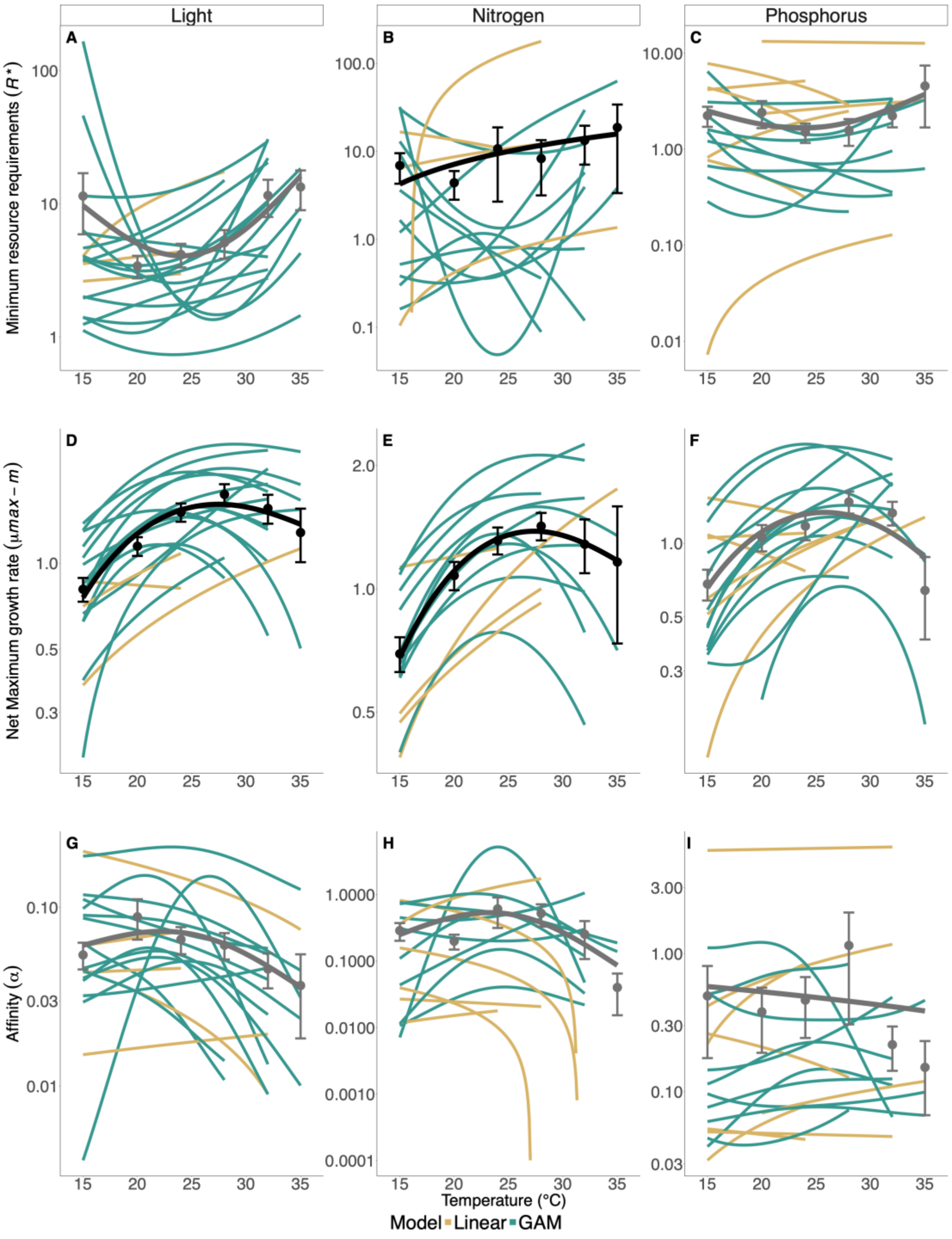

**Figure S11**. EDF categorizations for GAMs from Fig. 2 (main text). Resource-dependent growth parameters plotted as a function of temperature for each experiment (light, nitrogen and phosphorus from left to right). Lines represent model fits for each species. Gold and green colors represent the type of model that best fits the data, either a linear model or a generalized additive model (GAM- see methods).

Points represent the mean parameter across all species ("Mean spp"), and error bars represent the standard error of the mean. Black and grey lines represent the model fits of the mean parameter, black lines represent significant models ($p < 0.05$), and grey lines represent non-significant models. A-C are $R^*$ estimates as a function of temperature, which were calculated with a temperature-independent mortality term ($m_{ext} = 0.1$). The units for $R^*$ for light are μmol photons $m^{-2} \cdot s^{-1}$, while the units of $R^*$ for nitrogen and phosphorus are $\mu mol \cdot L^{-1}$. D-F shows $\mu_{max}$ as a function of temperature. The third row (G-I) shows the $\alpha$ parameter as a function of temperature. For A-C and G-I the y-axis is log-transformed, values are not. Due to this transformation, linear models display a curvature (appear monotonic).

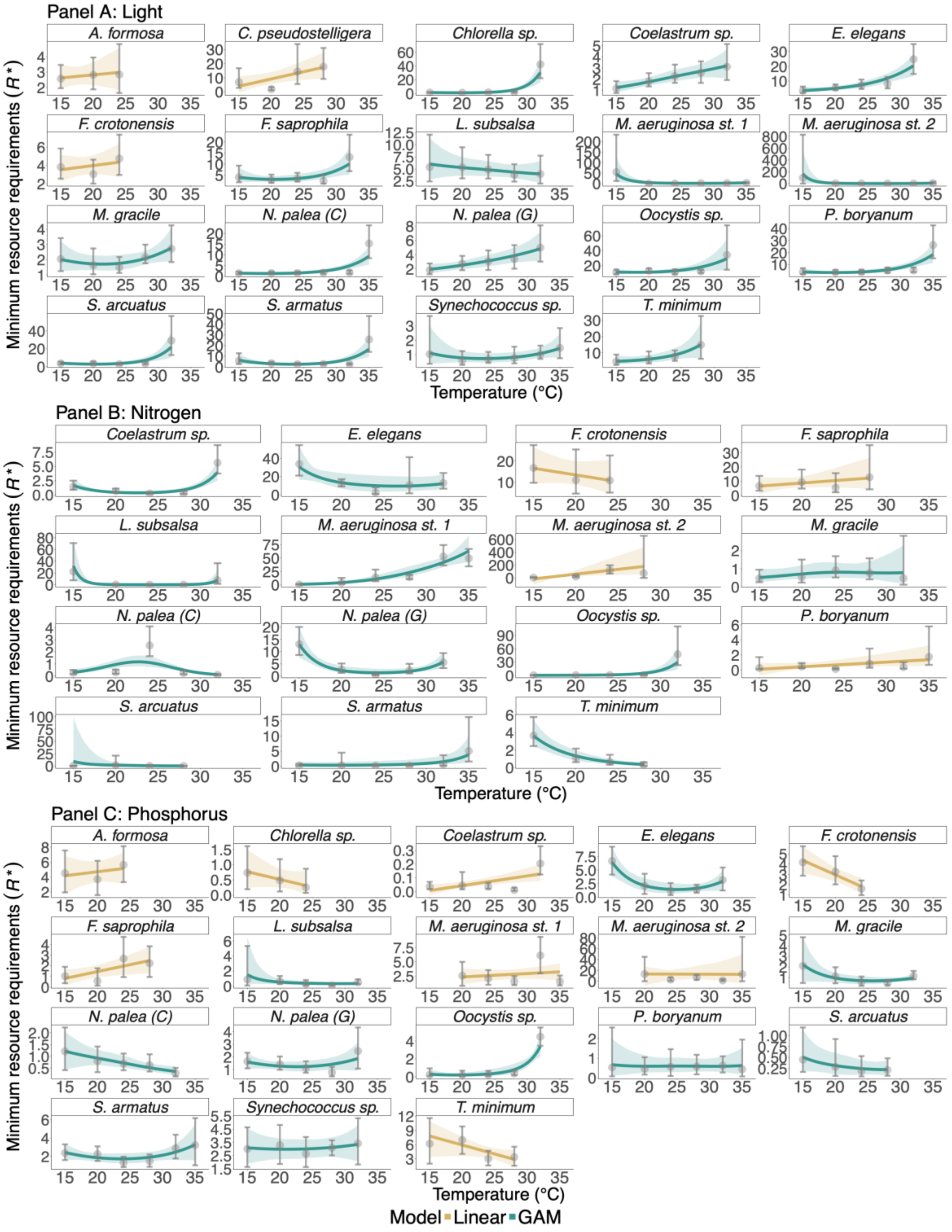

**Figure S12.** EDF categorizations for the minimum resource requirements ($R*$). We have one panel per resource and each panel is faceted per population. Here the y-axis is not logged. Gold and green colors represent the type of model that best fits the data, either a linear model or a generalized additive model

(GAM- see methods). Points and errors bars represent the medians value and their 95% credible interval. The curves are the mean fits from the posteriors bootstrapping procedure and the shaded ribbon their 95% credible interval (see method).

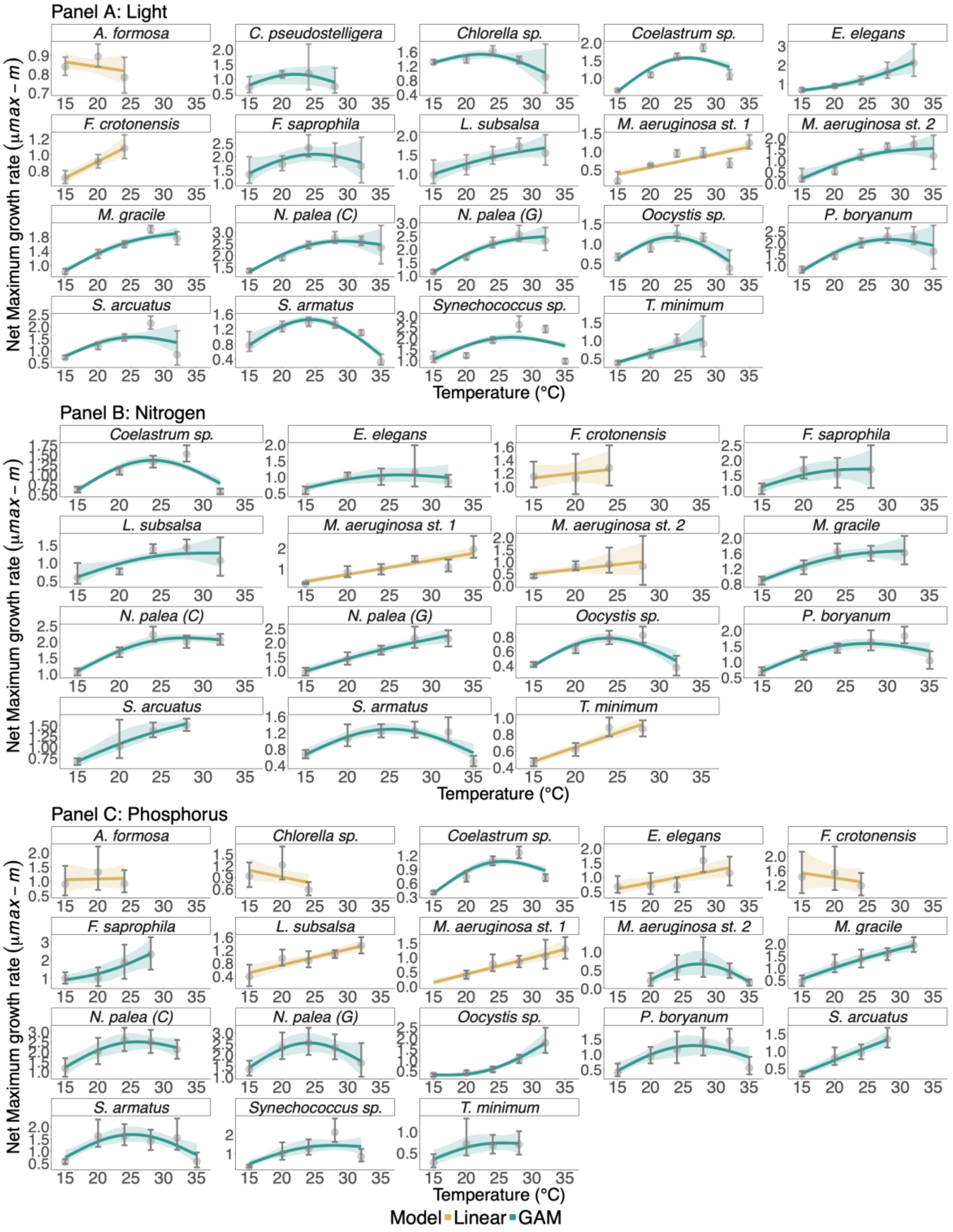

**Figure S13.** EDF categorizations for the net maximum population growth rate (*net-μmax*). We have one panel per resource and each panel is faceted per population. Gold and green colors represent the type of model that best fits the data, either a linear model or a generalized additive model (GAM- see methods).

Points and errors bars represent the medians value and their 95% credible interval. The curves are the mean fits from the posteriors bootstrapping procedure and the shaded ribbon their 95% credible interval (see method).

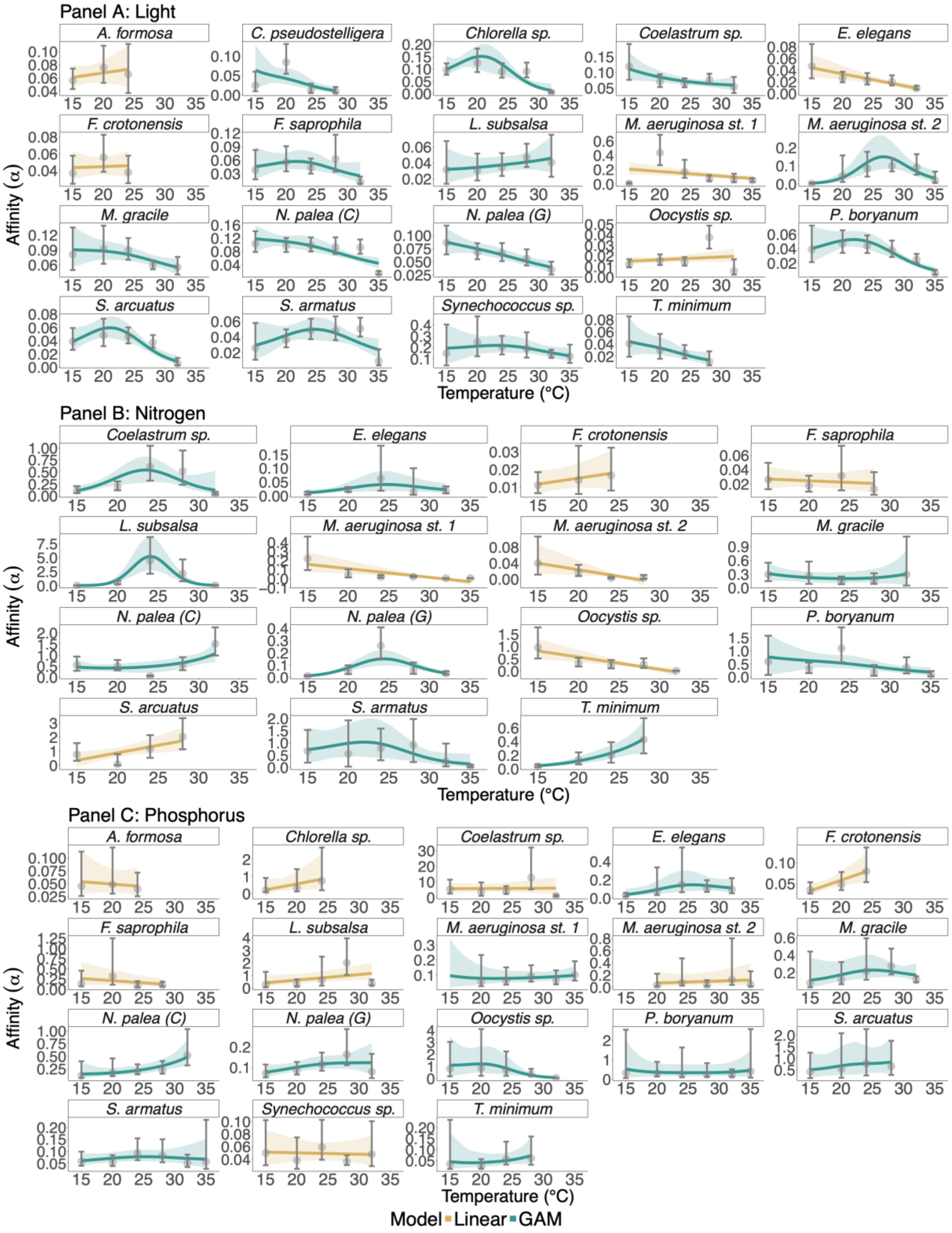

**Figure S14.** EDF categorizations for resource affinity ($\alpha$). We have one panel per resource and each panel is faceted per population. Here the y-axis is not logged. Gold and green colors represent the type of model that best fits the data, either a linear model or a generalized additive model (GAM- see methods). Points and errors bars represent the medians value and their 95% credible interval. The curves are the mean fits from the posteriors bootstrapping procedure and the shaded ribbon their 95% credible interval (see method).

**Table S7.** Summary table of the EDF categorizations for the resource-dependent growth parameters, for the three resources. In the "Data" column we have the parameters; in the "Experiment" column we have the resource type; in "Species" we find the short population name, see full name in the first column of Table S3; in "Model" we have whether a linear model (LM) or a generalized additive model (GAM) fitted the data best; in "M. fit Category" this correspond to the shape attributed to the curves from the posteriors bootstrapping procedure, we can find the following option: U-shape, hump-shape, monotonic or linear increase or decrease; the "Critical point" is the inflexion temperature, which is for a U-shape or hump-shape at which temperature the minimum or maximum was respectively reached; "Slope" if the mean fit curve was determined to be best fitted with a LM, otherwise contains NA; "U_sh" is U-shaped, "H_sh" is hump-shaped, "M_inc" and "M_dec" are respectively monotonic increase and decrease and "LM_dec" and "LM_inc" are linear decrease and increase, these categories show the percentage of the 5,000 posteriors that were sorted in each; and finally the "T. Category" which stands for "True Category", contains the identity of the shape for the parameter having more than 51% of the fits (see methods) in the absent of a majority; the "T. Category" says uncertain.

| Data | Experiment | Species | Model | M. fit Category | Critical point | Slope | U_sh | H_sh | M_inc | M_dec | LM_dec | LM_inc | T. Category |
|---|---|---|---|---|---|---|---|---|---|---|---|---|---|
| Rstar | Light | *A. formosa* | LM | l. decrease | NA | -0.009 | 0 | 0 | 0 | 0 | 36.32 | 63.68 | l. increase |
| Rstar | Light | *C. pseudostelligera* | LM | l. increase | NA | 0.862 | 0 | 0 | 0 | 0 | 1.66 | 98.34 | l. increase |
| Rstar | Light | *Chlorella sp.* | GAM | U-shaped | 19.5 | NA | 93.56 | 0 | 6.44 | 0 | 0 | 0 | U-shaped |
| Rstar | Light | *Coelastrum sp.* | GAM | m. increase | NA | NA | 1.14 | 17.54 | 81.3 | 0.02 | 0 | 0 | m. increase |
| Rstar | Light | *E. elegans* | GAM | m. increase | NA | NA | 11.4 | 0 | 88.6 | 0 | 0 | 0 | m. increase |
| Rstar | Light | *F. crotonensis* | LM | l. decrease | NA | -0.119 | 0 | 0 | 0 | 0 | 26.8 | 73.2 | l. increase |
| Rstar | Light | *F. saprophila* | GAM | U-shaped | 20.5 | NA | 68.34 | 0 | 31.44 | 0.22 | 0 | 0 | U-shaped |
| Rstar | Light | *L. subsalsa* | GAM | m. decrease | NA | NA | 13.02 | 20.74 | 11.6 | 54.64 | 0 | 0 | m. decrease |
| Rstar | Light | *M. aeruginosa st. 1* | GAM | U-shaped | 27.3 | NA | 97.32 | 0.1 | 0.54 | 2.04 | 0 | 0 | U-shaped |
| Rstar | Light | *M. aeruginosa st. 2* | GAM | U-shaped | 27 | NA | 98.76 | 0.02 | 1 | 0.22 | 0 | 0 | U-shaped |
| Rstar | Light | *M. gracile* | GAM | U-shaped | 21.5 | NA | 65.86 | 0.32 | 30.38 | 3.44 | 0 | 0 | U-shaped |
| Rstar | Light | *N. palea (C)* | GAM | U-shaped | 19.5 | NA | 72.26 | 0 | 27.74 | 0 | 0 | 0 | U-shaped |

| | | | | | | | | | | | | | |
|---|---|---|---|---|---|---|---|---|---|---|---|---|---|
| Rstar | Light | *N. palea (G)* | GAM | m. increase | NA | NA | 4.98 | 6.56 | 88.38 | 0.08 | 0 | 0 | m. increase |
| Rstar | Light | *Oocystis sp.* | GAM | U-shaped | 18.1 | NA | 56.78 | 0.56 | 42.06 | 0.6 | 0 | 0 | U-shaped |
| Rstar | Light | *P. boryanum* | GAM | U-shaped | 19.9 | NA | 69.24 | 0 | 30.76 | 0 | 0 | 0 | U-shaped |
| Rstar | Light | *S. arcuatus* | GAM | U-shaped | 20.4 | NA | 90.32 | 0 | 9.68 | 0 | 0 | 0 | U-shaped |
| Rstar | Light | *S. armatus* | GAM | U-shaped | 23.4 | NA | 98.42 | 0 | 1.58 | 0 | 0 | 0 | U-shaped |
| Rstar | Light | *Synechococcus sp.* | GAM | U-shaped | 23.4 | NA | 64.76 | 0.36 | 31.52 | 3.36 | 0 | 0 | U-shaped |
| Rstar | Light | *T. minimum* | GAM | m. increase | NA | NA | 24.34 | 4.08 | 70.96 | 0.62 | 0 | 0 | m. increase |
| Rstar | Nitrogen | *Coelastrum sp.* | GAM | U-shaped | 22.7 | NA | 99.72 | 0 | 0.28 | 0 | 0 | 0 | U-shaped |
| Rstar | Nitrogen | *E. elegans* | GAM | U-shaped | 26.6 | NA | 62.64 | 0 | 1.94 | 35.42 | 0 | 0 | U-shaped |
| Rstar | Nitrogen | *F. crotonensis* | LM | l. decrease | NA | -1.034 | 0 | 0 | 0 | 0 | 83.04 | 16.96 | l. decrease |
| Rstar | Nitrogen | *F. saprophila* | LM | l. increase | NA | 1.95 | 0 | 0 | 0 | 0 | 21.72 | 78.28 | l. increase |
| Rstar | Nitrogen | *L. subsalsa* | GAM | U-shaped | 24 | NA | 100 | 0 | 0 | 0 | 0 | 0 | U-shaped |
| Rstar | Nitrogen | *M. aeruginosa st. 1* | GAM | m. increase | NA | NA | 0 | 1.28 | 98.72 | 0 | 0 | 0 | m. increase |
| Rstar | Nitrogen | *M. aeruginosa st. 2* | LM | l. increase | NA | 3.825 | 0 | 0 | 0 | 0 | 0 | 100 | l. increase |
| Rstar | Nitrogen | *M. gracile* | GAM | hump-shaped | 24.6 | NA | 2.52 | 47.82 | 39.68 | 9.98 | 0 | 0 | hump-shaped |
| Rstar | Nitrogen | *N. palea (C)* | GAM | hump-shaped | 22.7 | NA | 0 | 99.98 | 0 | 0.02 | 0 | 0 | hump-shaped |
| Rstar | Nitrogen | *N. palea (G)* | GAM | U-shaped | 24.4 | NA | 98.54 | 0 | 0.02 | 1.44 | 0 | 0 | U-shaped |
| Rstar | Nitrogen | *Oocystis sp.* | GAM | m. increase | NA | NA | 8.86 | 0 | 91.14 | 0 | 0 | 0 | m. increase |
| Rstar | Nitrogen | *P. boryanum* | LM | l. increase | NA | 0.053 | 0 | 0 | 0 | 0 | 3.18 | 96.82 | l. increase |
| Rstar | Nitrogen | *S. arcuatus* | GAM | m. decrease | NA | NA | 0.1 | 59.36 | 0.04 | 40.5 | 0 | 0 | hump-shaped |
| Rstar | Nitrogen | *S. armatus* | GAM | U-shaped | 19.1 | NA | 64.64 | 0 | 34.28 | 1.08 | 0 | 0 | U-shaped |
| Rstar | Nitrogen | *T. minimum* | GAM | m. decrease | NA | NA | 3.66 | 0.2 | 0 | 96.14 | 0 | 0 | m. decrease |
| Rstar | Phosphorus | *A. formosa* | LM | l. increase | NA | 0.243 | 0 | 0 | 0 | 0 | 29.12 | 70.88 | l. increase |
| Rstar | Phosphorus | *Chlorella sp.* | LM | l. decrease | NA | -0.03 | 0 | 0 | 0 | 0 | 86.78 | 13.22 | l. decrease |
| Rstar | Phosphorus | *Coelastrum sp.* | LM | l. increase | NA | 0.003 | 0 | 0 | 0 | 0 | 0.28 | 99.72 | l. increase |
| Rstar | Phosphorus | *E. elegans* | GAM | U-shaped | 25 | NA | 92.9 | 0 | 0.04 | 7.06 | 0 | 0 | U-shaped |
| Rstar | Phosphorus | *F. crotonensis* | LM | l. decrease | NA | -0.223 | 0 | 0 | 0 | 0 | 99.9 | 0.1 | l. decrease |
| Rstar | Phosphorus | *F. saprophila* | LM | l. increase | NA | 0.129 | 0 | 0 | 0 | 0 | 1.46 | 98.54 | l. increase |
| Rstar | Phosphorus | *L. subsalsa* | GAM | U-shaped | 29.4 | NA | 34.16 | 0.02 | 7.16 | 58.66 | 0 | 0 | m. decrease |

| | | | | | | | | | | | | | |
|---|---|---|---|---|---|---|---|---|---|---|---|---|---|
| Rstar | Phosphorus | *M. aeruginosa st. 1* | LM | l. increase | NA | 0.017 | 0 | 0 | 0 | 0 | 21.62 | 78.38 | l. increase |
| Rstar | Phosphorus | *M. aeruginosa st. 2* | LM | l. decrease | NA | -1.242 | 0 | 0 | 0 | 0 | 55.68 | 44.32 | l. decrease |
| Rstar | Phosphorus | *M. gracile* | GAM | U-shaped | 25.8 | NA | 74.42 | 0.02 | 7.18 | 18.38 | 0 | 0 | U-shaped |
| Rstar | Phosphorus | *N. palea (C)* | GAM | m. decrease | NA | NA | 0.82 | 19.4 | 0.3 | 79.48 | 0 | 0 | m. decrease |
| Rstar | Phosphorus | *N. palea (G)* | GAM | U-shaped | 22.7 | NA | 67.76 | 0.04 | 19.2 | 13 | 0 | 0 | U-shaped |
| Rstar | Phosphorus | *Oocystis sp.* | GAM | U-shaped | 19.2 | NA | 67.22 | 0 | 32.78 | 0 | 0 | 0 | U-shaped |
| Rstar | Phosphorus | *P. boryanum* | GAM | U-shaped | 20.7 | NA | 17.24 | 23.94 | 28.58 | 30.24 | 0 | 0 | m. decrease |
| Rstar | Phosphorus | *S. arcuatus* | GAM | U-shaped | 27.7 | NA | 26.4 | 7.76 | 5.94 | 59.9 | 0 | 0 | m. decrease |
| Rstar | Phosphorus | *S. armatus* | GAM | U-shaped | 23.8 | NA | 76.56 | 0 | 13.2 | 10.24 | 0 | 0 | U-shaped |
| Rstar | Phosphorus | *Synechococcus sp.* | GAM | U-shaped | 21 | NA | 27.66 | 9.68 | 37.22 | 25.44 | 0 | 0 | m. increase |
| Rstar | Phosphorus | *T. minimum* | LM | l. decrease | NA | -0.208 | 0 | 0 | 0 | 0 | 93.92 | 6.08 | l. decrease |
| Net mumax | Light | *A. formosa* | LM | l. increase | NA | 0.001 | 0 | 0 | 0 | 0 | 81.98 | 18.02 | l. decrease |
| Net mumax | Light | *C. pseudostelligera* | GAM | hump-shaped | 21.9 | NA | 0.12 | 79.54 | 13.26 | 7.08 | 0 | 0 | hump-shaped |
| Net mumax | Light | *Chlorella sp.* | GAM | hump-shaped | 22.2 | NA | 0.04 | 90.1 | 9.52 | 0.34 | 0 | 0 | hump-shaped |
| Net mumax | Light | *Coelastrum sp.* | GAM | hump-shaped | 25.7 | NA | 0 | 100 | 0 | 0 | 0 | 0 | hump-shaped |
| Net mumax | Light | *E. elegans* | GAM | m. increase | NA | NA | 7.92 | 0.1 | 91.98 | 0 | 0 | 0 | m. increase |
| Net mumax | Light | *F. crotonensis* | LM | l. increase | NA | 0.046 | 0 | 0 | 0 | 0 | 0 | 100 | l. increase |
| Net mumax | Light | *F. saprophila* | GAM | hump-shaped | 25 | NA | 0.26 | 77.22 | 20.08 | 2.44 | 0 | 0 | hump-shaped |
| Net mumax | Light | *L. subsalsa* | GAM | m. increase | NA | NA | 0.58 | 18 | 81.24 | 0.18 | 0 | 0 | m. increase |
| Net mumax | Light | *M. aeruginosa st. 1* | LM | l. increase | NA | 0.035 | 0 | 0 | 0 | 0 | 0.04 | 99.96 | l. increase |
| Net mumax | Light | *M. aeruginosa st. 2* | GAM | m. increase | NA | NA | 0 | 42.56 | 57.44 | 0 | 0 | 0 | m. increase |
| Net mumax | Light | *M. gracile* | GAM | m. increase | NA | NA | 0 | 14.78 | 85.22 | 0 | 0 | 0 | m. increase |
| Net mumax | Light | *N. palea (C)* | GAM | hump-shaped | 29 | NA | 0 | 82.06 | 17.94 | 0 | 0 | 0 | hump-shaped |
| Net mumax | Light | *N. palea (G)* | GAM | m. increase | NA | NA | 0 | 33 | 67 | 0 | 0 | 0 | m. increase |
| Net mumax | Light | *Oocystis sp.* | GAM | hump-shaped | 23.4 | NA | 0 | 99.52 | 0.48 | 0 | 0 | 0 | hump-shaped |
| Net mumax | Light | *P. boryanum* | GAM | hump-shaped | 28.2 | NA | 0 | 86.78 | 13.22 | 0 | 0 | 0 | hump-shaped |
| Net mumax | Light | *S. arcuatus* | GAM | hump-shaped | 25.6 | NA | 0 | 85.56 | 14.44 | 0 | 0 | 0 | hump-shaped |
| Net mumax | Light | *S. armatus* | GAM | hump-shaped | 24.3 | NA | 0 | 99.72 | 0 | 0.28 | 0 | 0 | hump-shaped |
| Net mumax | Light | *Synechococcus sp.* | GAM | hump-shaped | 26.9 | NA | 0 | 99.62 | 0.3 | 0.08 | 0 | 0 | hump-shaped |

| | | | | | | | | | | | | | |
|---|---|---|---|---|---|---|---|---|---|---|---|---|---|
| Net mumax | Light | *T. minimum* | GAM | m. increase | NA | NA | 0.36 | 12.34 | 87.3 | 0 | 0 | 0 | m. increase |
| Net mumax | Nitrogen | *Coelastrum sp.* | GAM | hump-shaped | 24 | NA | 0 | 100 | 0 | 0 | 0 | 0 | hump-shaped |
| Net mumax | Nitrogen | *E. elegans* | GAM | hump-shaped | 26.2 | NA | 0 | 72.14 | 27.6 | 0.26 | 0 | 0 | hump-shaped |
| Net mumax | Nitrogen | *F. crotonensis* | LM | l. decrease | NA | -0.038 | 0 | 0 | 0 | 0 | 22.58 | 77.42 | l. increase |
| Net mumax | Nitrogen | *F. saprophila* | GAM | m. increase | NA | NA | 0.02 | 40.24 | 59.46 | 0.28 | 0 | 0 | m. increase |
| Net mumax | Nitrogen | *L. subsalsa* | GAM | hump-shaped | 30.9 | NA | 0.1 | 50.3 | 49.12 | 0.48 | 0 | 0 | hump-shaped |
| Net mumax | Nitrogen | *M. aeruginosa st. 1* | LM | l. increase | NA | 0.062 | 0 | 0 | 0 | 0 | 0 | 100 | l. increase |
| Net mumax | Nitrogen | *M. aeruginosa st. 2* | LM | l. increase | NA | 0.089 | 0 | 0 | 0 | 0 | 14.08 | 85.92 | l. increase |
| Net mumax | Nitrogen | *M. gracile* | GAM | m. increase | NA | NA | 0 | 44.7 | 55.3 | 0 | 0 | 0 | m. increase |
| Net mumax | Nitrogen | *N. palea (C)* | GAM | hump-shaped | 27.9 | NA | 0 | 85.1 | 14.9 | 0 | 0 | 0 | hump-shaped |
| Net mumax | Nitrogen | *N. palea (G)* | GAM | m. increase | NA | NA | 0 | 1.82 | 98.18 | 0 | 0 | 0 | m. increase |
| Net mumax | Nitrogen | *Oocystis sp.* | GAM | hump-shaped | 23.9 | NA | 0 | 99.9 | 0.1 | 0 | 0 | 0 | hump-shaped |
| Net mumax | Nitrogen | *P. boryanum* | GAM | hump-shaped | 27.6 | NA | 0 | 96.86 | 3.14 | 0 | 0 | 0 | hump-shaped |
| Net mumax | Nitrogen | *S. arcuatus* | GAM | m. increase | NA | NA | 0 | 9.58 | 90.42 | 0 | 0 | 0 | m. increase |
| Net mumax | Nitrogen | *S. armatus* | GAM | hump-shaped | 25.1 | NA | 0 | 99.86 | 0.14 | 0 | 0 | 0 | hump-shaped |
| Net mumax | Nitrogen | *T. minimum* | LM | l. increase | NA | 0.026 | 0 | 0 | 0 | 0 | 0 | 100 | l. increase |
| Net mumax | Phosphorus | *A. formosa* | LM | l. increase | NA | 0.012 | 0 | 0 | 0 | 0 | 42.2 | 57.8 | l. increase |
| Net mumax | Phosphorus | *Chlorella sp.* | LM | l. decrease | NA | -0.036 | 0 | 0 | 0 | 0 | 96.22 | 3.78 | l. decrease |
| Net mumax | Phosphorus | *Coelastrum sp.* | GAM | hump-shaped | 25.6 | NA | 0 | 99.92 | 0.08 | 0 | 0 | 0 | hump-shaped |
| Net mumax | Phosphorus | *E. elegans* | LM | l. increase | NA | 0.039 | 0 | 0 | 0 | 0 | 0.6 | 99.4 | l. increase |
| Net mumax | Phosphorus | *F. crotonensis* | LM | l. decrease | NA | -0.075 | 0 | 0 | 0 | 0 | 76.64 | 23.36 | l. decrease |
| Net mumax | Phosphorus | *F. saprophila* | GAM | m. increase | NA | NA | 22.82 | 0.86 | 76.32 | 0 | 0 | 0 | m. increase |
| Net mumax | Phosphorus | *L. subsalsa* | LM | l. increase | NA | 0.028 | 0 | 0 | 0 | 0 | 0.04 | 99.96 | l. increase |
| Net mumax | Phosphorus | *M. aeruginosa st. 1* | LM | l. increase | NA | 0.066 | 0 | 0 | 0 | 0 | 0 | 100 | l. increase |
| Net mumax | Phosphorus | *M. aeruginosa st. 2* | GAM | hump-shaped | 27.4 | NA | 0 | 97.4 | 0.78 | 1.82 | 0 | 0 | hump-shaped |
| Net mumax | Phosphorus | *M. gracile* | GAM | m. increase | NA | NA | 0 | 2.66 | 97.34 | 0 | 0 | 0 | m. increase |
| Net mumax | Phosphorus | *N. palea (C)* | GAM | hump-shaped | 26 | NA | 0 | 84.96 | 15.02 | 0.02 | 0 | 0 | hump-shaped |
| Net mumax | Phosphorus | *N. palea (G)* | GAM | hump-shaped | 24.1 | NA | 0 | 95.66 | 4.26 | 0.08 | 0 | 0 | hump-shaped |
| Net mumax | Phosphorus | *Oocystis sp.* | GAM | U-shaped | 17 | NA | 62.1 | 0 | 37.9 | 0 | 0 | 0 | U-shaped |

| | | | | | | | | | | | | | |
|---|---|---|---|---|---|---|---|---|---|---|---|---|---|
| Net mumax | Phosphorus | *P. boryanum* | GAM | hump-shaped | 26.5 | NA | 0 | 93 | 6.98 | 0.02 | 0 | 0 | hump-shaped |
| Net mumax | Phosphorus | *S. arcuatus* | GAM | m. increase | NA | NA | 0 | 1.08 | 98.92 | 0 | 0 | 0 | m. increase |
| Net mumax | Phosphorus | *S. armatus* | GAM | hump-shaped | 25.2 | NA | 0 | 99.14 | 0.86 | 0 | 0 | 0 | hump-shaped |
| Net mumax | Phosphorus | *Synechococcus sp.* | GAM | hump-shaped | 27.8 | NA | 0 | 62.1 | 37.9 | 0 | 0 | 0 | hump-shaped |
| Net mumax | Phosphorus | *T. minimum* | GAM | hump-shaped | 25.5 | NA | 0.36 | 50.16 | 49.32 | 0.16 | 0 | 0 | hump-shaped |
| Affinity | Light | *A. formosa* | LM | l. decrease | NA | -0.0018 | 0 | 0 | 0 | 0 | 27.92 | 72.08 | l. increase |
| Affinity | Light | *C. pseudostelligera* | GAM | m. decrease | NA | NA | 0 | 31.46 | 0.08 | 68.46 | 0 | 0 | m. decrease |
| Affinity | Light | *Chlorella sp.* | GAM | hump-shaped | 20.7 | NA | 0 | 93.94 | 0 | 6.06 | 0 | 0 | hump-shaped |
| Affinity | Light | *Coelastrum sp.* | GAM | m. decrease | NA | NA | 21.42 | 0.7 | 0.52 | 77.36 | 0 | 0 | m. decrease |
| Affinity | Light | *E. elegans* | LM | l. decrease | NA | -0.002 | 0 | 0 | 0 | 0 | 100 | 0 | l. decrease |
| Affinity | Light | *F. crotonensis* | LM | l. increase | NA | 0.0017 | 0 | 0 | 0 | 0 | 41.14 | 58.86 | l. increase |
| Affinity | Light | *F. saprophila* | GAM | hump-shaped | 21.7 | NA | 0 | 64.2 | 5.28 | 30.52 | 0 | 0 | hump-shaped |
| Affinity | Light | *L. subsalsa* | GAM | m. increase | NA | NA | 15.56 | 13.82 | 58.54 | 12.08 | 0 | 0 | m. increase |
| Affinity | Light | *M. aeruginosa st. 1* | LM | l. decrease | NA | -0.0095 | 0 | 0 | 0 | 0 | 99.88 | 0.12 | l. decrease |
| Affinity | Light | *M. aeruginosa st. 2* | GAM | hump-shaped | 26.7 | NA | 0 | 99.38 | 0.36 | 0.26 | 0 | 0 | hump-shaped |
| Affinity | Light | *M. gracile* | GAM | m. decrease | NA | NA | 0.3 | 44.96 | 0.64 | 54.1 | 0 | 0 | m. decrease |
| Affinity | Light | *N. palea (C)* | GAM | m. decrease | NA | NA | 0 | 40.6 | 0 | 59.4 | 0 | 0 | m. decrease |
| Affinity | Light | *N. palea (G)* | GAM | m. decrease | NA | NA | 0.14 | 18.4 | 0 | 81.46 | 0 | 0 | m. decrease |
| Affinity | Light | *Oocystis sp.* | LM | l. decrease | NA | -3.00E-04 | 0 | 0 | 0 | 0 | 12.66 | 87.34 | l. increase |
| Affinity | Light | *P. boryanum* | GAM | hump-shaped | 21.8 | NA | 0 | 90.24 | 0 | 9.76 | 0 | 0 | hump-shaped |
| Affinity | Light | *S. arcuatus* | GAM | hump-shaped | 21.1 | NA | 0 | 97.14 | 0 | 2.86 | 0 | 0 | hump-shaped |
| Affinity | Light | *S. armatus* | GAM | hump-shaped | 24.6 | NA | 0 | 82 | 1.46 | 16.54 | 0 | 0 | hump-shaped |
| Affinity | Light | *Synechococcus sp.* | GAM | hump-shaped | 23.2 | NA | 1.04 | 59.4 | 3.58 | 35.98 | 0 | 0 | hump-shaped |
| Affinity | Light | *T. minimum* | GAM | m. decrease | NA | NA | 3.18 | 29.64 | 0.54 | 66.64 | 0 | 0 | m. decrease |
| Affinity | Nitrogen | *Coelastrum sp.* | GAM | hump-shaped | 23.5 | NA | 0 | 94 | 5.84 | 0.16 | 0 | 0 | hump-shaped |
| Affinity | Nitrogen | *E. elegans* | GAM | hump-shaped | 24.7 | NA | 0 | 84.8 | 13.3 | 1.9 | 0 | 0 | hump-shaped |
| Affinity | Nitrogen | *F. crotonensis* | LM | l. increase | NA | 6.00E-04 | 0 | 0 | 0 | 0 | 16.7 | 83.3 | l. increase |
| Affinity | Nitrogen | *F. saprophila* | LM | l. decrease | NA | -0.002 | 0 | 0 | 0 | 0 | 71.8 | 28.2 | l. decrease |
| Affinity | Nitrogen | *L. subsalsa* | GAM | hump-shaped | 24.1 | NA | 0 | 100 | 0 | 0 | 0 | 0 | hump-shaped |

| | | | | | | | | | | | | | |
|---|---|---|---|---|---|---|---|---|---|---|---|---|---|
| Affinity | Nitrogen | *M. aeruginosa st. 1* | LM | l. decrease | NA | -0.0128 | 0 | 0 | 0 | 0 | 100 | 0 | l. decrease |
| Affinity | Nitrogen | *M. aeruginosa st. 2* | LM | l. decrease | NA | -0.0012 | 0 | 0 | 0 | 0 | 99.98 | 0.02 | l. decrease |
| Affinity | Nitrogen | *M. gracile* | GAM | U-shaped | 24.5 | NA | 45.84 | 2.24 | 14.02 | 37.9 | 0 | 0 | U-shaped |
| Affinity | Nitrogen | *N. palea (C)* | GAM | U-shaped | 18.9 | NA | 44.82 | 0 | 55.1 | 0.08 | 0 | 0 | m. increase |
| Affinity | Nitrogen | *N. palea (G)* | GAM | hump-shaped | 24.4 | NA | 0 | 100 | 0 | 0 | 0 | 0 | hump-shaped |
| Affinity | Nitrogen | *Oocystis sp.* | LM | l. decrease | NA | -0.0836 | 0 | 0 | 0 | 0 | 100 | 0 | l. decrease |
| Affinity | Nitrogen | *P. boryanum* | GAM | m. decrease | NA | NA | 0.02 | 33.16 | 0.56 | 66.26 | 0 | 0 | m. decrease |
| Affinity | Nitrogen | *S. arcuatus* | LM | l. increase | NA | 0.1155 | 0 | 0 | 0 | 0 | 0.94 | 99.06 | l. increase |
| Affinity | Nitrogen | *S. armatus* | GAM | hump-shaped | 22 | NA | 0 | 71.36 | 1.92 | 26.72 | 0 | 0 | hump-shaped |
| Affinity | Nitrogen | *T. minimum* | GAM | m. increase | NA | NA | 0.26 | 3.98 | 95.76 | 0 | 0 | 0 | m. increase |
| Affinity | Phosphorus | *A. formosa* | LM | l. decrease | NA | -1.00E-04 | 0 | 0 | 0 | 0 | 60.28 | 39.72 | l. decrease |
| Affinity | Phosphorus | *Chlorella sp.* | LM | l. increase | NA | 0.1039 | 0 | 0 | 0 | 0 | 9.84 | 90.16 | l. increase |
| Affinity | Phosphorus | *Coelastrum sp.* | LM | l. decrease | NA | -0.0099 | 0 | 0 | 0 | 0 | 48.06 | 51.94 | l. increase |
| Affinity | Phosphorus | *E. elegans* | GAM | hump-shaped | 25.6 | NA | 0.02 | 72.34 | 26 | 1.64 | 0 | 0 | hump-shaped |
| Affinity | Phosphorus | *F. crotonensis* | LM | l. increase | NA | 0.0022 | 0 | 0 | 0 | 0 | 0.4 | 99.6 | l. increase |
| Affinity | Phosphorus | *F. saprophila* | LM | l. decrease | NA | -0.0031 | 0 | 0 | 0 | 0 | 94.52 | 5.48 | l. decrease |
| Affinity | Phosphorus | *L. subsalsa* | LM | l. increase | NA | 0.0323 | 0 | 0 | 0 | 0 | 3.92 | 96.08 | l. increase |
| Affinity | Phosphorus | *M. aeruginosa st. 1* | GAM | U-shaped | 22.2 | NA | 25.4 | 9.72 | 47.78 | 17.1 | 0 | 0 | m. increase |
| Affinity | Phosphorus | *M. aeruginosa st. 2* | LM | l. increase | NA | 0.005 | 0 | 0 | 0 | 0 | 30.9 | 69.1 | l. increase |
| Affinity | Phosphorus | *M. gracile* | GAM | hump-shaped | 25.1 | NA | 0.02 | 63.16 | 25.14 | 11.68 | 0 | 0 | hump-shaped |
| Affinity | Phosphorus | *N. palea (C)* | GAM | m. increase | NA | NA | 21 | 0.32 | 78.26 | 0.42 | 0 | 0 | m. increase |
| Affinity | Phosphorus | *N. palea (G)* | GAM | m. increase | NA | NA | 0.26 | 43.42 | 53.1 | 3.22 | 0 | 0 | m. increase |
| Affinity | Phosphorus | *Oocystis sp.* | GAM | U-shaped | 15.4 | NA | 0 | 69.32 | 0.08 | 30.6 | 0 | 0 | hump-shaped |
| Affinity | Phosphorus | *P. boryanum* | GAM | U-shaped | 25.8 | NA | 29.7 | 15.3 | 29.02 | 25.98 | 0 | 0 | U-shaped |
| Affinity | Phosphorus | *S. arcuatus* | GAM | m. increase | NA | NA | 10.48 | 25.74 | 50.36 | 13.42 | 0 | 0 | m. increase |
| Affinity | Phosphorus | *S. armatus* | GAM | hump-shaped | 25.3 | NA | 1 | 55 | 29.7 | 14.3 | 0 | 0 | hump-shaped |
| Affinity | Phosphorus | *Synechococcus sp.* | LM | l. decrease | NA | -4.00E-04 | 0 | 0 | 0 | 0 | 58.04 | 41.96 | l. decrease |
| Affinity | Phosphorus | *T. minimum* | GAM | U-shaped | 18.4 | NA | 22.64 | 3.98 | 61.94 | 11.44 | 0 | 0 | m. increase |

**Table S8.** Summary table of the EDF categorizations for the mean value across populations ("Mean spp"– see method) resource-dependent growth parameters, for the three resources. In the "Data" column we have the parameters; in the "Experiment" column we have the resource type; in "Species" we have "mean_spp"; in "Model" we have whether a linear model (LM) or a generalized additive model (GAM) fitted the data best; in "Category" this correspond to the shape attributed to the curves fitted to the mean values across populations, we can find the following option: U-shape, hump-shape, monotonic or linear increase or decrease; the "Critical point" is the inflexion temperature, which is for a U-shape or hump-shape at which temperature the minimum or maximum was respectively reached; "Slope" if the mean fit curve was determined to be best fitted with a LM; "p-value" is the value from the fitted curve to assess if it is significant; finally "Sig. p-value" is the significance of the fitted curve in "yes"/"no" statement regardless of the value.

| **Data** | **Experiment** | **Species** | **Model** | **Category** | **Critical point** | **Slope** | **p-value** | **Sig p-value** |
|---|---|---|---|---|---|---|---|---|
| Rstar | Light | mean_spp | GAM | U-shaped | 24.1 | NA | 0.05210032 | no |
| Rstar | Nitrogen | mean_spp | LM | l. increase | NA | 0.577 | 0.03407732 | yes |
| Rstar | Phosphorus | mean_spp | GAM | U-shaped | 23.7 | NA | 0.17017633 | no |
| Net mumax | Light | mean_spp | GAM | hump-shaped | 27.6 | NA | 0.0252319 | yes |
| Net mumax | Nitrogen | mean_spp | GAM | hump-shaped | 27.5 | NA | 9.07E-04 | yes |
| Net mumax | Phosphorus | mean_spp | GAM | hump-shaped | 25.8 | NA | 0.16167964 | no |
| Affinity | Light | mean_spp | GAM | hump-shaped | 22.4 | NA | 0.06275534 | no |
| Affinity | Nitrogen | mean_spp | GAM | hump-shaped | 23.4 | NA | 0.13256952 | no |
| Affinity | Phosphorus | mean_spp | LM | l. decrease | NA | -0.01 | 0.69732458 | no |

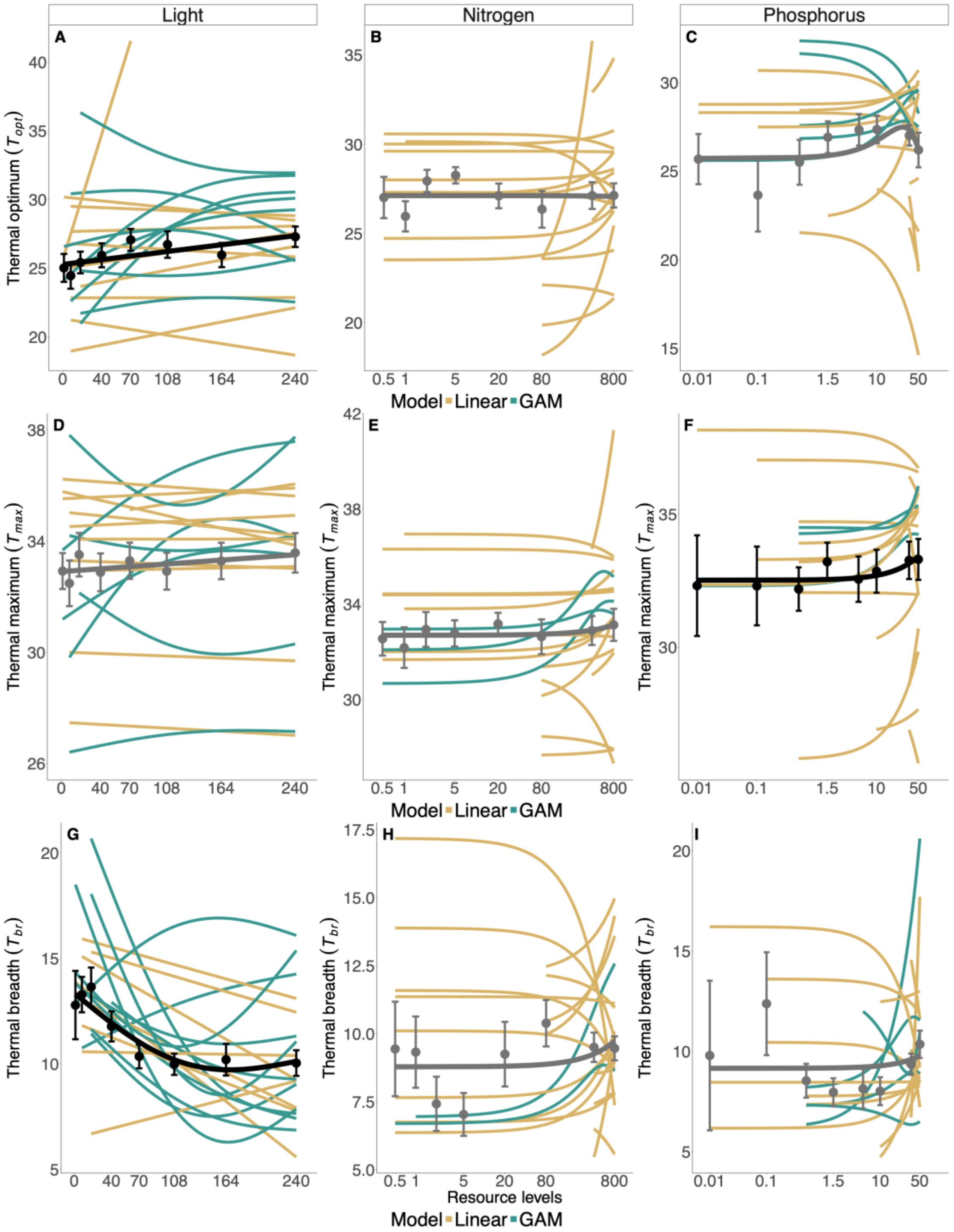

**Figure S15**. EDF categorizations for GAMs from Fig. 3 (main text). Thermal performance curve parameters plotted as a function of resource availability for each experiment (light, nitrogen and phosphorus from left to right). Lines represent model fits for each species and significant models are accompanied by shaded standard error ribbons. Gold and green colors represent the type of model that best fits the data, either a linear model or a generalized additive model (see methods). Points represent the mean parameter across all populations ("Mean spp"), and error bars represent the standard error of the mean. Black and grey lines represent the model fits of the mean parameter, black lines represent significant models ($p < 0.05$) and grey lines represent non-significant models. A-C are the thermal optimum (*Topt*). D-F are the thermal maximum (*Tmax*). G-I are the thermal breadths (*Tbr*). All the temperatures are reported in °C. For easier visual representation, the resource axes were logged for the nitrogen and phosphorus panels (B&C, E&F, and H&I), and models were fit to log-transformed values.

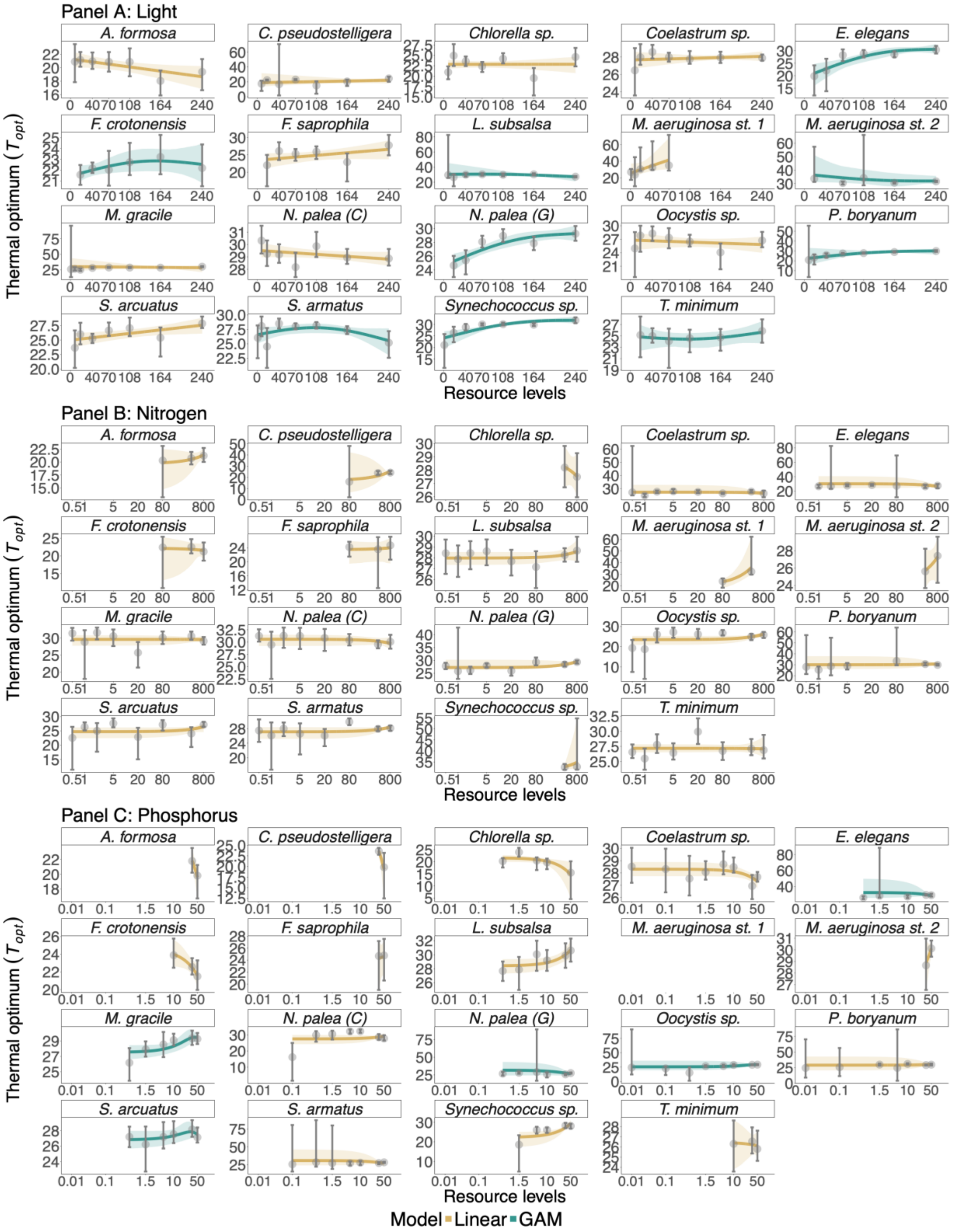

**<u>Figure S16</u>**. EDF categorizations for optimal temperature for growth rate (*Topt*). We have one panel per resource and each panel is faceted per population. Gold and green colors represent the type of model that best fits the data, either a linear model or a generalized additive model (GAM- see methods). Points and errors bars represent the medians value and their 95% credible interval. The curves are the mean fits from the posteriors bootstrapping procedure and the shaded ribbon their 95% credible interval (see method).

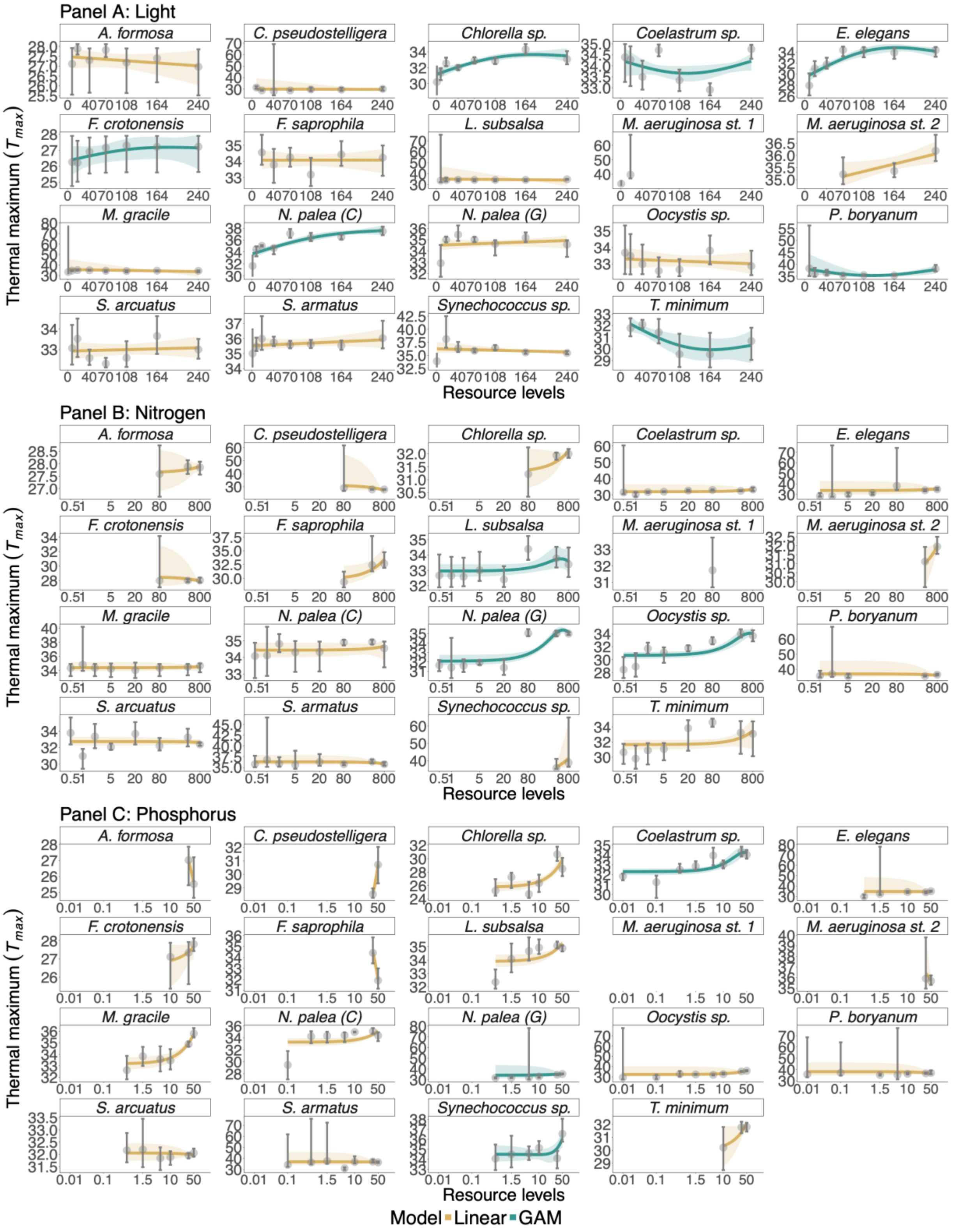

**Figure S17**. EDF categorizations for thermal maximum (*Tmax*). We have one panel per resource and each panel is faceted per population. Gold and green colors represent the type of model that best fits the data, either a linear model or a generalized additive model (GAM- see methods). Points and errors bars represent the medians value and their 95% credible interval. The curves are the mean fits from the posteriors bootstrapping procedure and the shaded ribbon their 95% credible interval (see method).

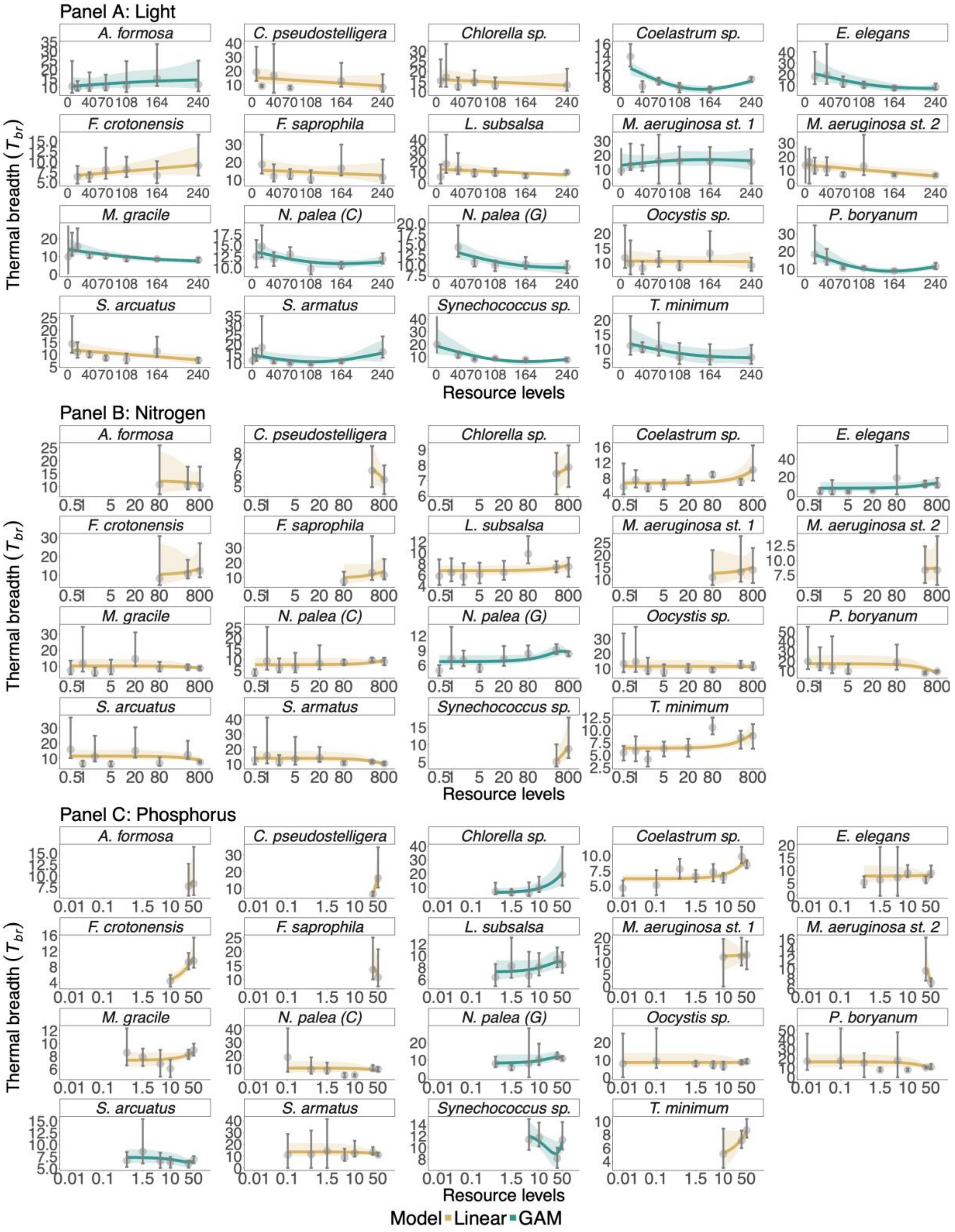

**<u>Figure S18</u>**. EDF categorizations for thermal breadth for growth rate (*Tbr*). We have one panel per resource and each panel is faceted per population. Gold and green colors represent the type of model that best fits the data, either a linear model or a generalized additive model (GAM- see methods). Points and errors bars represent the medians value and their 95% credible interval. The curves are the mean fits from the posteriors bootstrapping procedure and the shaded ribbon their 95% credible interval (see method).

**Table S9**. Summary table of the EDF categorizations for the thermal performance curve parameters, for the three resources. In the “Data” column we have the parameters; in the “Experiment” column we have the resource type; in “Species” we find the short species name, see full name in the first column of Table S3; in “Model” we have whether a linear model (LM) or a generalized additive model (GAM) fitted the data best; in “M. fit Category” this correspond to the shape attributed to the curves from the posteriors bootstrapping procedure, we can find the following option: U-shape, hump-shape, monotonic or linear increase or decrease; the “Critical point” is the inflexion resource level, which is for a U-shape or hump-shape at which resource value the minimum or maximum was respectively reached; “Slope” if the mean fit curve was determined to be best fitted with a LM, otherwise contains NA; “U_sh” is U-shaped, “H_sh” is hump-shaped, “M_inc” and “M_dec” are respectively monotonic increase and decrease and “LM_dec” and “LM_inc” are linear decrease and increase, these categories show the percentage of the 5,000 posteriors that were sorted in each; and finally the “T. Category” which stands for “True Category”, contains the identity of the shape for the parameter having more than 51% of the fits (see methods) in the absent of a majority; the “T. Category” says uncertain.

| Data | Experiment | Species | Model | M. fit Category | Critical point | Slope | U_sh | H_sh | M_inc | M_dec | LM_dec | LM_inc | T. Category |
|---|---|---|---|---|---|---|---|---|---|---|---|---|---|
| Topt | Light | *A. formosa* | LM | l. decrease | NA | -0.0093 | 0 | 0 | 0 | 0 | 98.68 | 1.32 | l. decrease |
| Topt | Light | *C. pseudostelligera* | LM | l. increase | NA | 0.0044 | 0 | 0 | 0 | 0 | 14.1 | 85.9 | l. increase |
| Topt | Light | *Chlorella sp.* | LM | l. decrease | NA | -0.0022 | 0 | 0 | 0 | 0 | 47.6 | 52.4 | l. increase |
| Topt | Light | *Coelastrum sp.* | LM | l. increase | NA | 6.00E-04 | 0 | 0 | 0 | 0 | 29.04 | 70.96 | l. increase |
| Topt | Light | *E. elegans* | GAM | m. increase | NA | NA | 0 | 49.2 | 50.84 | 0 | 0 | 0 | m. increase |
| Topt | Light | *F. crotonensis* | GAM | hump-shaped | 156.5 | NA | 1.1 | 57.5 | 38.02 | 3.38 | 0 | 0 | hump-shaped |
| Topt | Light | *F. saprophila* | LM | l. increase | NA | 0.0153 | 0 | 0 | 0 | 0 | 6.44 | 93.56 | l. increase |
| Topt | Light | *L. subsalsa* | GAM | hump-shaped | 67.2 | NA | 0 | 65.9 | 7.46 | 26.7 | 0 | 0 | hump-shaped |
| Topt | Light | *M. aeruginosa st. 1* | LM | l. increase | NA | 0.1106 | 0 | 0 | 0 | 0 | 3.16 | 96.84 | l. increase |
| Topt | Light | *M. aeruginosa st. 2* | GAM | U-shaped | 224.3 | NA | 37.3 | 0.02 | 3.44 | 59.2 | 0 | 0 | m. decrease |
| Topt | Light | *M. gracile* | LM | l. increase | NA | 0.0287 | 0 | 0 | 0 | 0 | 33.08 | 66.92 | l. increase |

| | | | | | | | | | | | | | |
|---|---|---|---|---|---|---|---|---|---|---|---|---|---|
| Topt | Light | *N. palea (C)* | LM | l. decrease | NA | -0.0043 | 0 | 0 | 0 | 0 | 93.42 | 6.58 | l. decrease |
| Topt | Light | *N. palea (G)* | GAM | m. increase | NA | NA | 0 | 38.8 | 61.2 | 0 | 0 | 0 | m. increase |
| Topt | Light | *Oocystis sp.* | LM | l. increase | NA | 0.0019 | 0 | 0 | 0 | 0 | 76.58 | 23.42 | l. decrease |
| Topt | Light | *P. boryanum* | GAM | m. increase | NA | NA | 0.06 | 32.4 | 63.76 | 3.8 | 0 | 0 | m. increase |
| Topt | Light | *S. arcuatus* | LM | l. increase | NA | 0.0111 | 0 | 0 | 0 | 0 | 1.18 | 98.82 | l. increase |
| Topt | Light | *S. armatus* | GAM | hump-shaped | 104.2 | NA | 0 | 87.1 | 4.88 | 8 | 0 | 0 | hump-shaped |
| Topt | Light | *Synechococcus sp.* | GAM | m. increase | NA | NA | 0 | 47.3 | 52.7 | 0 | 0 | 0 | m. increase |
| Topt | Light | *T. minimum* | GAM | U-shaped | 102.8 | NA | 47.7 | 2.14 | 40.14 | 10 | 0 | 0 | U-shaped |
| Topt | Nitrogen | *A. formosa* | LM | l. increase | NA | 0.0024 | 0 | 0 | 0 | 0 | 26.72 | 73.28 | l. increase |
| Topt | Nitrogen | *C. pseudostelligera* | LM | l. increase | NA | 0.015 | 0 | 0 | 0 | 0 | 7.8 | 92.2 | l. increase |
| Topt | Nitrogen | *Chlorella sp.* | LM | l. increase | NA | 0.0023 | 0 | 0 | 0 | 0 | 73.4 | 26.6 | l. decrease |
| Topt | Nitrogen | *Coelastrum sp.* | LM | l. increase | NA | 0.001 | 0 | 0 | 0 | 0 | 60.68 | 39.32 | l. decrease |
| Topt | Nitrogen | *E. elegans* | LM | l. decrease | NA | -0.0022 | 0 | 0 | 0 | 0 | 66.74 | 33.26 | l. decrease |
| Topt | Nitrogen | *F. crotonensis* | LM | l. increase | NA | 0.0084 | 0 | 0 | 0 | 0 | 66.06 | 33.94 | l. decrease |
| Topt | Nitrogen | *F. saprophila* | LM | l. decrease | NA | -0.0011 | 0 | 0 | 0 | 0 | 35.46 | 64.54 | l. increase |
| Topt | Nitrogen | *L. subsalsa* | LM | l. increase | NA | 3.00E-04 | 0 | 0 | 0 | 0 | 16.82 | 83.18 | l. increase |
| Topt | Nitrogen | *M. aeruginosa st. 1* | LM | l. increase | NA | 0.0204 | 0 | 0 | 0 | 0 | 0.04 | 99.96 | l. increase |
| Topt | Nitrogen | *M. aeruginosa st. 2* | LM | l. increase | NA | 0.0077 | 0 | 0 | 0 | 0 | 19.16 | 80.84 | l. increase |
| Topt | Nitrogen | *M. gracile* | LM | l. decrease | NA | -0.0016 | 0 | 0 | 0 | 0 | 56.26 | 43.74 | l. decrease |
| Topt | Nitrogen | *N. palea (C)* | LM | l. decrease | NA | -8.00E-04 | 0 | 0 | 0 | 0 | 84.24 | 15.76 | l. decrease |
| Topt | Nitrogen | *N. palea (G)* | LM | l. increase | NA | 0.0031 | 0 | 0 | 0 | 0 | 2.64 | 97.36 | l. increase |
| Topt | Nitrogen | *Oocystis sp.* | LM | l. increase | NA | 0.0042 | 0 | 0 | 0 | 0 | 2.34 | 97.66 | l. increase |
| Topt | Nitrogen | *P. boryanum* | LM | l. decrease | NA | -0.006 | 0 | 0 | 0 | 0 | 27.66 | 72.34 | l. increase |
| Topt | Nitrogen | *S. arcuatus* | LM | l. increase | NA | 0.0024 | 0 | 0 | 0 | 0 | 8 | 92 | l. increase |
| Topt | Nitrogen | *S. armatus* | LM | l. increase | NA | 0.0011 | 0 | 0 | 0 | 0 | 6.8 | 93.2 | l. increase |
| Topt | Nitrogen | *Synechococcus sp.* | LM | l. decrease | NA | -8.00E-04 | 0 | 0 | 0 | 0 | 37.98 | 62.02 | l. increase |
| Topt | Nitrogen | *T. minimum* | LM | l. decrease | NA | -0.0011 | 0 | 0 | 0 | 0 | 58.58 | 41.42 | l. decrease |
| Topt | Phosphorus | *A. formosa* | LM | l. decrease | NA | -0.1453 | 0 | 0 | 0 | 0 | 97.08 | 2.92 | l. decrease |

| | | | | | | | | | | | | | |
|---|---|---|---|---|---|---|---|---|---|---|---|---|---|
| Topt | Phosphorus | *C. pseudostelligera* | LM | l. decrease | NA | -0.2509 | 0 | 0 | 0 | 0 | 98.4 | 1.6 | l. decrease |
| Topt | Phosphorus | *Chlorella sp.* | LM | l. decrease | NA | -0.0488 | 0 | 0 | 0 | 0 | 99.32 | 0.68 | l. decrease |
| Topt | Phosphorus | *Coelastrum sp.* | LM | l. decrease | NA | -0.0341 | 0 | 0 | 0 | 0 | 98.78 | 1.22 | l. decrease |
| Topt | Phosphorus | *E. elegans* | GAM | m. decrease | NA | NA | 0.06 | 14.3 | 47.42 | 38.2 | 0 | 0 | m. increase |
| Topt | Phosphorus | *F. crotonensis* | LM | l. decrease | NA | -0.0868 | 0 | 0 | 0 | 0 | 98.38 | 1.62 | l. decrease |
| Topt | Phosphorus | *F. saprophila* | LM | l. decrease | NA | -0.2106 | 0 | 0 | 0 | 0 | 46.58 | 53.42 | l. increase |
| Topt | Phosphorus | *L. subsalsa* | LM | l. increase | NA | 0.0522 | 0 | 0 | 0 | 0 | 0.88 | 99.12 | l. increase |
| Topt | Phosphorus | *M. aeruginosa st. 2* | LM | l. decrease | NA | -1.0544 | 0 | 0 | 0 | 0 | 6.98 | 93.02 | l. increase |
| Topt | Phosphorus | *M. gracile* | GAM | hump-shaped | 41.2 | NA | 0.04 | 47.8 | 52.12 | 0.04 | 0 | 0 | m. increase |
| Topt | Phosphorus | *N. palea (C)* | LM | l. increase | NA | 0.0544 | 0 | 0 | 0 | 0 | 24.22 | 75.78 | l. increase |
| Topt | Phosphorus | *N. palea (G)* | GAM | m. decrease | NA | NA | 31 | 0 | 15.06 | 53.9 | 0 | 0 | m. decrease |
| Topt | Phosphorus | *Oocystis sp.* | GAM | m. increase | NA | NA | 0 | 22.8 | 50.3 | 26.9 | 0 | 0 | m. increase |
| Topt | Phosphorus | *P. boryanum* | LM | l. decrease | NA | -0.2155 | 0 | 0 | 0 | 0 | 35.04 | 64.96 | l. increase |
| Topt | Phosphorus | *S. arcuatus* | GAM | hump-shaped | 31.3 | NA | 0.32 | 52.2 | 38.58 | 8.86 | 0 | 0 | hump-shaped |
| Topt | Phosphorus | *S. armatus* | LM | l. increase | NA | 0.0059 | 0 | 0 | 0 | 0 | 46.02 | 53.98 | l. increase |
| Topt | Phosphorus | *Synechococcus sp.* | LM | l. increase | NA | 0.0974 | 0 | 0 | 0 | 0 | 0 | 100 | l. increase |
| Topt | Phosphorus | *T. minimum* | LM | l. increase | NA | 0.0329 | 0 | 0 | 0 | 0 | 61.56 | 38.44 | l. decrease |
| Tmax | Light | *A. formosa* | LM | l. decrease | NA | -3.00E-04 | 0 | 0 | 0 | 0 | 76.78 | 23.22 | l. decrease |
| Tmax | Light | *C. pseudostelligera* | LM | l. decrease | NA | -3.00E-04 | 0 | 0 | 0 | 0 | 36.4 | 63.6 | l. increase |
| Tmax | Light | *Chlorella sp.* | GAM | hump-shaped | 177.8 | NA | 0 | 76 | 23.94 | 0.04 | 0 | 0 | hump-shaped |
| Tmax | Light | *Coelastrum sp.* | GAM | U-shaped | 122.1 | NA | 80.1 | 0 | 16.4 | 3.5 | 0 | 0 | U-shaped |
| Tmax | Light | *E. elegans* | GAM | hump-shaped | 167.6 | NA | 0 | 92.9 | 7.06 | 0 | 0 | 0 | hump-shaped |
| Tmax | Light | *F. crotonensis* | GAM | hump-shaped | 185.8 | NA | 2.76 | 34.6 | 55.42 | 7.2 | 0 | 0 | m. increase |
| Tmax | Light | *F. saprophila* | LM | l. increase | NA | 0.0026 | 0 | 0 | 0 | 0 | 48.06 | 51.94 | l. increase |
| Tmax | Light | *L. subsalsa* | LM | l. increase | NA | 0.0013 | 0 | 0 | 0 | 0 | 16.8 | 83.2 | l. increase |
| Tmax | Light | *M. aeruginosa st. 2* | LM | l. increase | NA | 0.0106 | 0 | 0 | 0 | 0 | 1.52 | 98.48 | l. increase |
| Tmax | Light | *M. gracile* | LM | l. decrease | NA | -0.001 | 0 | 0 | 0 | 0 | 63.66 | 36.34 | l. decrease |

| | | | | | | | | | | | | | |
|---|---|---|---|---|---|---|---|---|---|---|---|---|---|
| Tmax | Light | *N. palea (C)* | GAM | m. increase | NA | NA | 0 | 27.7 | 72.3 | 0 | 0 | 0 | m. increase |
| Tmax | Light | *N. palea (G)* | LM | l. decrease | NA | -0.0011 | 0 | 0 | 0 | 0 | 17.88 | 82.12 | l. increase |
| Tmax | Light | *Oocystis sp.* | LM | l. increase | NA | 0.0028 | 0 | 0 | 0 | 0 | 69.04 | 30.96 | l. decrease |
| Tmax | Light | *P. boryanum* | GAM | U-shaped | 123.3 | NA | 98.4 | 0 | 1.12 | 0.5 | 0 | 0 | U-shaped |
| Tmax | Light | *S. arcuatus* | LM | l. decrease | NA | -0.003 | 0 | 0 | 0 | 0 | 34.52 | 65.48 | l. increase |
| Tmax | Light | *S. armatus* | LM | l. increase | NA | 0.0015 | 0 | 0 | 0 | 0 | 19.62 | 80.38 | l. increase |
| Tmax | Light | *Synechococcus sp.* | LM | l. decrease | NA | -0.0095 | 0 | 0 | 0 | 0 | 80.28 | 19.72 | l. decrease |
| Tmax | Light | *T. minimum* | GAM | U-shaped | 166.5 | NA | 69.4 | 0 | 0.14 | 30.5 | 0 | 0 | U-shaped |
| Tmax | Nitrogen | *A. formosa* | LM | l. decrease | NA | -7.00E-04 | 0 | 0 | 0 | 0 | 27.92 | 72.08 | l. increase |
| Tmax | Nitrogen | *C. pseudostelligera* | LM | l. decrease | NA | -1.00E-04 | 0 | 0 | 0 | 0 | 80.56 | 19.44 | l. decrease |
| Tmax | Nitrogen | *Chlorella sp.* | LM | l. increase | NA | 0.0017 | 0 | 0 | 0 | 0 | 6.36 | 93.64 | l. increase |
| Tmax | Nitrogen | *Coelastrum sp.* | LM | l. increase | NA | 0.0025 | 0 | 0 | 0 | 0 | 3.34 | 96.66 | l. increase |
| Tmax | Nitrogen | *E. elegans* | LM | l. decrease | NA | -0.0016 | 0 | 0 | 0 | 0 | 28.9 | 71.1 | l. increase |
| Tmax | Nitrogen | *F. crotonensis* | LM | l. decrease | NA | -0.0016 | 0 | 0 | 0 | 0 | 47.46 | 52.54 | l. increase |
| Tmax | Nitrogen | *F. saprophila* | LM | l. increase | NA | 0.0053 | 0 | 0 | 0 | 0 | 0.1 | 99.9 | l. increase |
| Tmax | Nitrogen | *L. subsalsa* | GAM | hump-shaped | 510 | NA | 0 | 51.6 | 45.84 | 2.52 | 0 | 0 | hump-shaped |
| Tmax | Nitrogen | *M. aeruginosa st. 2* | LM | l. increase | NA | 0.0042 | 0 | 0 | 0 | 0 | 3.78 | 96.22 | l. increase |
| Tmax | Nitrogen | *M. gracile* | LM | l. increase | NA | 0.001 | 0 | 0 | 0 | 0 | 28.76 | 71.24 | l. increase |
| Tmax | Nitrogen | *N. palea (C)* | LM | l. increase | NA | 9.00E-04 | 0 | 0 | 0 | 0 | 25.06 | 74.94 | l. increase |
| Tmax | Nitrogen | *N. palea (G)* | GAM | hump-shaped | 583.1 | NA | 0 | 92.1 | 6 | 1.94 | 0 | 0 | hump-shaped |
| Tmax | Nitrogen | *Oocystis sp.* | GAM | hump-shaped | 718.1 | NA | 0 | 55.2 | 44.76 | 0.04 | 0 | 0 | hump-shaped |
| Tmax | Nitrogen | *P. boryanum* | LM | l. increase | NA | 6.00E-04 | 0 | 0 | 0 | 0 | 46.48 | 53.52 | l. increase |
| Tmax | Nitrogen | *S. arcuatus* | LM | l. decrease | NA | -1.00E-04 | 0 | 0 | 0 | 0 | 60.78 | 39.22 | l. decrease |
| Tmax | Nitrogen | *S. armatus* | LM | l. increase | NA | 2.00E-04 | 0 | 0 | 0 | 0 | 72.08 | 27.92 | l. decrease |
| Tmax | Nitrogen | *Synechococcus sp.* | LM | l. increase | NA | 0.0098 | 0 | 0 | 0 | 0 | 3.95 | 96.05 | l. increase |
| Tmax | Nitrogen | *T. minimum* | LM | l. increase | NA | 6.00E-04 | 0 | 0 | 0 | 0 | 9.18 | 90.82 | l. increase |
| Tmax | Phosphorus | *A. formosa* | LM | l. increase | NA | 0.0286 | 0 | 0 | 0 | 0 | 90.26 | 9.74 | l. decrease |
| Tmax | Phosphorus | *C. pseudostelligera* | LM | l. increase | NA | 0.1595 | 0 | 0 | 0 | 0 | 0.18 | 99.82 | l. increase |

| | | | | | | | | | | | | | |
|---|---|---|---|---|---|---|---|---|---|---|---|---|---|
| Tmax | Phosphorus | *Chlorella sp.* | LM | l. increase | NA | 0.0744 | 0 | 0 | 0 | 0 | 0 | 100 | l. increase |
| Tmax | Phosphorus | *Coelastrum sp.* | GAM | hump-shaped | 43 | NA | 0 | 56.7 | 43.34 | 0 | 0 | 0 | hump-shaped |
| Tmax | Phosphorus | *E. elegans* | LM | l. decrease | NA | -0.2084 | 0 | 0 | 0 | 0 | 22.86 | 77.14 | l. increase |
| Tmax | Phosphorus | *F. crotonensis* | LM | l. increase | NA | 0.0077 | 0 | 0 | 0 | 0 | 12.62 | 87.38 | l. increase |
| Tmax | Phosphorus | *F. saprophila* | LM | l. decrease | NA | -0.18 | 0 | 0 | 0 | 0 | 99.84 | 0.16 | l. decrease |
| Tmax | Phosphorus | *L. subsalsa* | LM | l. increase | NA | 0.0278 | 0 | 0 | 0 | 0 | 0.36 | 99.64 | l. increase |
| Tmax | Phosphorus | *M. aeruginosa st. 2* | LM | l. decrease | NA | -1.7333 | 0 | 0 | 0 | 0 | 63.73 | 36.27 | l. decrease |
| Tmax | Phosphorus | *M. gracile* | LM | l. increase | NA | 0.0564 | 0 | 0 | 0 | 0 | 0 | 100 | l. increase |
| Tmax | Phosphorus | *N. palea (C)* | LM | l. increase | NA | 0.0446 | 0 | 0 | 0 | 0 | 0.12 | 99.88 | l. increase |
| Tmax | Phosphorus | *N. palea (G)* | GAM | m. increase | NA | NA | 0 | 1.84 | 71.9 | 26.3 | 0 | 0 | m. increase |
| Tmax | Phosphorus | *Oocystis sp.* | LM | l. increase | NA | 0.1051 | 0 | 0 | 0 | 0 | 15.36 | 84.64 | l. increase |
| Tmax | Phosphorus | *P. boryanum* | LM | l. decrease | NA | -0.117 | 0 | 0 | 0 | 0 | 47 | 53 | l. increase |
| Tmax | Phosphorus | *S. arcuatus* | LM | l. decrease | NA | -0.0024 | 0 | 0 | 0 | 0 | 60.12 | 39.88 | l. decrease |
| Tmax | Phosphorus | *S. armatus* | LM | l. increase | NA | 0.0431 | 0 | 0 | 0 | 0 | 29.8 | 70.2 | l. increase |
| Tmax | Phosphorus | *Synechococcus sp.* | GAM | U-shaped | 8.9 | NA | 43.9 | 0 | 55.92 | 0.14 | 0 | 0 | m. increase |
| Tmax | Phosphorus | *T. minimum* | LM | l. increase | NA | 0.036 | 0 | 0 | 0 | 0 | 3.22 | 96.78 | l. increase |
| Tbr | Light | *A. formosa* | GAM | m. increase | NA | NA | 7.9 | 24 | 54.2 | 13.9 | 0 | 0 | m. increase |
| Tbr | Light | *C. pseudostelligera* | LM | l. decrease | NA | -0.0047 | 0 | 0 | 0 | 0 | 90.98 | 9.02 | l. decrease |
| Tbr | Light | *Chlorella sp.* | LM | l. decrease | NA | -0.0112 | 0 | 0 | 0 | 0 | 82.02 | 17.98 | l. decrease |
| Tbr | Light | *Coelastrum sp.* | GAM | U-shaped | 149.5 | NA | 99.6 | 0 | 0 | 0.38 | 0 | 0 | U-shaped |
| Tbr | Light | *E. elegans* | GAM | m. decrease | NA | NA | 47.3 | 0 | 0.02 | 52.7 | 0 | 0 | m. decrease |
| Tbr | Light | *F. crotonensis* | LM | l. increase | NA | 0.0366 | 0 | 0 | 0 | 0 | 8.8 | 91.2 | l. increase |
| Tbr | Light | *F. saprophila* | LM | l. decrease | NA | -0.0242 | 0 | 0 | 0 | 0 | 77.44 | 22.56 | l. decrease |
| Tbr | Light | *L. subsalsa* | LM | l. decrease | NA | -0.0172 | 0 | 0 | 0 | 0 | 94.04 | 5.96 | l. decrease |
| Tbr | Light | *M. aeruginosa st. 1* | GAM | hump-shaped | 155.8 | NA | 2.44 | 39.7 | 42.94 | 15 | 0 | 0 | m. increase |
| Tbr | Light | *M. aeruginosa st. 2* | LM | l. decrease | NA | -0.0238 | 0 | 0 | 0 | 0 | 99.92 | 0.08 | l. decrease |
| Tbr | Light | *M. gracile* | GAM | m. decrease | NA | NA | 24.7 | 0 | 0.02 | 75.3 | 0 | 0 | m. decrease |
| Tbr | Light | *N. palea (C)* | GAM | U-shaped | 165.7 | NA | 63.3 | 0 | 0.56 | 36.1 | 0 | 0 | U-shaped |

| | | | | | | | | | | | | | |
|---|---|---|---|---|---|---|---|---|---|---|---|---|---|
| Tbr | Light | *N. palea (G)* | GAM | m. decrease | NA | NA | 38.4 | 0.08 | 0.04 | 61.5 | 0 | 0 | m. decrease |
| Tbr | Light | *Oocystis sp.* | LM | l. decrease | NA | -0.0014 | 0 | 0 | 0 | 0 | 49.56 | 50.44 | l. increase |
| Tbr | Light | *P. boryanum* | GAM | U-shaped | 157.6 | NA | 97.2 | 0 | 0 | 2.78 | 0 | 0 | U-shaped |
| Tbr | Light | *S. arcuatus* | LM | l. decrease | NA | -0.0226 | 0 | 0 | 0 | 0 | 98.56 | 1.44 | l. decrease |
| Tbr | Light | *S. armatus* | GAM | U-shaped | 112.7 | NA | 90.8 | 0 | 0.36 | 8.86 | 0 | 0 | U-shaped |
| Tbr | Light | *Synechococcus sp.* | GAM | U-shaped | 166.2 | NA | 95.1 | 0 | 0 | 4.88 | 0 | 0 | U-shaped |
| Tbr | Light | *T. minimum* | GAM | m. decrease | NA | NA | 36.1 | 1.22 | 1 | 61.6 | 0 | 0 | m. decrease |
| Tbr | Nitrogen | *A. formosa* | LM | l. decrease | NA | -0.0061 | 0 | 0 | 0 | 0 | 52.98 | 47.02 | l. decrease |
| Tbr | Nitrogen | *C. pseudostelligera* | LM | l. decrease | NA | -0.0047 | 0 | 0 | 0 | 0 | 81.66 | 18.34 | l. decrease |
| Tbr | Nitrogen | *Chlorella sp.* | LM | l. decrease | NA | -0.0015 | 0 | 0 | 0 | 0 | 31 | 69 | l. increase |
| Tbr | Nitrogen | *Coelastrum sp.* | LM | l. increase | NA | 0.0029 | 0 | 0 | 0 | 0 | 1.38 | 98.62 | l. increase |
| Tbr | Nitrogen | *E. elegans* | GAM | m. increase | NA | NA | 0.04 | 20.2 | 74.9 | 4.86 | 0 | 0 | m. increase |
| Tbr | Nitrogen | *F. crotonensis* | LM | l. decrease | NA | -0.0012 | 0 | 0 | 0 | 0 | 23.28 | 76.72 | l. increase |
| Tbr | Nitrogen | *F. saprophila* | LM | l. increase | NA | 0.0072 | 0 | 0 | 0 | 0 | 10.66 | 89.34 | l. increase |
| Tbr | Nitrogen | *L. subsalsa* | LM | l. increase | NA | 0.0022 | 0 | 0 | 0 | 0 | 13.14 | 86.86 | l. increase |
| Tbr | Nitrogen | *M. aeruginosa st. 1* | LM | l. increase | NA | 0.0136 | 0 | 0 | 0 | 0 | 27.36 | 72.64 | l. increase |
| Tbr | Nitrogen | *M. aeruginosa st. 2* | LM | l. decrease | NA | -8.00E-04 | 0 | 0 | 0 | 0 | 50.28 | 49.72 | l. decrease |
| Tbr | Nitrogen | *M. gracile* | LM | l. increase | NA | 0.0014 | 0 | 0 | 0 | 0 | 72.46 | 27.54 | l. decrease |
| Tbr | Nitrogen | *N. palea (C)* | LM | l. increase | NA | 0.0038 | 0 | 0 | 0 | 0 | 9.46 | 90.54 | l. increase |
| Tbr | Nitrogen | *N. palea (G)* | GAM | hump-shaped | 563.6 | NA | 0 | 57.9 | 40.72 | 1.38 | 0 | 0 | hump-shaped |
| Tbr | Nitrogen | *Oocystis sp.* | LM | l. increase | NA | 5.00E-04 | 0 | 0 | 0 | 0 | 48.24 | 51.76 | l. increase |
| Tbr | Nitrogen | *P. boryanum* | LM | l. decrease | NA | -0.0059 | 0 | 0 | 0 | 0 | 100 | 0 | l. decrease |
| Tbr | Nitrogen | *S. arcuatus* | LM | l. decrease | NA | -0.0026 | 0 | 0 | 0 | 0 | 92.3 | 7.7 | l. decrease |
| Tbr | Nitrogen | *S. armatus* | LM | l. decrease | NA | -0.0045 | 0 | 0 | 0 | 0 | 99.78 | 0.22 | l. decrease |
| Tbr | Nitrogen | *Synechococcus sp.* | LM | l. increase | NA | 0.0119 | 0 | 0 | 0 | 0 | 4.15 | 95.85 | l. increase |
| Tbr | Nitrogen | *T. minimum* | LM | l. increase | NA | 0.0021 | 0 | 0 | 0 | 0 | 0.86 | 99.14 | l. increase |
| Tbr | Phosphorus | *A. formosa* | LM | l. increase | NA | 0.1731 | 0 | 0 | 0 | 0 | 40.4 | 59.6 | l. increase |
| Tbr | Phosphorus | *C. pseudostelligera* | LM | l. increase | NA | 0.5269 | 0 | 0 | 0 | 0 | 0.02 | 99.98 | l. increase |

| | | | | | | | | | | | | | |
|---|---|---|---|---|---|---|---|---|---|---|---|---|---|
| Tbr | Phosphorus | *Chlorella sp.* | GAM | m. increase | NA | NA | 3.86 | 8.44 | 87.58 | 0.12 | 0 | 0 | m. increase |
| Tbr | Phosphorus | *Coelastrum sp.* | LM | l. increase | NA | 0.0857 | 0 | 0 | 0 | 0 | 0.02 | 99.98 | l. increase |
| Tbr | Phosphorus | *E. elegans* | LM | l. increase | NA | 0.0545 | 0 | 0 | 0 | 0 | 34.92 | 65.08 | l. increase |
| Tbr | Phosphorus | *F. crotonensis* | LM | l. increase | NA | 0.1033 | 0 | 0 | 0 | 0 | 0.02 | 99.98 | l. increase |
| Tbr | Phosphorus | *F. saprophila* | LM | l. increase | NA | 2.00E-04 | 0 | 0 | 0 | 0 | 77.94 | 22.06 | l. decrease |
| Tbr | Phosphorus | *L. subsalsa* | GAM | hump-shaped | 40.6 | NA | 0.28 | 33.6 | 60.54 | 5.58 | 0 | 0 | m. increase |
| Tbr | Phosphorus | *M. aeruginosa st. 1* | LM | l. decrease | NA | -0.0907 | 0 | 0 | 0 | 0 | 44.43 | 55.57 | l. increase |
| Tbr | Phosphorus | *M. aeruginosa st. 2* | LM | l. decrease | NA | -0.9299 | 0 | 0 | 0 | 0 | 98.52 | 1.48 | l. decrease |
| Tbr | Phosphorus | *M. gracile* | LM | l. increase | NA | 0.0232 | 0 | 0 | 0 | 0 | 5.82 | 94.18 | l. increase |
| Tbr | Phosphorus | *N. palea (C)* | LM | l. decrease | NA | -0.0514 | 0 | 0 | 0 | 0 | 67.24 | 32.76 | l. decrease |
| Tbr | Phosphorus | *N. palea (G)* | GAM | hump-shaped | 38.9 | NA | 0 | 42 | 52.3 | 5.7 | 0 | 0 | m. increase |
| Tbr | Phosphorus | *Oocystis sp.* | LM | l. increase | NA | 0.011 | 0 | 0 | 0 | 0 | 38.16 | 61.84 | l. increase |
| Tbr | Phosphorus | *P. boryanum* | LM | l. decrease | NA | -0.1054 | 0 | 0 | 0 | 0 | 98.14 | 1.86 | l. decrease |
| Tbr | Phosphorus | *S. arcuatus* | GAM | U-shaped | 35 | NA | 43 | 0.04 | 8.28 | 48.7 | 0 | 0 | m. decrease |
| Tbr | Phosphorus | *S. armatus* | LM | l. increase | NA | 0.0297 | 0 | 0 | 0 | 0 | 64.1 | 35.9 | l. decrease |
| Tbr | Phosphorus | *Synechococcus sp.* | GAM | U-shaped | 30.7 | NA | 78.8 | 0.1 | 2.08 | 19.1 | 0 | 0 | U-shaped |
| Tbr | Phosphorus | *T. minimum* | LM | l. increase | NA | 0.0634 | 0 | 0 | 0 | 0 | 4.06 | 95.94 | l. increase |

**Table S10.** Summary table of the EDF categorizations for the mean value across populations ("Mean spp"– see method) thermal performance curve parameters, for the three resources. In the "Data" column we have the parameters; in the "Experiment" column we have the resource type; in "Species" we have "mean_spp"; in "Model" we have whether a linear model (LM) or a generalized additive model (GAM) fitted the data best; in "Category" this correspond to the shape attributed to the curves fitted to the mean values across populations, we can find the following option: U-shape, hump-shape, monotonic or linear increase or decrease; the "Critical point" is the inflexion temperature, which is for a U-shape or hump-shape at which temperature the minimum or maximum was respectively reached; "Slope" if the mean fit curve was determined to be best fitted with a LM; "p-value" is the value from the fitted curve to assess if it is significant; finally "Sig. p-value" is the significance of the fitted curve in "yes"/"no" statement regardless of the value.

| **Data** | **Experiment** | **Species** | **Model** | **Category** | **Critical point** | **Slope** | **p_value_sig** | **P_value_sig** |
|---|---|---|---|---|---|---|---|---|
| Topt | Light | mean_spp | LM | l. increase | NA | 0.0087 | 0.0357781 | yes |
| Topt | Nitrogen | mean_spp | LM | l. decrease | NA | -1.00E-04 | 0.95857222 | no |
| Topt | Phosphorus | mean_spp | GAM | hump-shaped | 27.4 | NA | 0.39538588 | no |
| Tmax | Light | mean_spp | LM | l. increase | NA | 0.0025 | 0.13591331 | no |
| Tmax | Nitrogen | mean_spp | LM | l. increase | NA | 5.00E-04 | 0.23359061 | no |
| Tmax | Phosphorus | mean_spp | LM | l. increase | NA | 0.0181 | 0.03717541 | yes |
| Tbr | Light | mean_spp | GAM | U-shaped | 169.6 | NA | 0.0048305 | yes |
| Tbr | Nitrogen | mean_spp | LM | l. increase | NA | 0.0012 | 0.46555138 | no |
| Tbr | Phosphorus | mean_spp | LM | l. increase | NA | 0.0127 | 0.70697078 | no |

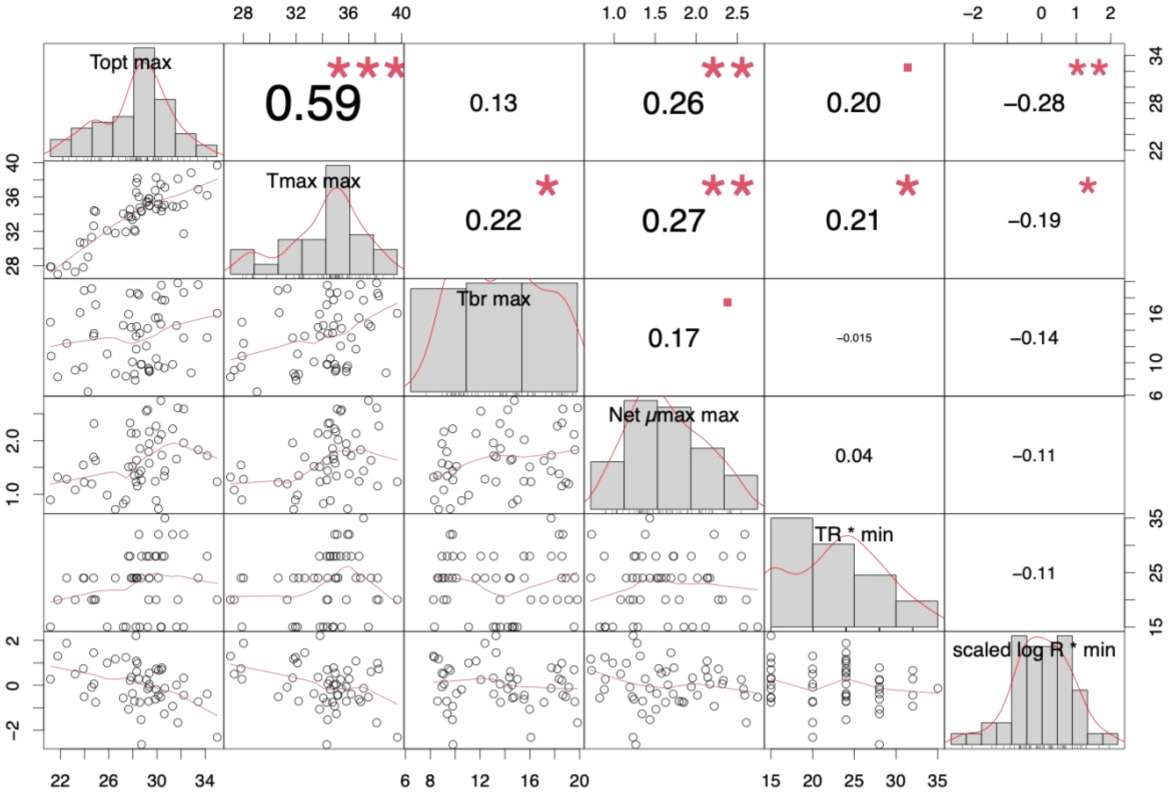

**Figure S19.** The correlation matrix of the summary variables from the PCA in Fig. 4 (main text). We can observe the correlation coefficient of the variables that cluster together in Fig. 4 A&B. The different levels of significance of the correlations are: "." p-value = 0.05; "*" p-value < 0.05; "**" p-value < 0.01; and "***" p-value < 0.001. There are 15 correlation pairs among our 6 summary variables, one pair displays a strong positive relationship, net-$\mu max_{max}$ -$TR*_{min}$ (0.59, p-value < 0.001), among the three pairs that display a significance of p-value < 0.01, two are positive $TR*_{min}$ - $Tmax_{max}$, and net-$\mu max_{max}$ - $Tmax_{max}$, but one is negative net-$\mu max_{max}$ - $scaled\ logR*_{min}$.

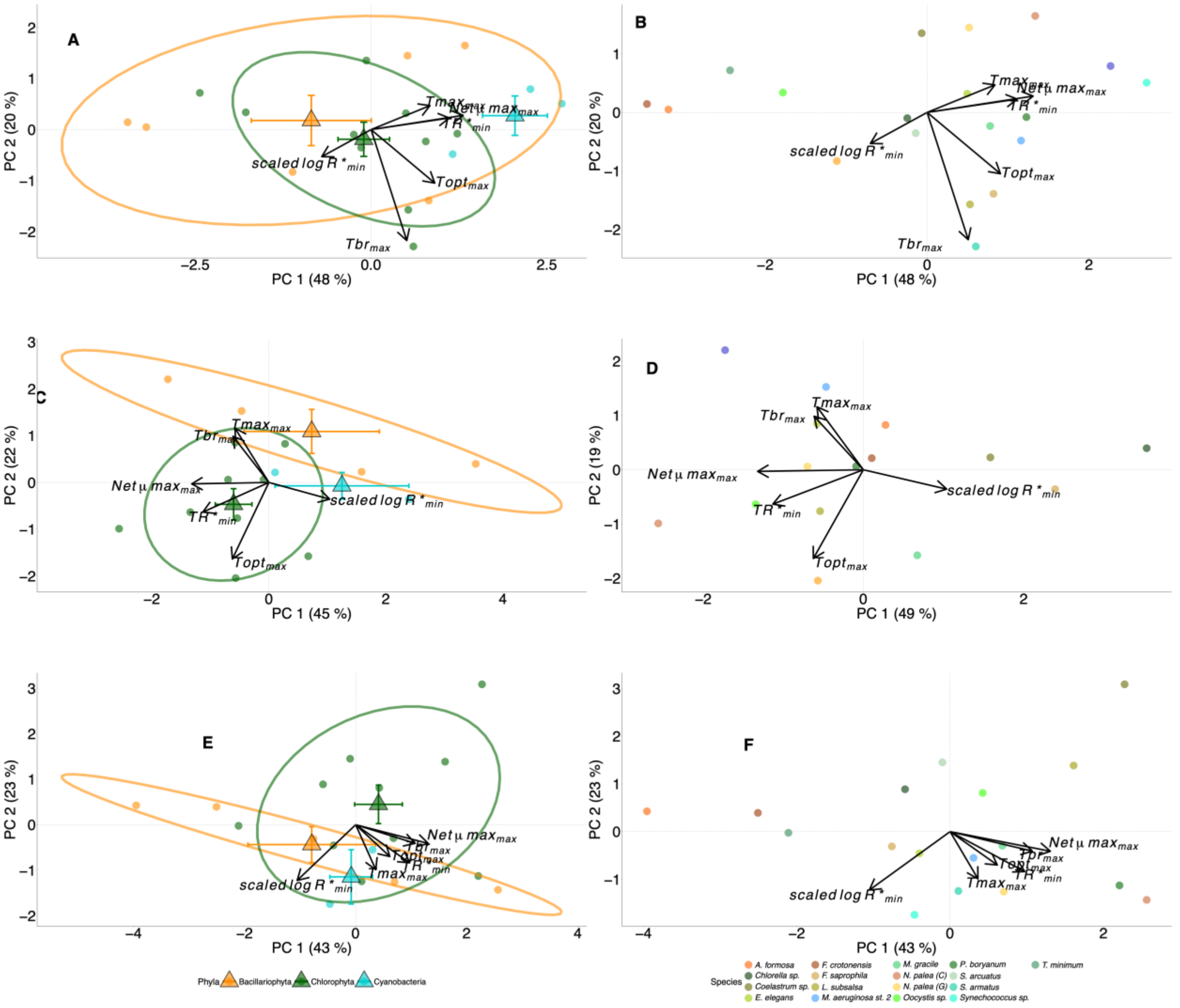

**Figure S20.** Principal components analysis (PCA) of the summary variables (minimum and maximum) of various Monod and TPC parameters for all three resources. The summary variables are calculated across temperature levels for each resource experiment and across resource levels for TPCs fitted for each resource. Parameters included in the PCAs are: the scaled log-transformed lowest minimum resource requirement across temperature for each resource (*scaled* $logR*_{min}$), the temperature at which this minimum $R*$ occurred ($TR*_{min}$), the greatest maximum growth rate ($\mu max_{max}$), the greatest thermal optimum ($Topt_{max}$), the greatest maximum temperature ($Tmax_{max}$), and the maximum thermal breadth ($Tbr_{max}$). Centroids are indicated with large triangular symbols (+/- standard error bars for PC1 and PC2) and ellipses represent one standard error of the means of the centroids (68% confidence ellipses). The

PCAs were run per resource type, so panels A&B are for light, C&D are for nitrogen and E&F are for phosphorus. The panels A, C & E are colored by phyla, but the B, D & F are by population name.

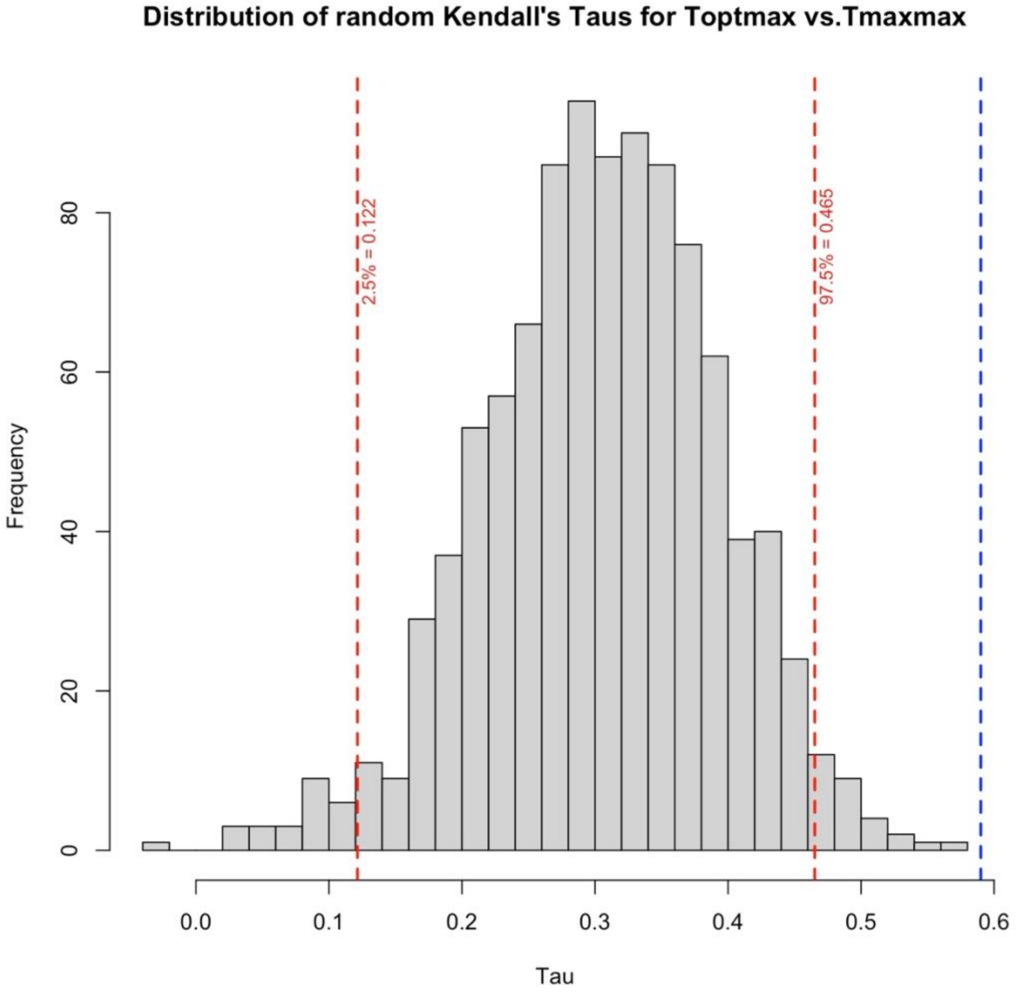

**<u>Figure S21.</u>** Random distribution of Kendall's Tau (**τ**) correlations between $Topt_{max}$ and $Tmax_{max}$. We generated this distribution to investigate the expected association between $Tmax_{max}$ and any other thermal parameter (i.e. temperature) that must be constrained by *Tmax* because, by definition species do not grow above *Tmax*. To generate this distribution of correlations of $Topt_{max}$ with $Tmax_{max}$, we first randomly selected a set of 56 $Tmax_{max}$ values that vary between 27 and 40°C, as they do in our data (Fig. 5A). We then also randomly selected 56 associated random $Topt_{max}$ values that vary between the random $Tmax_{max}$ selected for a single draw and the minimum value of $Topt_{max}$ observed in our dataset. We then calculated the Kendall's Tau correlation between $Topt_{max}$ and $Tmax_{max}$. We repeated this 1,000 times to generate the random distribution in the figure. The blue dashed line indicates the observed Kendall's Tau in our dataset and the red dashed vertical lines indicate the upper and lower 2.5 percentiles.

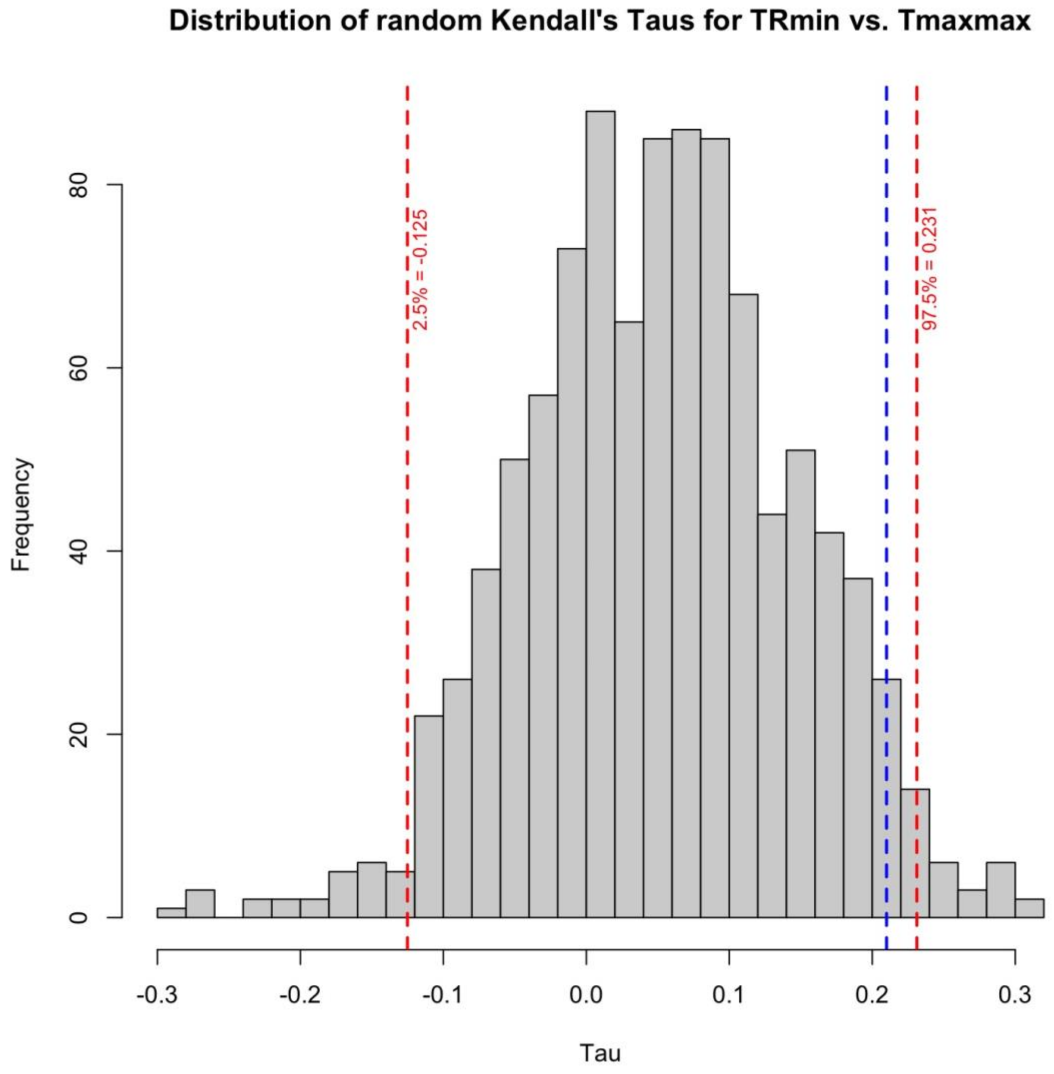

**<u>Figure S22.</u>** Random distribution of Kendall's Tau (**τ**) correlations between $TR^*_{min}$ and $Tmax_{max}$. The purpose and procedure are identical to that for Fig. S21, with the following exceptions. We randomly sampled 52 values of *TR*min* and $Tmax_{max}$ to match the number of data points in Fig. 5D. $TR^*_{min}$ values were randomly sampled but rounded to the nearest integer value of experimental temperatures at which *R**s could have been minimized in our experimental framework (i.e. 15, 20, 24, 28 32 or 35°C, as in Panel 5D). $TR^*_{min}$ random samples were drawn from values between the observed minimum in our dataset (15°C) and the randomly sampled $Tmax_{max}$ value (up to 40°C), but values of $TR^*_{min}$ values > 35 °C and their matching $Tmax_{max}$ values were filtered out of the simulated dataset to match the constraints of our actual dataset.
